# Biochemical and Immunological Properties of Engineered Low-Immunogenic Staphylokinases for Next-Generation Thrombolytic Therapy

**DOI:** 10.64898/2026.01.04.697554

**Authors:** Alan Strunga, Michaela Peskova, Michal Krupka, Katerina Zachova, Lenka Sindlerova, Nikola Neumeisterova, Jan Mican, Jan Vitecek, David Bednar, Stanislav Mazurenko, Lukas Kubala, Milan Raska, Jiri Damborsky, Zbynek Prokop

## Abstract

Staphylokinase (SAK) is a highly fibrin-specific plasminogen activator with significant potential as a safe and affordable thrombolytic. Yet, its clinical translation can be limited by potential immunogenicity. To accelerate the development of improved thrombolytics, a critical step is identifying the most suitable molecular template. Therefore, we performed a comparative analysis of biochemical and immunological properties of three engineered low-immunogenic variants (SAK SY155, SAK THR174, and SAK STAR FRIDA) and two wild-types (SAK STAR and SAK 42D), using a newly established panel of assays. All variants retained potent thrombolytic activity, with SAK SY155 displaying the highest catalytic efficiency and fibrin-clot permeability. However, this advantage did not fully translate into improved clot reduction under flow conditions. Comprehensive immunological profiling, including T lymphocyte proliferation, dendritic cell maturation, mouse immunization models, and human serum reactivity tests, confirmed decreased immunogenicity for two low-immunogenic variants. Overall, low-immunogenic SAK SY155 emerged as the most promising template for rational engineering of next-generation thrombolytics.

## Introduction

Stroke is a leading cause of death and long-term disability worldwide, placing a major burden on public health. According to the World Health Organization (WHO), it is the second most common cause of global mortality and a major contributor to disability-adjusted life years lost each year ^1^. Most cases (80-85%) are ischemic, caused by obstruction of cerebral blood flow ^2,3^. The current standard treatment is intravenous thrombolysis with alteplase, which improves outcomes when given within 4.5 hours ^4^, but its use is limited by the narrow therapeutic window and the risk of intracranial haemorrhage ^5^. After nearly thirty years, an engineered variant with greater fibrin specificity, a longer half-life, and single-bolus administration, called tenecteplase, was approved as an alternative ^6–8^.

Despite advances, the demand for safer and more effective thrombolytics continues to grow. Staphylokinase (SAK), a bacterial plasminogen activator, is a promising candidate. Unlike alteplase, it acts only in complex with plasmin and selectively activates fibrin-bound plasminogen, enabling targeted clot dissolution while minimizing systemic fibrinogen degradation and bleeding ^9–13^. Preclinical studies and early clinical data indicate strong efficacy and safety, suggesting that SAK may overcome several limitations of existing therapies ^14–19^. However, its bacterial origin makes it immunogenic, and antibody formation has been reported, potentially limiting repeated or long-term use ^20–22^.

Multiple less-immunogenic SAK variants have been engineered to address this challenge. Extensive work by Collen and colleagues (1995-2005) mapped immunogenic epitopes and generated numerous mutants with reduced immunogenicity and preserved activity ^23–28^. In this study, we focus on several of the most promising variants, compare their functional properties relevant to future applications, and evaluate their potential as templates for advanced engineering of safer thrombolytics. We selected three variants: (i) SAK SY155, a 12-point mutant (K35A, E65Q, K74R, D82A, S84A, T90A, E99D, T101S, E108A, K109A, K130T, K135R) identified as one of the most effective ^29^; (ii) SAK THR174, an 8-point mutant (E65D, K74Q, R77E, E80S, D82S, V112T, K130Y, E134R) previously advanced toward commercialization ^30^; and (iii) SAK STAR FRIDA, a simpler triple-alanine mutant (K74A, E75A, R77A) related to a variant clinically used in Russia ^14^.

Because staphylokinase exists in several naturally occurring forms, we also compared two wild-type backgrounds. SAK STAR is widely regarded as the default sequence from *Staphylococcus aureus*, while alternative phage-derived sequences, SAK 42D (S34G, G36R, H43R) and SAK ΦC (S34G), differ at a few positions ^31^. To capture functional diversity, we selected the more distinct SAK 42D variant and compared its performance with SAK STAR, ensuring consistent evaluation across engineered and wild-type forms.

## Material and Methods

### Construction, production, and purification of low-immunogenic SAK mutants

Previous studies show that both terminal regions of staphylokinase are critical for its function ^32–36^. Therefore, we designed a construct in which a tag can be fully removed, leaving no additional residues after purification. For this purpose, a HIS-tag was fused to a TEV protease recognition site (ENLYFQ/S(A,G)), enabling complete removal of the added sequence (highlighted in red in the Supplementary Data). TEV protease was selected as the cleavage enzyme because the native protein begins with a serine residue, for which TEV exhibits particularly high cleavage efficiency.

Recombinant staphylokinase (SAK) variants were expressed in *Escherichia coli (E. coli)* BL21(DE3) harboring pET21b-based constructs. Constructs for all variants (SAK STAR WT, SAK 42D WT, SAK STAR FRIDA, SAK THR174 and SAK SY155) were obtained from gene synthesis. Cultures were grown in 2× Lysogeny broth (LB - 20 g/L tryptone, 10 g/L yeast extract, 10 g/L NaCl; pH 7.0) supplemented with ampicillin (100 µg/mL) at 37 °C to an OD₆₀₀ of ∼0.8, followed by induction with 0.5 mM IPTG for 4 h at 37 °C. Cells were harvested (4000 ×g, 30 min, 4 °C), resuspended in binding buffer A (20 mM K₂HPO₄, 500 mM NaCl, 10 mM imidazole, pH 7.5), and stored at −70 °C.

Cell lysis was performed by sonication on ice, and lysates were clarified by centrifugation (19,000 ×g, 1 h, 4 °C) and 0.45 µm filtration. The supernatant was loaded onto a Ni²⁺-IMAC column (HisTrap™ Excel) using an ÄKTA Pure system (Cytiva, Germany). After washing with binding buffer A, SAK proteins were eluted with elution buffer B (20 mM K₂HPO₄, 500 mM NaCl, 500 mM imidazole; pH 7.5). His-tag removal was carried out by overnight TEV protease digestion during dialysis into phospate-buffered saline (PBS, 10 mM Na₂HPO₄, 1.8 mM KH₂PO₄, 2.7 mM KCl, 137 mM NaCl; pH 7.4), using 1 mg TEV per 100 mg SAK. A second IMAC step was used to collect tag-free SAK in the flow-through. Final purification was achieved by size-exclusion chromatography (HiLoad 26/600 Superdex 75 pg) in PBS. Pure fractions were pooled, concentrated, flash-frozen in liquid nitrogen, and stored at −70 °C. Protein concentration and purity were assessed by UV spectrophotometry and SDS–PAGE.

### Endotoxin removal procedure

Endotoxins were removed from protein samples using Triton X-114 phase separation ^37,38^. Triton X-114 was added to the protein solution to a final concentration of 1% (v/v), and the mixture was incubated on ice for 20 min until clarification. Samples were then warmed to 25–30 °C to induce phase separation and centrifuged at 5000 rpm for 10 min at 25 °C. The aqueous phase containing proteins was collected, while the detergent phase containing endotoxins was discarded. This procedure was repeated three times to maximize endotoxin removal. All steps were performed using sterile, pyrogen-free plasticware.

### Thermostability measurement

Thermostability of SAK variants was evaluated by differential scanning fluorimetry (DSF) using a Prometheus Panta instrument (Nanotemper Technologies, Germany). Protein samples (∼1 mg/mL; ≈65 µM) were loaded into standard glass capillaries in quadruplicate. Thermal scans were recorded in the range of 25–90 °C at a 1 °C/min rate, monitoring intrinsic tryptophan fluorescence at 330 and 350 nm. Unfolding transitions were derived from the 350/330 nm fluorescence ratio, and melting (T_m_) and onset (T_on_) temperatures were determined using Prometheus Panta Evaluation Software. Variants were classified as stable when both T_m_ and T_on_ exceeded 37 °C. Long-term stability was assessed by an isothermal DSF/DLS measurement at 37 °C for 20 h using the same protein concentration.

### Fluorescence activity measurement

Unless otherwise stated, reactions were performed in PBS. A combination of a fluorogenic assay, an absobance assay, and surface plasmon resonance (SPR) measurement was selected for thrombolytic activity characterisation ^39,40^.

A calibration curve was constructed using varying 7-amino-4-methylcoumarin (AMC) concentrations (0, 10, 20, 50, 100, 200, 400, and 600 µM) (Merck, Germany). Fluorescence was monitored continuously for 4 h to ensure signal stability, and a mean value was taken for each concentration. Data were fitted to the conversion Eq. 1.

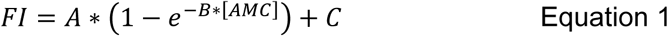

where FI is the fluorescence intensity, [AMC] is the AMC concentration, and A, B, and C are fitting parameters. The obtained function was used to convert fluorescence intensities to AMC concentrations in all subsequent assays.

To characterize human plasmin kinetics, reactions were performed with plasmin (32 nM) (Athens R&T, USA) and varying D-Valyl-Leucyl-Lysyl-7-amido-4-methylcoumarin (VLK-AMC) concentrations (0, 10, 20, 50, 100, 200, 400, and 600 µM) (AAT Bioquest, USA). Fluorescence data were converted to AMC concentrations using the calibration curve. Linear slopes at each substrate concentration were extracted and fitted to the Michaelis–Menten equation to obtain the maximum reaction velocity (V_max_) and Michaelis constant (K_M_). The turnover number (k_cat_) and catalytic efficiency (k_cat_/K_M_) of plasmin were then calculated from these parameters.

SAK activity was measured using human plasminogen (PLG, Athens R&T, USA) as substrate in the presence of VLK-AMC (200 µM). Plasminogen concentrations were 0, 0.16, 0.81, 2.17, 4.35, 8.15, 16.3, and 43.5 µM, while SAK concentration was fixed at 0.5 nM. The mixture was spiked with 0.5 nM of human plasmin (PLM, Athens R&T, USA) to allow the creation of SAK-PLM complex. The fluorescence was monitored at λ_ex_ = 355-20 nm and λ_em_ = 460-10 nm using an Omega plate reader (BMG Labtech, Germany). Fluorescence progress curves displayed four characteristic phases: lag, parabolic, linear, and plateau. Only the initial parabolic phase, where both plasminogen activation and VLK-AMC cleavage occur simultaneously, was used for kinetic evaluation. This phase was fitted with a second-degree polynomial, and quadratic coefficients were taken as apparent reaction velocities. These values were plotted against plasminogen concentration and fitted by the Michaelis–Menten equation to obtain apparent kinetic parameters (V_max_app_ and K_M_app_).The turnover number of the SAK–PLM complex was calculated using Eq. 2, which accounts for plasmin kinetics determined independently:

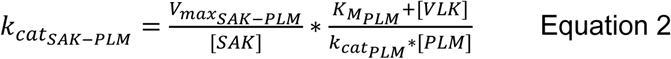

Uncertainty in k_cat,SAK–PLM_ was estimated by error propagation using the standard deviations of V_max,SAK–PLM_, K_M,PLM_, and k_cat,PLM_.

### Absorbance thrombolytic efficiency measurement

The thrombolytic efficiency of SAK variants was assessed using a simplified version of a previously published Clot Formation and Lysis (CloFAL) assay ^40^. For each reaction, 50 µL of thrombin (1 IU/mL) (T6884, Merck, Germany) was premixed with 5 µL of SAK variant (50 nM). Clotting was initiated by adding 50 µL of fibrinogen (10 mg/mL) (F3879, Merck, Germany), which contains traces of plasminogen, and the reaction was immediately monitored in a plate reader by measuring absorbance at 405 nm at 30 s intervals. Reactions were carried out in PBS supplemented with 0.9 mM CaCl₂ and 0.1% Tween-80. Calcium was included to support clot formation, while Tween-80 prevented clot adherence to the well walls and improved homogeneity of the mixture.

Data analysis was performed using the Shiny App ^41^. For each variant, two parameters were determined: the time to 90% clot formation and the time to 50% clot lysis. Thrombolytic efficiency was reported as the difference between these two values, representing the time required for a fully formed clot to undergo 50% lysis.

### Surface plasmon resonance (SPR) measurement

To investigate whether differences in activity among SAK variants could be attributed to altered affinity for plasmin, binding kinetics were analyzed by surface plasmon resonance (SPR) using a Biacore S200 instrument (Cytiva, USA). Human plasmin (Invitrogen, USA) was immobilized on a CM5 sensor chip via amine coupling. SAK variants were injected in the mobile phase at concentrations of 3.7, 11, 33, 100, 300, 900 nM at a flow rate of 50 µL/min. For each variant, three measurements were performed on the same chip, with alternating injections to minimize systematic bias. The samples ran across all four channels. Channels 1 and 3 served as blank channels, while channels 2 and 4 had two different coverages: 850 units on channel 2 and 550 units on channel 4. After subtraction, we obtained two independent traces (2-1 and 4-3). The association and dissociation phases were both set to 300 s. Regeneration was achieved using 10 mM Glycine/HCl, pH = 2.5, for 30 s. The measurement was performed at 25 °C.

All sensorgrams collected for a given SAK variant were analyzed globally using Kintek software (Kintek Corporation, USA). Data fitting was performed by numerically integrating of the differential rate equations defined by the kinetic model (Scheme 1), and the resulting simulated sensorgrams were fitted to the experimental data. The best-fitting model suggests two parallel binding events, a two-step induced binding and an independent one-step binding. Parameter estimation was performed by minimizing the χ² objective function using nonlinear least-squares regression based on the Levenberg–Marquardt algorithm. The forward integration of the model equations was computed using the Bulirsch–Stoer method with adaptive step-size control to ensure numerical accuracy and stability during optimization ^42^. Residuals were normalized by the sigma value for each data point. The observable sensorgram signal was defined as the sum of the contributions of each species: Signal = a*(PLM-SAK_A1_ + b*PLM-SAK_A2_ + c*PLM-SAK_B_) + d*SAK, where a scales the signal to the sensitivity of the measurement, factors b and c define the relative change in signal in forming PLM-SAK_A1_, PLM-SAK_A2_, and PLM-SAK_B_ complexes, respectively. The factor d defines the signal contribution of the non-specific binding of SAK. Different fluorescence scaling factors relating to the different sensitivity of the measurements were used for different datasets, but the factors defining the relative change in signal in forming individual complexes was constant globally for all datasets. The scaling factors were used as fitted parameters. The standard error of the obtained parameters was calculated from the covariance matrix during nonlinear regression. The standard error estimates in fitted parameters were propagated to yield error estimates in calculated values, the equilibrium dissociation constants of PLM-SAK_A1_, PLM-SAK_A2_, and PLM-SAK_B_ complexes. In addition to standard error values, a more rigorous analysis of the variation of the kinetic parameters was performed using confidence contour analysis with FitSpace Explorer (KinTek, USA) ^43^. In this analysis, the lower and upper limits for each parameter were derived from the confidence contours for the χ2 threshold at the boundary of 0.98.

**Scheme 1.**
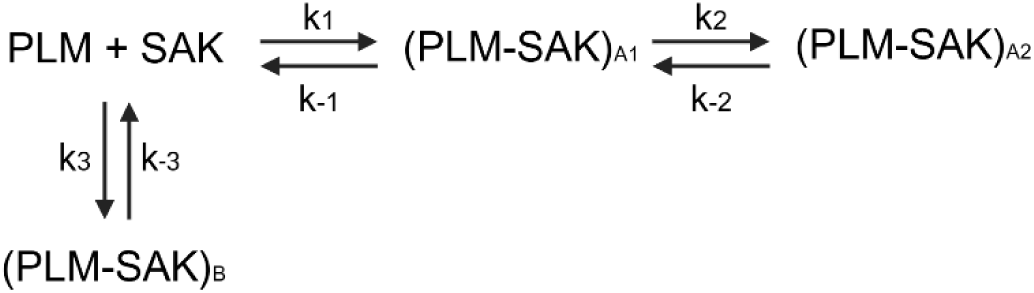

### Blood collection and processing

Venous blood was collected from healthy adult donors (both males and females) after obtaining written informed consent. Donors who received acetylsalicylic acid, nonsteroidal anti-inflammatory drugs, or antiplatelet agents within 7 days were excluded. The study protocol was approved by the Ethics Committee of the St. Anne’s University Hospital Brno (EK-FNUSA-25/2023).

Red blood cells (RBCs) were isolated from whole blood that had been anticoagulated with 3.8% sodium citrate (at a ratio of 1:9) through three centrifugation cycles (300×g for 10 min at 4°C), with two intermediate PBS washes. The final wash used a saline-adenine-glucose-mannitol solution (1.3 mM adenine, 45.4 mM glucose, and 28.8 mM mannitol in PBS, pH 7.4). Isolated RBCs were stored at 4°C for up to 21 days and centrifuged (150×g for 10 min at 4°C) before use ^44^. Serum was obtained from coagulated blood accelerated by thrombin (5 IU/mL) and CaCl₂ (14 mM), and then centrifuged (1000×g for 10 min at 4°C), aliquoted, and stored at −80°C.

### Clot and fibrin gel preparation

Semi-synthetic clots were formed using human fibrinogen (12.1 mg/mL; Tisseel Kit Lyo, Baxter) and human thrombin (0.77 IU/mL; Tisseel Kit Lyo, Baxter), combined with 4.2×109/mL isolated RBCs, 15-fold diluted serum, and 4.6 mM CaCl₂. Clot formation was performed in 0.5- or 0.2-mL polypropylene tubes for 45 min at 37 °C with >90% relative humidity. The total clot volumes were 90 µL for the static model and 25 µL for the flow model.

For the penetration experiments, fibrin gels were prepared according to a previously published protocol ^39^ with the following modifications: the final concentrations were as follows: fibrinogen (12.1 mg/mL), thrombin (0.77 IU/mL), and CaCl₂ (6.9 mM). A 30 µL aliquot of the mixture was cast directly into µ-Slide VI 0.4 micro-slide channels (Ibidi, Germany) and allowed to clot under the conditions described above.

### Thrombolytic agents in vessel models

To evaluate thrombolytic efficacy, plasminogen activators were tested at a concentration of 1.3 µg/mL, which corresponds to the clinically relevant dose of alteplase (ALT; Actilyse®, Boehringer Ingelheim, Germany). This concentration was derived from pharmacokinetic data ^45^ and the manufacturer’s information. For the penetration assay, the agents were fluorescently labelled with rhodamine B isothiocyanate (HY-112697, MedChemExpress, USA) according to the manufacturer’s protocol ^39^. Stabilizing additives were removed from ALT using PD MidiTrap G-25 desalting columns (Cytiva, USA) according to the manufacturer’s instructions, enabling buffer exchange with PBS prior to fluorescence labelling. Betaine hydrochloride (90 mM per mg of protein) was added to the ALT sample to prevent denaturation during labelling. After labelling, the fluorophore-to-protein ratio was set to 0.39 for all thrombolytic agents. The final concentration in the penetration experiments was set to 0.022 µM, which is equivalent to the clinical concentration of ALT.

### Determination of thrombolytic efficacy in a static model

Individual clots were incubated in 1.5 mL polypropylene tubes containing 0.5 mL of human plasma (Human Plasma – Pooled, Seqens In Vitro Diagnostic, France) at 37 °C for 60 min, which corresponds to the standard ALT infusion duration used in clinical stroke treatment. Lytic efficacy was assessed by measuring the reduction in clot weight and by spectrophotometrically quantifying the release of RBC at 575 nm (hemoglobin) with 700 nm turbidity correction ^46,47^.

### Determination of thrombolytic efficacy in a flow model

An *in vitro* middle cerebral artery (MCA) flow model was constructed using silicone chips (Sylgard 184, Dow Corning, USA), designed based on the dimensions of patient CT scans (n=4). The models incorporated anatomical narrowing and bifurcation to enable vessel occlusion and permanent circulation. Each MCA chip was connected to a peristaltic pump (Gilson Minipuls 3, Gilson, USA) to maintain plasma flow for up to 120 min at 37 °C or until complete recanalization (clot displacement and restored flow) occurred ^48^.

Real-time imaging captured clot area changes at 5-minute intervals using a web camera (C525, Logitech, Switzerland) with a resolution of 2 megapixels. Clot lysis was quantified by percentage area reduction, recanalization time, and RBC release. Recanalization frequency represented the percentage of complete recanalization per treatment group.

### Clot penetration assay

Fluorescently labelled thrombolytics were applied to individual fibrin gel channels, and the fluorescence intensity was recorded at 5-minute intervals at distances of 0.25, 0.50, 0.75, 1.00, and 1.25 mm from the application site. Imaging was performed using a Zeiss Axio Observer Z1 microscope (Zeiss, Germany) with metal halide fluorescence illumination (HXP 120, LEJ, Germany) and a high-efficiency Filter Set (Filter Set 43 HE: excitation BP 550/25, beam splitter FT 570, and emission BP 605/70). Penetration success was defined as the percentage of runs exceeding three times the standard deviation of background autofluorescence within 180 min.

### Image analysis and data processing

Clot area measurements were performed using semi-automated procedures with Python (version 3.13) and ImageJ (version 1.54d). Penetration analysis utilized custom Python scripts with background subtraction and winsorization at the 5th and 95th percentiles to reduce the influence of outliers ^49,50^, and normalization to the baseline (time 0 min).

### Statistics for vessel model experiments

Statistical analyses were performed using GraphPad Prism (version 9.5.1, GraphPad Software, USA) and Python. Outliers were identified using Grubb’s test, and normality was assessed using the Shapiro–Wilk test. Multiple group comparisons were performed using one-way ANOVA with Tukey’s correction or the Kruskal-Wallis test with Dunn’s correction. Recanalization and penetration success frequencies were analyzed using Fisher’s exact test with Holm–Bonferroni correction. The correlation between the penetration rate and distance was evaluated using Spearman’s correlation coefficient. Statistical significance was set at p<0.05. Measurements were performed in 2-6 technical replicates.

### Monocyte activation test (MAT) and T lymphocyte proliferation

Low-binding Eppendorf tubes were filled with 800 µL 0.9% NaCl, 100 µL heparinized human whole blood (40 IU/mL), and 100 µL of either NaCl (negative control), lipopolysaccharide (LPS; E. coli O5:B55; 0.007 EU/mL representing the acceptable endotoxin threshold for intravenously administered drugs as defined by the European Pharmacopoeia ^51^, and 0.07 EU/mL corresponding to a 10-fold higher concentration), ALT (1.3 µg/mL, clinically relevant, and 13 µg/mL), or the SAK variants (1.3 and 13 µg/mL). Samples were incubated for 24 h at 37°C in a humidified 5% CO₂ incubator under sterile conditions. Following incubation, samples were centrifuged at 500 × g for 10 min at room temperature (RT), and plasma was carefully collected without disturbing the cell pellet. IL-6 concentrations were quantified by ELISA (Human IL-6 ELISA kit, Invitrogen, USA) according to the manufacturer’s protocol.

Peripheral blood was collected from healthy donors into 4 mM EDTA as an anticoagulant. Peripheral blood mononuclear cells (PBMCs) were isolated, and CD8⁺ T lymphocytes were depleted using the RosetteSep™ Human CD8 Depletion Cocktail (STEMCELL Technologies, Canada) according to the manufacturer’s protocol. CD8-depleted PBMC population was labeled immediately after isolation with CellTrace™ Violet (Thermo Fisher Scientific, USA) to determine cell proliferation, resuspended in RPMI 1640 culture medium (Thermo Fisher Scientific, USA) supplemented with 10% low endotoxin fetal bovine serum (FBS LE; Diagnovum, Germany) and antibiotics (100 U/mL penicillin and 100 µg/mL streptomycin, Thermo Fisher Scientific, USA), and seeded in 24-well plates at a density of 5 × 105 cells per well in 500 µL. Cultures were maintained overnight at 37 °C in a humidified 5% CO₂ incubator, under sterile conditions.

On the following day, cells were stimulated with either medium alone (negative control), phytohemagglutinin (PHA, 5 µg/mL; positive control), ALT (1.3 µg/mL, clinically relevant and 13 µg/mL), or SAK variants (1.3 and 13 µg/mL). Cells were incubated for 5 days at 37 °C in a humidified 5% CO₂ incubator under sterile conditions. Lymphocyte proliferation was assessed by flow cytometry (BD FACSVerse; BD Biosciences, USA) using anti-CD3 antibody (Caltag Laboratories, USA) and 7-aminoactinomycin D (7-AAD; Cayman Chemical Company, USA) staining to identify viable, proliferating T cells.

Statistical analyses were performed using GraphPad Prism (version 9.5.1, GraphPad Software, USA). For MAT data, outliers were identified using Grubb’s test, and normality was assessed using the Shapiro-Wilk test. Multiple group comparisons were performed using the Kruskal-Wallis test with Dunn’s post hoc correction, with all test conditions compared to the response at the negative control and 0.007 EU/mL LPS. For T lymphocyte proliferation data, the variance homogeneity was tested using the Brown-Forsythe test. Multiple group comparisons were performed using ordinary one-way ANOVA with Tukey’s post hoc correction.

### Mouse immunization experiments

All experiments were performed on 6- to 8-week-old female BALB/c mice (Anlab, Czech Republic). All animals were free of known pathogens and were kept in a climate-controlled environment and were provided with pellet food and water ad libitum. Each of the mice was immunized three times intradermally with 100 µL of SAK formulation (containing 5 µg of SAK protein) with a three-week interval between doses. For each SAK, one group of mice was immunized with a SAK protein without adjuvant and one with an aluminum adjuvant, as reported earlier ^52^. Each group of mice consisted of 3 individuals. Two weeks after the last dose, mice were euthanized under Ketamine/Xylazine anesthesia ^53^ and blood samples were taken from which serum was isolated by centrifugation.

### Determination of autoantibodies in the mouse sera

An immunoblotting method was used to determine the potential cross-reactivity of SAK antigens with human autoantigens. We used diagnostic kits EUROLINE Autoimmune Inflammatory Myopathies 16 Ag (IgG) containing the following antigens: Mi-2a, Mi-2b, TIF1g, MDA5, NXP2, SAE1, Ku, PM-Scl100, PM-Scl75, Jo-1, SRP, PL-7, PL-12, EJ, OJ, Ro-52, EUROLINE ANA profile 3 plus DFS70 (IgG) kit containing antigens nRNP/Sm, Sm, SS-A, SS-B, Scl-70, Jo-1, dsDNA, nucleosomes, histones, ribosomal P-protein, AMA M2, Ro-52, PM-Scl, CENP B, PCNA, and DFS70 and EUROLINE Anti-MPO, -PR3 and -GBM (IgG) (all EUROIMMUN, Germany). As Jo-1 and Ro-52 antigens are included in two sets, a result was obtained for a total of 33 autoantigens. For use with mouse sera, the secondary antibody contained in the kit was replaced by an anti-mouse IgG AP-conjugate antibody (A16093, Invitrogen, USA) as described previously ^54^.

### Determination of murine antibodies against individual SAKs by ELISA

Wells of ELISA plates (Multisorp, Thermo Fisher Scientific, USA) were coated with 100 μL of individual SAK solutions with a protein concentration of 5 μg/mL in carbonate buffer (100 mM bicarbonate/carbonate, pH 9.6) at 4 °C overnight. The plates were washed with PBS-T solution (PBS buffer containing 0.05% Tween, pH 7.4), and wells were blocked with blocking solution (5% dry skimmed milk in PBS-T). Mouse sera samples were applied at a 1:500 dilution in PBS-T supplemented with 1% bovine serum albumin (BSA, Merck, Germany). Samples were incubated at 4 °C overnight, wells were washed three times with PBS-T and secondary antibody (anti-mouse IgG γ-chain specific-peroxidase antibody produced in goat, Merck, Germany) was applied at a dilution of 1:5000 in PBS-T with 1% BSA. After two hours of incubation at RT, wells were washed 4× with PBS-T, 2× with PBS and OPD tablet (Merck, Germany) in Phosphate-Citrate buffer (Merck, Germany) was used to visualize the reaction. The reaction was stopped by 1 M sulphuric acid, and absorbance at 450 nm was measured using BioTek Synergy HTX multi-mode reader (Agilent Technologies, USA).

### Determination of murine antibodies against SAK by measuring inhibitory potency

The inhibitory potency of murine sera after administration of SAK was studied using the fluorogenic activity assay. 9 samples were measured for each variant – 3 negative control serum samples, 3 serum samples from adjuvant-free administration and 3 samples from adjuvant-rich administration. All serum samples were first incubated at 56 °C for 30 minutes to inactivate the complement. Then the sera were diluted 10-fold and incubated with SAK at 37 °C for one hour to create complexes between SAK molecules and newly formed antibodies in the sera. Preliminary experiments showed that the optimal concentration of SAK for incubation is 333 nM. The inhibitory potency of individual serum samples was compared by measuring the decrease in reaction rate. The reaction mixture contained 1 µM human PLG, 200 µM fluorogenic substrate, 13.3 nM human PLM, 13.3 nM SAK, and 250-times diluted serum. The decrease in the rates between negative control samples and samples of serum after administration of SAK was studied.

### Determination of human antibodies against individual SAKs by ELISA

Wells of ELISA plates (Multisorp, Thermo Fisher Scientific, USA) were coated with 100 μL of individual SAK solutions with a protein concentration of 5 μg/mL in carbonate buffer (100 mM bicarbonate/carbonate, pH 9.6) at 4 °C overnight. The plates were washed with PBS-T solution (PBS buffer containing 0.05% Tween, pH 7.4), and wells were blocked with blocking solution (5% dry skimmed milk in PBS-T). Mouse sera samples were applied at a 1:500 dilution in PBS-T supplemented with 1% bovine serum albumin (BSA, Merck, Germany). Samples were incubated at 4 °C overnight, wells were washed three times with PBS-T and secondary antibody (anti-mouse IgG γ-chain specific-peroxidase antibody produced in goat, Merck, Germany) was applied at a dilution of 1:5000 in PBS-T with 1% BSA. After two hours of incubation at RT, wells were washed 4× with PBS-T, 2× with PBS and OPD tablet (Merck, Germany) in Phosphate-Citrate buffer (Merck, Germany) was used to visualize the reaction. The reaction was stopped by 1 M sulphuric acid, and absorbance at 450 nm was measured using BioTek Synergy HTX multi-mode reader (Agilent Technologies, USA).

The principal component analysis was then performed on the centered non-normalized values of absorbances, averaged over two independent experiments, using sklearn.decomposition.PCA function from scikit-learn library (v1.18.0) in Python 3.14. The dendrogram with the SAK variants was calculated on the same values using cluster.hierarchy methods of SciPy library (v1.17.0) using default linkage settings.

### Bone marrow-derived dendritic cells differentiation

The preparation and differentiation of bone marrow-derived dendritic cells (BMDCs) from mice was performed with the aseptic harvesting of bone marrow from the femurs and tibias of BALB/c mice. All reagents used in this protocol were purchased from Gibco (Thermo Fisher Scientific, USA). After euthanizing the mouse under ketamine-xylazine anesthesia, the bones were cleaned of muscle tissue, sterilized with ethanol, and their ends were cut off to expose the marrow. Using a sterile syringe, the marrow was flushed out with complete RPMI culture medium (10% FBS, 1% penicillin/streptomycin) into a collection tube. Red blood cells were lysed, followed by centrifugation at 250 g for 5 min at RT. Bone marrow cells were incubated for 7 days in complete RPMI culture medium (10% FBS, 1% penicillin/streptomycin) supplemented with 20 ng/mL GM-CSF and 20 ng/mL IL-4 at 37°C, 5% CO_2_. On days 3 and 5, one-third of the media was replaced by fresh media with cytokines. On day 7, 50 µM of particular SAK variants, or 150 ng/ml LPS (Merck, Germany) was added to differentiated BMDC and cultivated for another 24 hours.

Cells were washed with PBS by centrifugation at 250 g for 5 min at RT, and Fc receptors were blocked by TruStain FcX antibody (SONY Biotechnology, USA). A set of cell surface markers anti-MHCII – Alexa Fluor700, anti-MHCI – Pacific Blue, anti-CD11c – APC-cy7, anti-CD40 – PE, anti-CD80 – PE-cy5, anti-CD86 – PE-cy7, anti-CD69 – APC (SONY), and Fixable Viability Dye – eFluor506 (e-Bioscience, Thermo Fisher Scientific) was used in 2% FBS in PBS for 30 min, RT, in the dark. Cells were washed in PBS and analysed by SONY SP6800 spectral analyser. Flow cytometry Fcs files were analysed in SONY software, more detailed analyses (t-SNE) were performed by FlowJo 10.6.1. software (Becton Dickinson, USA). All statistical analyses were performed in GraphPad Prism 8 software (GraphPad Software, USA).

## Results

### Protein purity, yield, stability, and aggregation propensity

The purity of all SAK variants (sequences shown in Table S11) was assessed by SDS–PAGE, using samples collected after each purification step and final protein preparations (Fig. S3 and S4). The purification strategy consistently yielded highly pure proteins across all variants; however, protein yields varied considerably between constructs (Table S11). SAK STAR WT was obtained in substantially higher amounts compared to SAK 42D WT. Incorporation of multiple mutations generally resulted in reduced expression yields; however, the high yield of STAR WT ensured sufficient quantities of even the lowest-yielding variants.

Stability analysis by differential scanning fluorimetry (DSF) (Fig. 1A, S5) revealed that reduced yields were accompanied by decreased thermostability. Among all constructs, SAK STAR WT again showed the most favourable profile, with both melting temperature (T_m_) and onset of unfolding temperature (T_on_) substantially above 37 °C. Importantly, all variants showed T_m_ and T_on_ values above the physiological threshold. To further confirm long-term stability at physiological conditions, a combined isothermal DSF/DLS experiment was performed at 37 °C for 20 h (Fig. S6 and S7). No signs of unfolding or aggregation were observed, demonstrating that all variants remain stable during storage, handling, and conditions relevant to administration.

**Figure 1.**
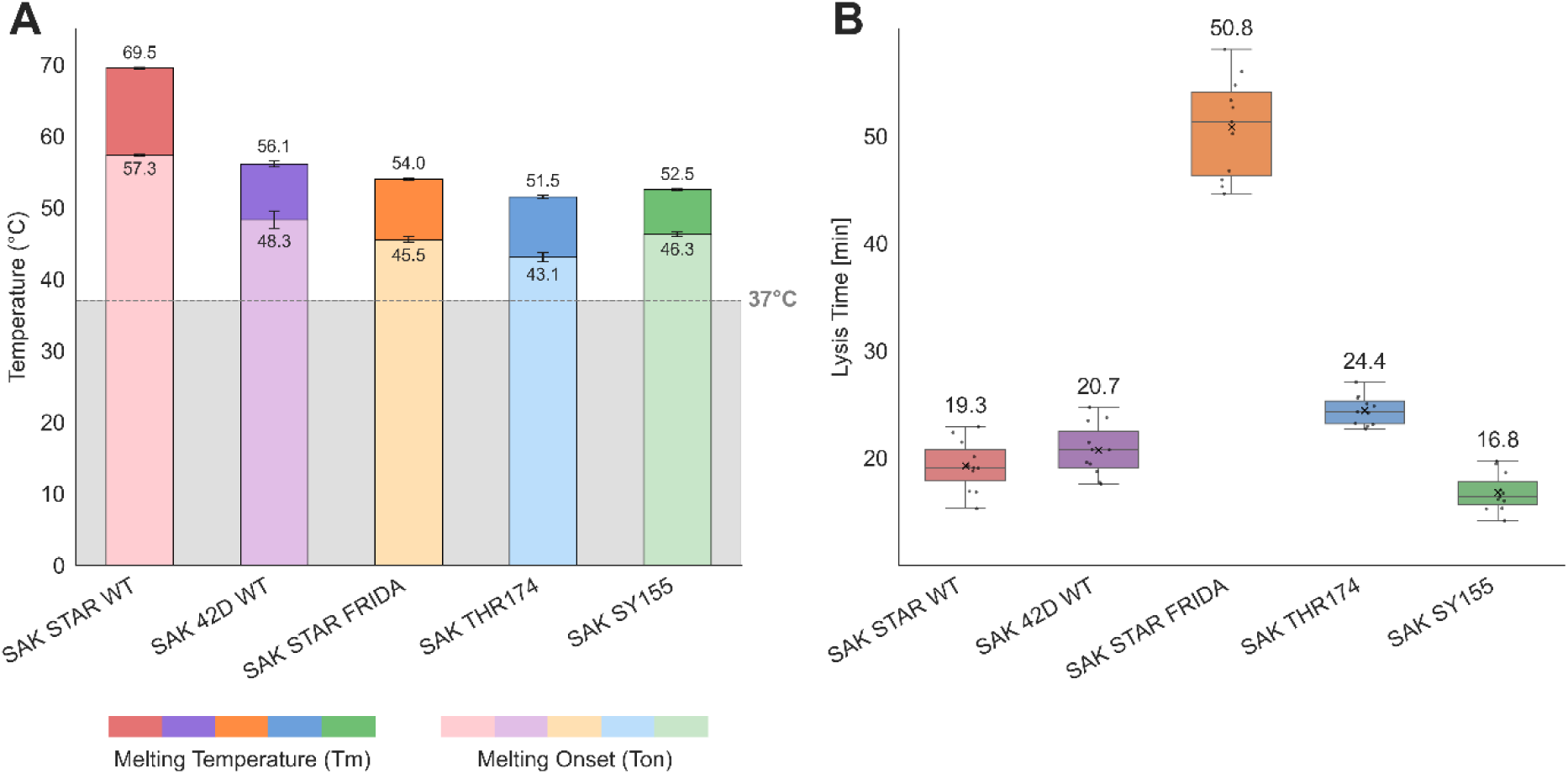
**A)** Differential scanning fluorimetry (DSF) analysis of the thermal stability of the five SAK variants. For each variant, the melting temperature (Tm) and onset melting temperature (Ton) are shown. All measurements were performed at ∼1 mg/mL protein concentration (≈65 µM) in quadruplicate. Error bars represent one standard deviation above and below the average. The data demonstrates that all variants maintain Tm and Ton values above physiological temperature (grey line = 37 °C). **B)** The results of the thrombolytic activity assay on fibrin clots, run at 37°C in PBS/Ca2+/Tween80; pH=7.4, number of repetitions was 11. The reported value is the time it takes for the variants to half-lyse a fibrin clot once the clot has been almost fully formed (90% clotting). The box plot shows mean value (X), median (line), interquartile ranges (boxes), and minimum and maximum values (whiskers).

### Comparison of catalytic activities using in vitro assays

To assess the performance and thrombolytic potential of the SAK variants, we conducted a series of kinetic analyses, affinity measurements, and activity profiling. We analyzed the steady-state kinetics of all staphylokinase variants using the fluorogenic assay (Fig. S8). The resulting kinetic parameters are summarized in Table 1. The variants exhibited measurable differences in the proteolytic activity. Notably, SAK STAR FRIDA exhibited a significantly elevated Michaelis constant, K_M_ >> 50 uM, so only the catalytic efficiency (k_cat_/K_M_) could be reliably estimated, using linear fitting with the Eq. 3.

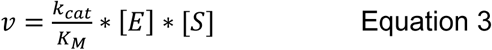

**Table 1.** Apparent kinetic parameters, maximum reaction velocity (V_max_), Michaelis constant (K_M_), turnover number (k_cat_), and catalytic efficiency (k_cat_/K_M_) were determined from the fluorogenic assay, run with human plasminogen (0 - 43.5 µM) in PBS buffer; pH=7.4; 37°C in triplicate. ** Due to significantly elevated K_M_ for SAK STAR FRIDA, Michaelis-Menten fit was replaced with simple linear fitting that only allows for assessing catalytic efficiency, but not the remaining parameters.

| Steady-state kinetics parameters |  |  |  |  |  |
| --- | --- | --- | --- | --- | --- |
|  | SAK STAR<br>WT | SAK 42D WT | SAK STAR<br>FRIDA | SAK THR174 | SAK SY155 |
| $V_{max\_app}$ μM·s$^{-1}$ ($\cdot 10^{-3}$) | 0.193 ± 0.009 | 0.134 ± 0.006 | ** | 0.194 ± 0.024 | 0.199 ± 0.11 |
| $K_M\_app$ [μM] | 6.75 ± 0.93 | 7.54 ± 0.98 | ** | 14.17 ± 4.06 | 4.78 ± 0.84 |
| $k_{cat\_app}$ [s$^{-1}$] | 1.88 ± 0.16 | 1.30 ± 0.11 | ** | 1.89 ± 0.27 | 1.94 ± 0.17 |
| $k_{cat}/K_M\_app$ [μM$^{-1}$ ·s$^{-1}$] | 0.278 ± 0.045 | 0.173 ± 0.027 | 0.016 ± 0.0003 | 0.133 ± 0.043 | 0.405 ± 0.080 |

Among the remaining variants, measurements were consistent and yielded reliable kinetic parameters. SAK SY155 demonstrated the highest turnover number (k_cat_ = 1.94 ± 0.17 s^−1^) and catalytic efficiency (k_cat_/K_M_ = 0.405 ± 0.080 uM^−1^*s^−1^), indicating superior catalytic performance. SAK THR174, another low-immunogenic variant, showed a turnover number comparable to SY155 (k_cat_ = 1.89±0.27 s^−1^), but reduced catalytic efficiency (k_cat_/K_M_ = 0.133 ± 0.043 uM^−1^*s^−1^), due to an elevated K_M_. Among the two wild-type forms, SAK STAR WT displayed higher activity than SAK 42D WT across all parameters.

Next, we evaluated the thrombolytic efficiency of the SAK variants using the clot formation and lysis (CloFAL) assay (Fig. 1B, Table S14). Among the tested variants, SAK SY155 exhibited the fastest clot lysis (14.14-17.01 min), followed by SAK STAR WT (15.29-20.13 min), SAK 42D WT (17.59-23.74 min), and SAK THR174 (22.71-25.72 min). In contrast, SAK STAR FRIDA (44.64-58.11 min) displayed a markedly reduced thrombolytic efficiency. This ranking is consistent with the results obtained from the steady-state kinetics, using the fluorogenic assay, confirming that while four variants show broadly comparable efficiency, SAK STAR FRIDA underperforms.

The affinity of all five variants to their interaction partner plasmin was measured using SPR. The fitting of the SPR binding curves for all variants is shown in the supplement (Fig. S9-13). Reliable estimates were obtained for both the binding equilibrium constants and the association and dissociation rate constants for all tested variants. The equilibrium binding parameters are summarized in Table 2, while the kinetic rates are summarized in Table S16. In contrast to its lower activity, the 42D template (≈ 39 nM) has a higher affinity than the STAR template (≈ 95 nM). SAK STAR FRIDA is again the worst-performing variant by a significant margin (≈ 132 nM). SAK SY155 exhibited the strongest binding (≈ 6 nM), owing mostly to the slower dissociation rates, and again showed superior performance to the other non-immunogenic variant SAK THR174 (≈ 67 nM).

**Table 2.** Dissociation constants of staphylokinase variants binding to human plasmin determined by SPR. Human plasmin was immobilized on a CM5 chip via amine coupling, and staphylokinase (3.7 - 900 nM) was injected as the mobile phase in PBS/Tween 80 (pH 7.4) at 25°C. Binding curves were fitted to a numeric model corresponding to the proposed mechanism. K_d_app_ is the apparent dissociation constant of the two-step induced binding. At the same time, K_d_1_ is the dissociation constant of the first step of the two-step induced binding, and K_d_3_ is the dissociation constant of the parallel unproductive single-step binding.

| SPR affinity kinetic parameters |  |  |  |  |  |
| --- | --- | --- | --- | --- | --- |
|  | SAK STAR WT | SAK 42D WT | SAK STAR FRIDA | SAK THR174 | SAK SY155 |
| $K_{d\_app}$ [nM] | 94.9 ± 19.4 | 38.4 ± 7.1 | 132.1 ± 36.6 | 66.6 ± 40.7 | 6.1 ± 0.8 |
| $K_{d\_1}$ [nM] | 142.5 ± 20.3 | 68.8 ± 12.3 | 294.7 ± 64.5 | 88.1 ± 52.1 | 29.0 ± 2.0 |
| $K_{d\_3}$ [nM] | 15.3 ± 1.1 | 12.5 ± 0.8 | 192.9 ± 43.9 | 22.5 ± 3.4 | 48.0 ± 7.9 |

### Comparison of thrombolytic activities with blood clots in a static model

In the *in vitro* static model, all thrombolytic treatments significantly increased clot lysis, as assessed by relative clot weight loss and RBC release (Fig. 2, Table S1, S2), compared to the control. Among the SAK variants, SAK STAR WT and SAK SY155 exhibited lytic efficacy comparable to that of clinically used ALT (p>0.05 for both), while SAK THR174 and SAK STAR FRIDA were significantly worse than ALT (p<0.05 for both).

**Figure 2.**
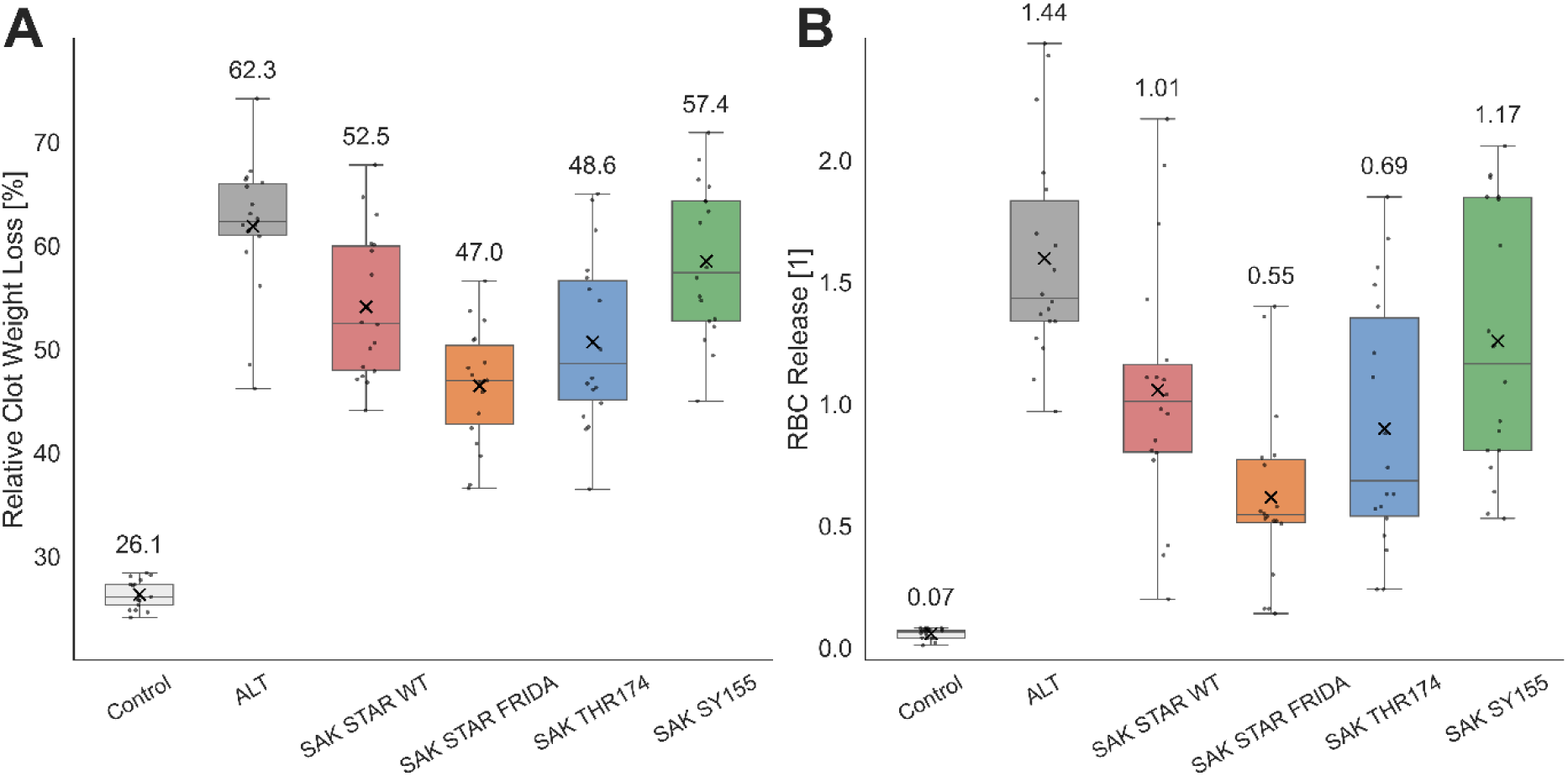
Thrombolytic efficacy in a static *in vitro* model **A)** The results of the relative clot weight loss determination. The number of repetitions ranges from 17 to 18, depending on the variant. The concentration of all agents was 1.3 µg/mL. The experiment was performed in 0.5 mL of human plasma at 37 °C for 60 minutes. **B)** The results of the red blood cell (RBC) release determination. The number of repetitions was 18. The concentration of all agents was 1.3 µg/mL. The box plots show mean value (X), median (line), interquartile ranges (boxes), and minimum and maximum values (whiskers). The value for the mean is also stated above the boxes. The experiment was performed in 0.5 mL of human plasma at 37 °C for 60 minutes; the release was monitored by determination of hemoglobin absorbance at 575 nm against the turbidity at 700 nm.

### Comparison of thrombolytic activities with blood clots in a flow model

In the flow model, all treatments, except SAK SY155 (p=0.221), significantly reduced recanalization time compared to the control (p<0.05; Fig. 3A, Table S3). No significant differences were observed between the individual treatments. RBC release analysis confirmed enhanced lytic efficacy for all treatments compared to the control (p<0.05 for each group; Fig. 3B, Table S4). Clot area evaluation revealed that ALT, SAK STAR WT, SAK STAR FRIDA, and SAK THR174 produced a significant reduction compared to the control after 60 min (p<0.05; Fig. 3C, Table S5), whereas SAK SY155 had no significant effect at 60 min. By 120 min, all treatments achieved a significant reduction in clot size compared to the control (p<0.001 for all; Fig. 3C, Table S5), with no significant differences among the treatments. At 60 min, SAK STAR WT achieved a significantly higher recanalization frequency than the control (p=0.006; Table 3, S6). By 120 min, all thrombolytics had significantly increased recanalization frequencies compared to the control (p<0.05 for all groups; Table 3, S6), with no significant differences among the treatments.

**Figure 3.**
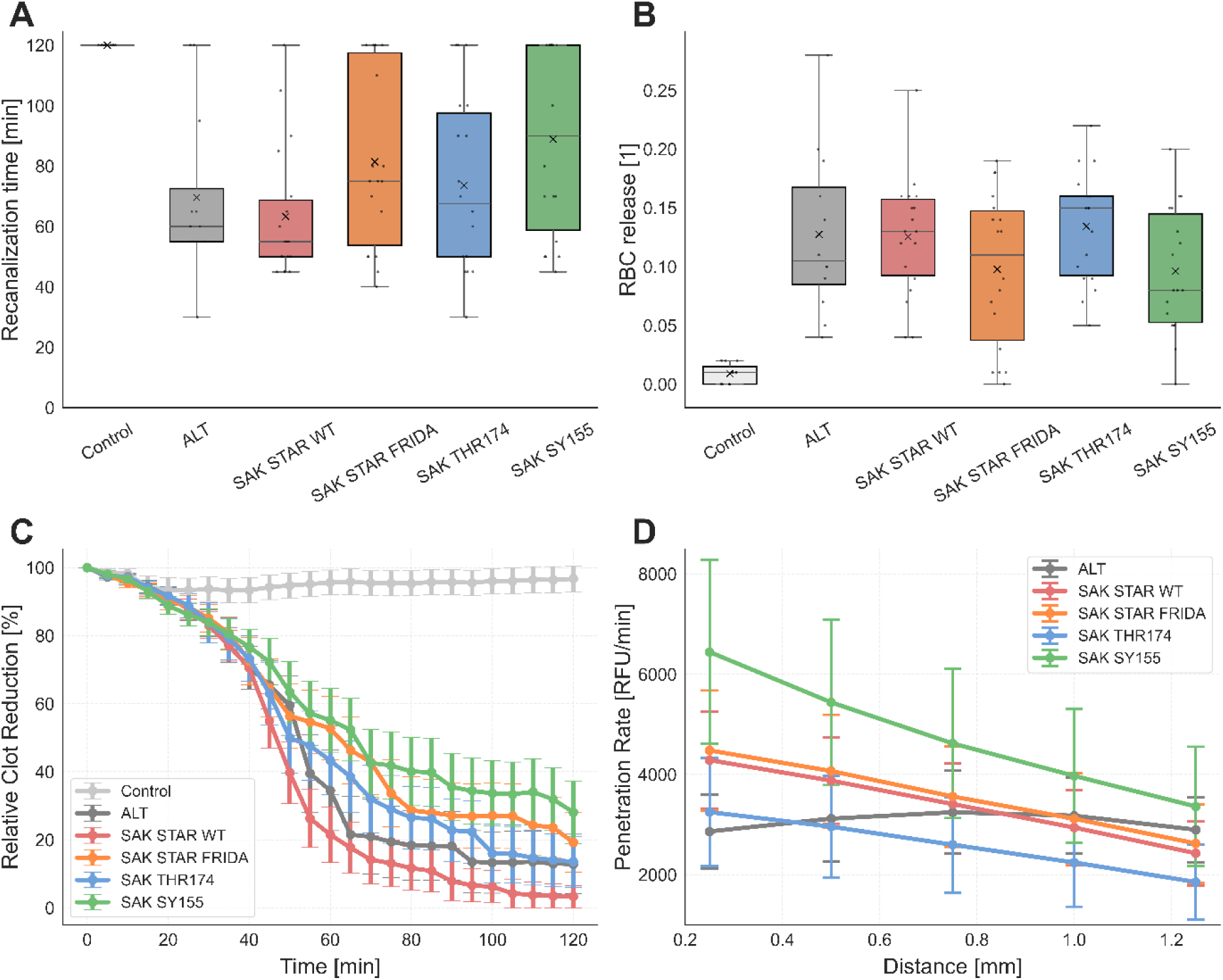
Clot lysis using an *in vitro* flow model and the clot penetration experiment. **A)** Recanalization time. Experiments were conducted at 37 °C for 120 minutes in human plasma, with 11–22 repetitions per condition. All agents were used at a concentration of 1.3 µg/mL. Box plots indicate the mean (X), median (line), interquartile range (boxes), and minimum and maximum values (whiskers). **B)** Red blood cell (RBC) release. Experiments were conducted at 37 °C for 120 minutes in human plasma, with 11–18 repetitions per condition. RBC release was monitored by the determination of hemoglobin absorbance at 575 nm against the turbidity at 700 nm. Box plots indicate the mean (X), median (line), interquartile range (boxes), and minimum and maximum values (whiskers). **C)** Relative clot reduction over time. Data are shown as mean ± standard error of the mean at 5-minute intervals. Experiments were conducted at 37 °C for 120 minutes in human plasma, with 11–18 repetitions per condition. All agents were used at 1.3 µg/mL. **D)** Clot penetration rates. Data are presented as mean ± standard deviation of the mean of the penetration rate versus distance. Experiments were conducted at 37 °C for 180 minutes in PBS, with 19-23 repetitions per condition. All agents were used at a concentration of 0.022 µM.

**Table 3.** Descriptive statistics of the recanalization frequency in the flow model. Thrombolytic agent efficacy expressed as recanalization frequency at 60 min and 120 min. Experiments were conducted at 37 °C for 120 minutes in human plasma, with 11–18 repetitions per condition. All agents were used at a concentration of 1.3 µg/mL.

| Recanalization frequency: Descriptive statistics |  |  |  |  |  |  |
| --- | --- | --- | --- | --- | --- | --- |
|  | Control | ALT | SAK STAR<br>WT | SAK STAR<br>FRIDA | SAK<br>THR174 | SAK<br>SY155 |
| **At 60 minutes** |  |  |  |  |  |  |
| Recanalization | 0 (0%) | 6 (50%) | 12 (67%) | 5 (28%) | 8 (44%) | 5 (28%) |
| No recanalization | 11 (100%) | 6 (50%) | 6 (33%) | 13 (72%) | 10 (56%) | 13 (72%) |
| Count (100%) | 11 | 12 | 18 | 18 | 18 | 18 |
| **At 120 minutes** |  |  |  |  |  |  |
| Recanalization | 0 (0%) | 10 (83%) | 17 (94%) | 14 (78%) | 15 (83%) | 11 (61%) |
| No recanalization | 11 (100%) | 2 (17%) | 1 (6%) | 4 (22%) | 3 (17%) | 7 (39%) |
| Count (100%) | 11 | 12 | 18 | 18 | 18 | 18 |

### Penetration efficiency testing using the clot penetration assay

Distance-dependent analysis revealed significant negative correlations between penetration rate and depth for all SAK variants (rs = −1.0, p<0.017; Fig. 3D, Table S7), whereas ALT showed no correlation (rs = 0.3; p=0.683; Fig. 3D, Table S7). At 0.25 mm, SAK STAR WT, SAK STAR FRIDA, and SAK SY155 exhibited higher penetration rates than ALT (p<0.05; Fig. S1, Table S8), with SAK SY155 also outperforming SAK STAR WT, SAK STAR FRIDA, and SAK THR174 (p<0.001; Fig. S1, Table S8). SAK SY155 maintained higher rates than ALT up to 0.75 mm and, together with SAK STAR FRIDA, surpassed SAK THR174 across all distances up to 1.25 mm (p<0.05; Fig. S1, Table S8). Penetration time analysis (Fig. S2, Table S9) confirmed that SAK SY155 reached thresholds significantly faster than ALT (0.25-0.50 mm, p<0.05), SAK STAR WT (0.25, 0.75-1.25 mm, p<0.05), and SAK THR174 (across all distances, p<0.001). The ALT showed shorter penetration times than the SAK THR174 at 0.75 and 1.0 mm (p<0.05). Penetration frequency analysis (Table S10) revealed that ALT and SAK SY155 exceeded SAK THR174 at 0.75 and 1.00 mm (p<0.05). SAK SY155 further demonstrated higher success than ALT and SAK THR174 at 1.25 mm (p<0.05).

### MAT test

The MAT test was performed to confirm the removal of endotoxin from the SAK variants. As expected, the assay controls confirmed validity: both the negative control and LPS at 0.007 EU/mL, corresponding to the maximal level permitted in intravenously administered drugs, induced only very low cytokine release. In contrast, a 10-fold higher concentration of LPS (0.07 EU/mL) triggered a markedly stronger cytokine release, thereby confirming the proper functionality and dynamic range of the assay (Fig. S18A, Table S12). The clinically used reference, ALT, did not induce significant cytokine release at either the clinically relevant concentration (1.3 μg/mL) or the 10-fold higher concentration (13 μg/mL), with responses comparable to those of the negative control and 0.007 EU/mL LPS. Similarly, none of the SAK variants triggered cytokine release above this baseline level at either concentration (Fig. S18A, Table S12).

### Immunogenicity testing using T lymphocyte proliferation

T lymphocyte proliferation was evaluated in CD8-depleted PBMCs from four healthy donors (Fig. S18B, Table S13). The negative control induced only minimal proliferation, whereas the positive control (PHA) elicited robust proliferative responses (p<0.001) compared to the negative control and all treatment variants, confirming cell viability and validating assay performance. Neither ALT nor any of the SAK variants induced proliferation significantly above the negative control at either tested concentration (p>0.05).

### Determination of autoantibodies in the sera of immunized mice

Using human diagnostic immunoblot kits, we tested the ability of individual SAK variants to induce antibodies cross-reacting with known human organ-nonspecific autoantigens as a marker of potential autoimmune reaction elicitation by SAK. No borderline or positive results were detected for individual autoantigens present in diagnostic immunoblots with any of the SAK-immunized mouse groups. This experiment did not identify any evidence about the ability of SAK proteins to induce antibodies cross-reacting with autoantigens commonly recognized in human systemic autoimmune diseases (Figures S14, S15, S16).

### Induction of specific antibody response after vaccination with individual SAK variants

We compared the immunogenicity of individual SAK types in experimental mice immunized either with individual SAK proteins or with a combination of SAK protein and aluminum adjuvant. Serum level of individual SAK protein after two immunizations was evaluated using the ELISA method (Fig. S20, Table S15) and serum-inhibited SAK activity measurement.

In the ELISA method, all groups of mice were tested on plates coated with each of the SAK variants used for vaccination. In the groups vaccinated with the protein alone, only very low levels of antibodies were detected, in most groups comparable to negative sera from unvaccinated mice (Table S15). Antibody levels were an order of magnitude higher in the groups of mice vaccinated with aluminum-adjuvanted SAK proteins (Fig. S20). For most SAKs, the highest reactivity was observed in the group vaccinated with the same variant as was used to coat the individual ELISA plates, but differences between groups were minimal and statistically insignificant due to the high level of cross-reactivity. The result indicates only a marginal effect of the sequence variation among the tested SAKs on their overall antigenicity. The low antibody response of mice immunized with individual SAK variants without adjuvants indicates low, if any, spontaneous immunogenicity and generally high SAK purity devoid of contaminants, which could otherwise stimulate the immune system to antigen-specific response. Well detectable antibody response to SAK plus adjuvant immunization indicates that, at least in mice, there is no active tolerance to the tested SAK variants.

In the inhibitory potency measurement, the activity in the presence of each serum was assessed and normalized to the negative control serum activity, which is set to 100%. As shown in Figure S19, as with ELISA, no significant differences between variants were observed. For all variants, the adjuvant-free serum showed similar or higher activity, while adjuvant-rich serum generally had lower activity than the negative control (the exception here was SAK STAR FRIDA). This experiment suggests that, without an adjuvant, no variant induces significant antibody generation.

Furthermore, we compared the reactivity of a set of randomly selected human sera to all five SAK variants and irrelevant protein – human myomesin. Results are shown in Figures 4 and S21. Variants SAK SY155 and SAK THR174 exhibited a significant reduction in reactivity with sera in comparison to both wild types SAK 42D (p < 0.01) and SAK STAR (p < 0.0001), respectively. Although variant SAK STAR FRIDA also exhibited reduced reactivity (PC score for SAK STAR FRIDA 1.022 versus 1.78 for SAK 42D or 3.80 for SAK STAR), it did not reach statistical significance.

**Figure 4.**
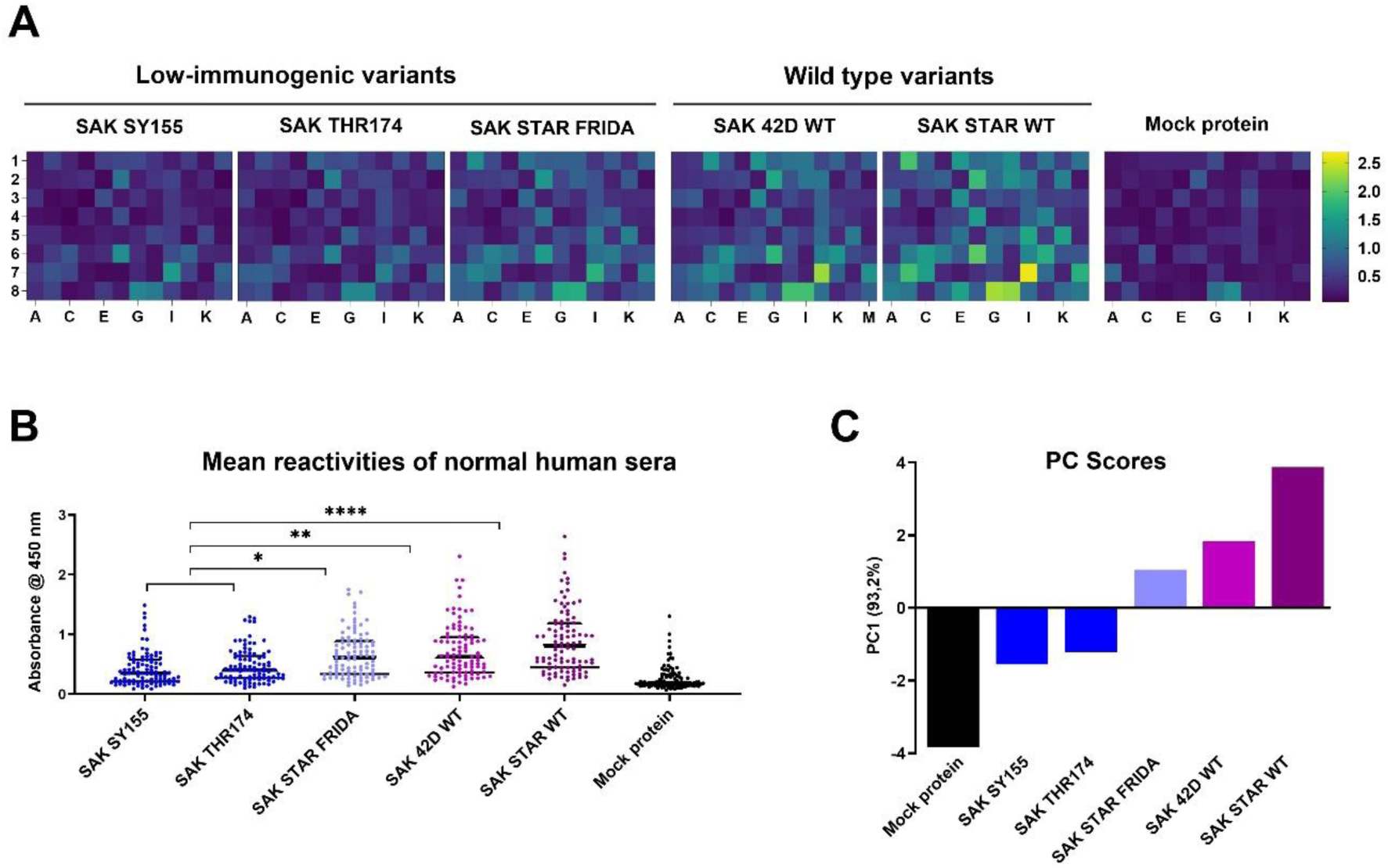
Reactivity of human sera with individual SAK variants. A) Frozen-stored human sera from randomly selected healthy subjects providing informed consent were tested for reactivity with individual recombinant SAK variants coated on the ELISA plates. Analyses were performed with anti-human IgG γ-chain HRP-conjugated antibody in two independent replicates, and mean values of absorbances for each serum are presented in heat maps for each SAK. B) Individual absorbances are present in dots, and median +/− quartiles are shown in black lines. Data were analyzed for statistical significance using the Kruskal-Wallis test followed by Dunn’s multiple comparisons test. *, **, **** denote p < 0.05, 0.01, 0.0001, respectively. C) The first principal component of the human sera reactivity (mean values of absorbances) separates the immunogenic and low-immunogenic SAK variants, capturing 93.2% of the variance in the data distribution. The immunogenicity follows the order: SAK SY155 < SAT THR174 < SAK FRIDA < SAK 42D < SAK STAR.

### Analysis of dendritic cell activation

Flow cytometry analysis tested the potency of individual SAK variants to activate murine dendritic cells *in vitro*. The intensity of activation was compared with the negative control (medium plus phosphate buffer) group, labeled Ctrl, and the positive control LPS using several surface markers (MHCII, CD40, CD69, CD80, and CD86). No significant activation over the control (Ctrl) group was detected for most of the tested SAK variants (SAK STAR WT, SAK STAR FRIDA, and SAK SY155 using dendritic cell activation markers (Fig. 5A). In the case of the SAK THR174 variant, a significant but biologically moderate increase in the number of CD40 high cell population (58%) over the Ctrl group (52%)(Fig. 5).

**Figure 5.**
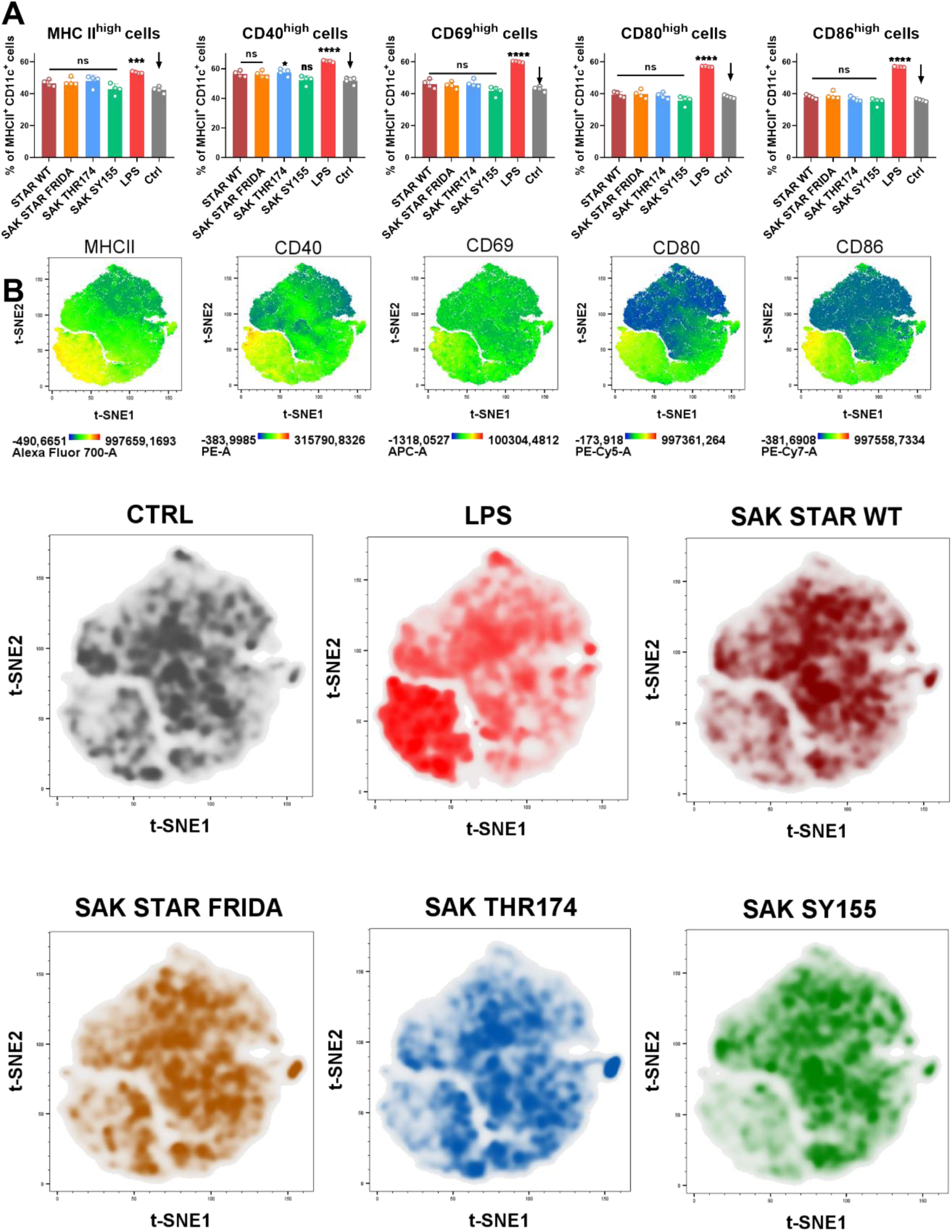
Analysis of bone marrow-derived dendritic cells activation mediated by different SAK variants. **A)** Activation of BMDCs mediated by different SAK was characterized by determining MHCII high, CD40 high, CD69 high, CD80 high and CD86 high cell populations in CD11c+ MHCII+ cells. **B)** Visualisation of high-dimensional data describing the potential of SAK variants to activate dendritic cells in vitro was performed using t-SNE (t-distributed stochastic neighbour embedding). LPS (150 ng/ml) as a positive control and PBS as a negative control were used. Significance values were calculated by the one-way ANOVA and Dunnett’s post hoc test, where *p < 0.05 and **** p < 0.0001.

t-SNE visualization shows the expression of particular markers by heatmap layouts, from where it is evident, that the MHCII high, CD40 high, CD69 high, CD80 high and CD86 high cell populations (yellow color) are focused in the lower left quadrants of the heatmaps marked for each characterized marker (Fig. 5B). Analyzed SAK variants together with LPS as positive control and PBS as negative control are expressed as density plots. Whereas the most cells exposed to individual SAK variants are localized in the middle of the graph, similarly to the Ctrl group, LPS activation shifts the cells to the lower left part of the density graph. Our results show that generally all the SAK variants (SAK STAR WT, SAK STAR FRIDA, SAK THR174, and SAK SY155) show a similar pattern as the negative control compared to the LPS group (Fig. 5B). From these results, it could be summarized that none of the SAK variants can effectively activate murine BMDC cells *in vitro*, with the only exception of SAK THR174, where the results indicate a moderate potential to increase a population of activated CD40 high BMDC.

## Discussion

To accelerate the development of safer and more effective thrombolytics, it is essential to identify the most suitable starting scaffold ^55^. We conducted a systematic comparative analysis of the functional and immunogenic properties of selected SAK wild-types (SAK STAR and SAK 42D) and engineered low-immunogenic variants SAK SY155, SAK THR174, and SAK STAR FRIDA. The aim was to determine the variant with the most favourable molecular profile to serve as a template for the design of the next generation of thrombolytics.

Biochemical characterization, including production yields, thermal stability, aggregation propensity, together with enzyme kinetics, yielded two main conclusions. First, SAK STAR WT significantly outperforms SAK 42D WT template across all measured parameters, except affinity. Second, the engineered variants differ considerably in kinetic performance: SAK SY155 shows the highest catalytic efficiency and affinity towards plasmin, surpassing even the SAK STAR WT, whereas SAK THR174 performs slightly worse, and SAK STAR FRIDA is markedly weaker. In SPR measurements, the SAK 42D template showed higher binding affinity (∼39 nM) than the SAK STAR template (∼95 nM), despite its lower activity. SAK STAR FRIDA performed the worst (∼132 nM), while SAK SY155 was the strongest binder (∼6 nM), largely due to slow dissociation, outperforming the other low-immunogenic variant SAK THR174 (∼67 nM). These kinetic differences are particularly notable given the absence of complementary structural insights into the interaction between staphylokinase and full-length plasmin and plasminogen, as only the interaction with microplasmin has been studied by X-ray crystallography so far ^56^.

In static thrombolysis assays, all SAK variants enhanced clot lysis compared with the control. SAK STAR WT, SAK THR174, and especially SAK SY155 achieved efficacy comparable to alteplase, consistent with their kinetic profiles. However, under flow conditions, the performance ranking changed: SAK SY155 showed the weakest activity, followed by SAK STAR FRIDA. These discrepancies likely reflect flow-dependent transport phenomena, including the balance between diffusion and convection and variant-specific fibrin-binding dynamics ^57–60^. Diffusion experiments further supported this view, revealing that SAK SY155 penetrates fibrin gels more efficiently than other SAK variants and alteplase, while SAK THR174 exhibits reduced penetration, mirroring its static assay performance. Possible limitations of these findings are the use of flow rates lower than those typically observed *in vivo*, which may reduce the pressure gradients across the clot. This adjustment was necessary to maintain clot stability while preserving susceptibility to enzymatic degradation. Although both static and flow-based *in vitro* models lack excretory mechanisms, this simplification allowed us to sustain a consistent local concentration of thrombolytic agents without continuous plasma supplementation. The concentrations of alteplase and SAK variants were chosen to reflect therapeutically relevant levels, corresponding to clinical alteplase dosing in ischemic stroke treatment ^45^.

Immunogenicity, a major barrier to address for clinical deployment of SAK ^20,24,61,62^, was assessed using several complementary approaches: (i) T-cell proliferation, (ii) human autoantigen cross-reactivity, (iii) ELISA detection of SAK-specific antibodies developed in immunized mice, (iv) inhibitory potency of SAK-specific antibodies developed in immunized mice, (v) reactivity of human sera with individual SAK variants, and (vi) murine dendritic-cell activation assays. None of the tested SAK variants activated T lymphocyte proliferation. Moreover, none of the variants induced human autoantigen-cross-reactive antibodies developed after immunization of experimental mice, and non-adjuvanted SAK failed to elicit detectable antibody responses in mice. Only under strong inflammatory stimulation using aluminum adjuvant, all variants generate SAK-specific antibodies, consistent with their nature as foreign proteins. None of the tested SAK variants activated murine dendritic cells *in vitro*, suggesting a limited innate immune stimulatory potential. Finally, in the human serum reactivity test, we detected significant differences between the variants. The experiment showed that the low-immunogenic variants SAK THR174 and SAK SY155 exhibit lower reactivity with human sera than either wild type (SAK STAR WT or SAK 42D WT). The third studied low-immunogenic variant, SAK STAR FRIDA, exhibits lower reactivity as well, but this decrease is not statistically significant.

These experiments are in agreement with previously published differences in immunogenicity of these variants ^26,29^. However, the differences are only observable when human sera are used, not in mice, especially in inbred pathogen-free mice, which do not fully reflect the reality in an HLA-variable human population exposed to SAK after potential prior contact with Staphylococcus sp. bacteria, a natural producer of SAK. Future studies evaluating immunogenicity under repeated-dosing conditions and using human cells and sera samples will be essential to fully determine the clinical safety profile of SAK variants.

Overall, our findings reveal clear variant-specific differences in catalytic behaviour, transport dynamics, thrombolytic efficacy, and immunogenicity. These insights highlight the wild-type SAK STAR and non-immunogenic variant SAK SY155 as the most promising scaffolds for further engineering, while emphasising the importance of considering both biochemical properties and physiological flow conditions in the development of next-generation thrombolytic agents.

## Conclusions

This study reconfirms that staphylokinase (SAK) is a potent thrombolytic agent with catalytic efficacy comparable to alteplase. Importantly, this activity was maintained across all tested variants, even after introducing mutations designed to reduce immunogenicity. While the SAK STAR FRIDA and SAK THR174 variants showed small to moderate decreases in activity, the SAK SY155 variant displayed higher activity and affinity with plasmin than SAK STAR WT. Moreover, SAK SY155 showed significantly higher permeability than alteplase, adding to its favorable profile. Although this did not yet translate into greater clot reduction under flow conditions, low-immunogenic SAK SY155 is the most promising candidate for further engineering. Our study also confirmed that low-immunogenic variants SAK THR174 and SAK SY155 significantly decrease reactivity with the human immune system.

## Acknowledgements

This project has received funding from the European Union’s Horizon 2020 research and innovation programme under grant agreements No. 857560 (CETOCOEN) and Horizon Europe Framework programme No. 101136607 (CLARA). ZP acknowledges the Czech-Swiss grant No. 25-15784L and DB grant No. 25-18233M supported by the Czech Grant Agency. MP and JV were supported by the Ministry of Health of the Czech Republic in cooperation with the Czech Health Research Council under project No. NW24-08-00064. Computational resources were provided by the e-INFRA CZ, ELIXIR-CZ, and RECETOX RI projects (Nos. 90254, LM2023055, and LM2023069), supported by the Ministry of Education, Youth and Sports of the Czech Republic. AS and JM are supported by the scholarship Brno Ph.D. Talent. CIISB, Instruct-CZ Centre of Instruct-ERIC EU consortium, funded by MEYS CR infrastructure project LM2023042 and European Regional Development Fund-Project „Innovation of Czech Infrastructure for Integrative Structural Biology” (No. CZ.02.01.01/00/23_015/0008175), is gratefully acknowledged for the financial support of the measurements at the CF Biomolecular Interactions and Crystallography. This publication reflects only the authors‘ view, and the European Commission is not responsible for any use that may be made of the information it contains.

## Author contributions

Alan Strunga – Laboratory work (protein production, endotoxin removal, and biochemical characterisation – activity, affinity, stability, inhibition assay), data evaluation and interpretation, conceptualization, manuscript writing (all parts), preparation of the figures Michaela Peskova - Laboratory work (thrombolysis assays in vessel models, penetration experiments), data evaluation and interpretation, manuscript writing (methodology, results, discussion)

Michal Krupka - Laboratory work (endotoxin testing and removal, immunization experiments on the mouse model, evaluation of the humoral immune response), data evaluation and interpretation, manuscript writing (methodology, results, discussion)

Katerina Zachova - Laboratory work (flow cytometry analysis), manuscript writing (methodology, results, discussion), preparation of figures

Lenka Sindlerova – Laboratory work (immunology – MAT test, T cell proliferation), manuscript writing (methodology, results, discussion), data evaluation and interpretation, supervision Nikola Neumeisterova - Laboratory work (immunology – MAT test, T cell proliferation), manuscript writing (methodology, results, discussion)

Jan Mican – Manuscript revisions

Jan Vitecek – Data interpretation (thrombolysis assays in vessel models, penetration experiments)

David Bednar - Manuscript revisions, supervision

Lukas Kubala - Data interpretation (immunological characterisation), manuscript writing (methodology, results, discussion), manuscript revisions, supervision

Milan Raska – Data interpretation (immunological characterisation), conceptualization, manuscript revisions, supervision

Stanislav Mazurenko – Data interpretation (PCA analysis)

Jiri Damborsky – Data interpretation, conceptualization, manuscript revisions, supervision

Zbynek Prokop – Kinetic data analysis, data interpretation, conceptualization, manuscript revisions, supervision

