## Supporting information for "Biochemical and Immunological Properties of Engineered Low-Immunogenic Staphylokinases for Next-Generation Thrombolytic Therapy"

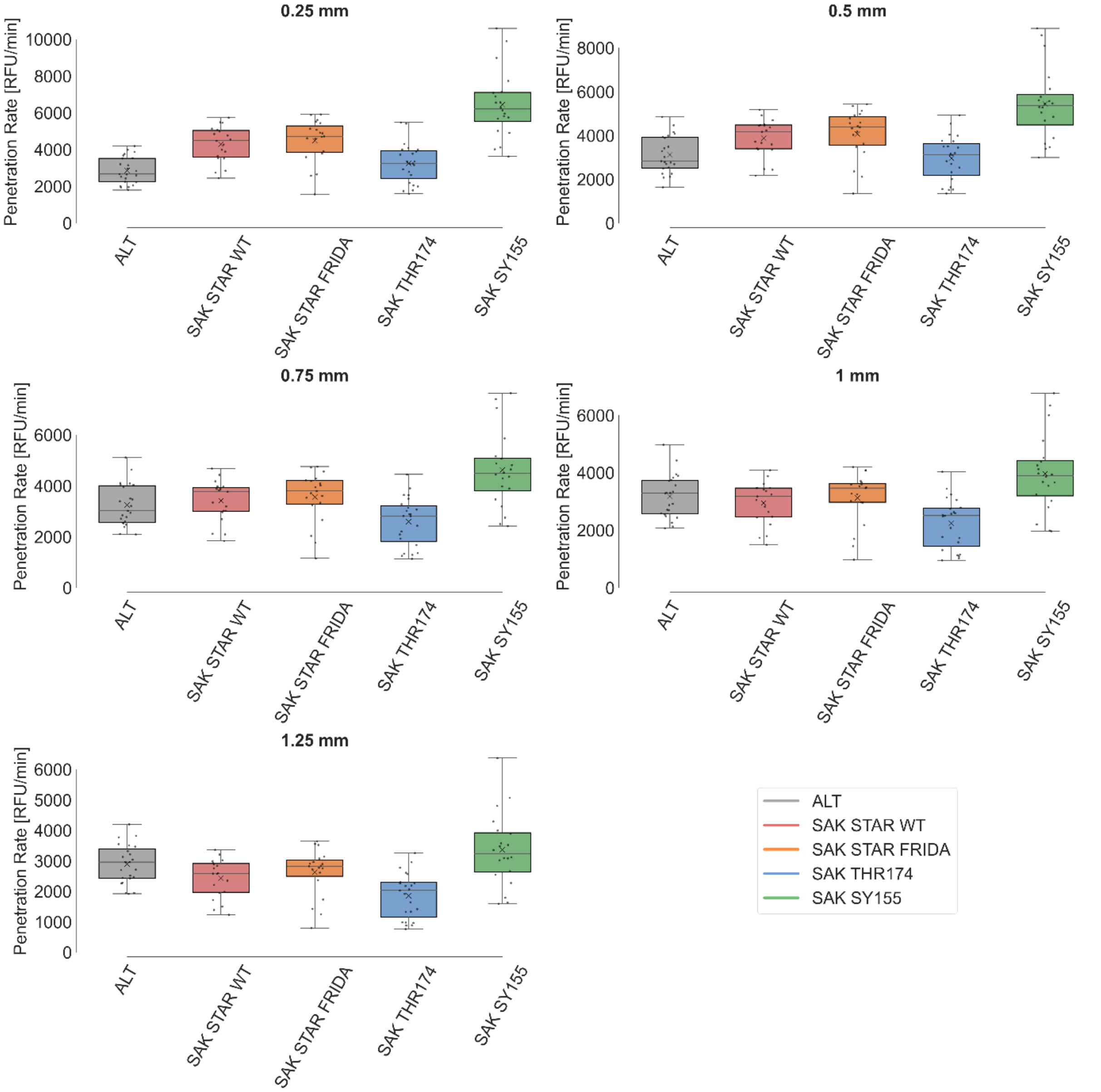


**Figure S1** Penetration rates at individual distances. Experiments were conducted at 37 °C for 180 minutes in PBS, with 19-23 repetitions per condition. All agents were used at the concentration of 0.022 μM. The box plots show mean value (X), median (line), interquartile ranges (boxes), and minimum and maximum values (whiskers).

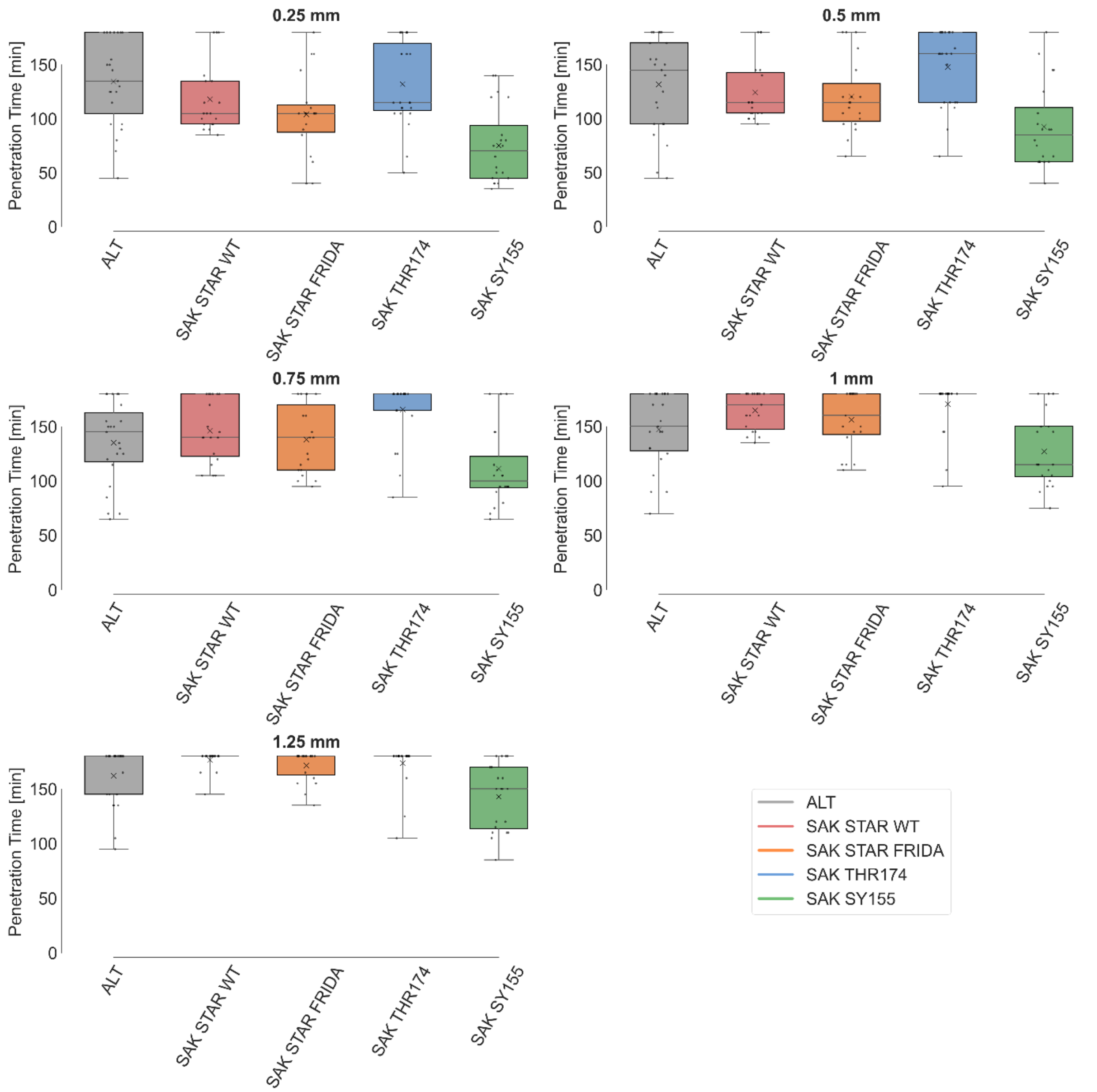


**Figure S2** Penetration time at individual distances. Experiments were conducted at 37 °C for 180 minutes in PBS, with 19-23 repetitions per condition. All agents were used at a concentration of 0.022 µM. The box plots show mean value (X), median (line), interquartile ranges (boxes), and minimum and maximum values (whiskers).

**Table S1** Static model: descriptive statistics and p-values for clot lysis expressed as relative weight loss. The concentration of all agents was 1.3 µg/mL. The experiment was performed in 0.5 mL of human plasma at 37 °C for 60 minutes **A)** Descriptive statistics for each group (mean, SD = standard deviation, median, n = number of replicates). **B)** Pairwise p-values from Kruskal-Wallis test with post-hoc Dunn's correction. Statistical significance was set at p<0.05.

| **A) Clot weight loss: Descriptive statistics** | | | | | | |
|---|---|---|---|---|---|---|
| | **Control** | **ALT** | **SAK STAR WT** | **SAK STAR FRIDA** | **SAK THR174** | **SAK SY155** |
| **Mean [%]** | 26.3 | 61.9 | 54.1 | 46.5 | 50.7 | 58.5 |
| **SD [%]** | 1.4 | 6.5 | 7.0 | 5.6 | 8.2 | 7.4 |
| **Median [%]** | 26.1 | 62.3 | 52.5 | 47.0 | 48.6 | 57.4 |
| **Count** | 17 | 18 | 18 | 18 | 18 | 18 |

| **B) Relative clot weight loss: p-values** | | | | | | |
|---|---|---|---|---|---|---|
| | **Control** | **ALT** | **SAK STAR WT** | **SAK STAR FRIDA** | **SAK THR174** | **SAK SY155** |
| **Control** | - | <0.001 | <0.001 | 0.049 | 0.001 | <0.001 |
| **ALT** | - | - | 0.485 | <0.001 | 0.020 | >0.999 |
| **SAK STAR WT** | - | - | - | 0.495 | >0.999 | >0.999 |
| **SAK STAR FRIDA** | - | - | - | - | >0.999 | 0.008 |
| **SAK THR174** | - | - | - | - | - | 0.255 |

**Table S2** Static model: descriptive statistics and p-values for clot lysis expressed as RBC release. The concentration of all agents was 1.3 µg/mL. The experiment was performed in 0.5 mL of human plasma at 37 °C for 60 minutes **A)** Descriptive statistics for each group (mean, SD = standard deviation, median, n = number of replicates). **B)** Pairwise p-values from Kruskal-Wallis test with post-hoc Dunn's correction. Statistical significance was set at p<0.05.

| **A) RBC release: Descriptive statistics** | | | | | | |
|---|---|---|---|---|---|---|
| | **Control** | **ALT** | **SAK STAR WT** | **SAK STAR FRIDA** | **SAK THR174** | **SAK SY155** |
| **Mean [1]** | 0.06 | 1.60 | 1.06 | 0.62 | 0.90 | 1.26 |
| **SD [1]** | 0.02 | 0.44 | 0.52 | 0.36 | 0.52 | 0.55 |
| **Median [1]** | 0.07 | 1.44 | 1.01 | 0.55 | 0.69 | 1.17 |
| **Count** | 18 | 18 | 18 | 18 | 18 | 18 |

| **B) RBC release: p-values** | | | | | | |
|---|---|---|---|---|---|---|
| | **Control** | **ALT** | **SAK STAR WT** | **SAK STAR FRIDA** | **SAK THR174** | **SAK SY155** |
| **Control** | - | <0.001 | <0.001 | 0.039 | <0.001 | <0.001 |
| **ALT** | - | - | 0.389 | <0.001 | 0.044 | >0.999 |
| **SAK STAR WT** | - | - | - | 0.492 | >0.999 | >0.999 |
| **SAK STAR FRIDA** | - | - | - | - | >0.999 | 0.050 |
| **SAK THR174** | - | - | - | - | - | >0.999 |

**Table S3** Flow model: descriptive statistics and p-values for thrombolytic agent efficacy expressed as recanalization time. The concentration of all agents was 1.3 µg/mL. The experiment was performed at 37 °C for 120 minutes in human plasma **A)** Descriptive statistics for each group (mean, SD = standard deviation, median, n = number of replicates). **B)** Pairwise p-values from Kruskal-Wallis test with post-hoc Dunn's correction. Statistical significance was set at p<0.05.

| **A) Recanalization time: Descriptive statistics** | | | | | | |
|---|---|---|---|---|---|---|
| | **Control** | **ALT** | **SAK STAR WT** | **SAK STAR FRIDA** | **SAK THR174** | **SAK SY155** |
| **Mean [min]** | 120 | 70 | 63 | 81 | 74 | 89 |
| **SD [min]** | 0 | 28 | 22 | 30 | 29 | 31 |
| **Median [min]** | 120 | 60 | 55 | 75 | 68 | 90 |
| **Count** | 11 | 12 | 18 | 18 | 18 | 18 |

| **B) Recanalization time: p-values** | | | | | | |
|---|---|---|---|---|---|---|
| | **Control** | **ALT** | **SAK STAR WT** | **SAK STAR FRIDA** | **SAK THR174** | **SAK SY155** |
| **Control** | - | 0.007 | <0.001 | 0.034 | 0.002 | 0.211 |
| **ALT** | - | - | >0.999 | >0.999 | >0.999 | >0.999 |
| **SAK STAR WT** | - | - | - | 0.912 | >0.999 | 0.159 |
| **SAK STAR FRIDA** | - | - | - | - | >0.999 | >0.999 |
| **SAK THR174** | - | - | - | - | - | >0.999 |

**Table S4** Flow model: descriptive statistics and p-values for clot lysis expressed as RBC release. The concentration of all agents was 1.3 µg/mL. The experiment was performed at 37 °C for 120 minutes in human plasma **A)** Descriptive statistics for each group (mean, SD = standard deviation, median, n = number of replicates). **B)** Pairwise p-values from Kruskal-Wallis test with post-hoc Dunn's correction. Statistical significance was set at p<0.05.

| **A) RBC release: Descriptive statistics** | | | | | | |
|---|---|---|---|---|---|---|
| | **Control** | **ALT** | **SAK STAR WT** | **SAK STAR FRIDA** | **SAK THR174** | **SAK SY155** |
| **Mean [1]** | 0.01 | 0.13 | 0.13 | 0.10 | 0.13 | 0.10 |
| **SD [1]** | 0.01 | 0.07 | 0.05 | 0.07 | 0.05 | 0.05 |
| **Median [1]** | 0.01 | 0.11 | 0.13 | 0.11 | 0.15 | 0.08 |
| **Count** | 11 | 12 | 18 | 18 | 18 | 18 |

| **B) RBC release: p-values** | | | | | | |
|---|---|---|---|---|---|---|
| | **Control** | **ALT** | **SAK STAR WT** | **SAK STAR FRIDA** | **SAK THR174** | **SAK SY155** |
| **Control** | - | 0.001 | <0.001 | 0.006 | <0.001 | 0.010 |
| **ALT** | - | - | >0.999 | >0.999 | >0.999 | >0.999 |
| **SAK STAR WT** | - | - | - | >0.999 | >0.999 | >0.999 |
| **SAK STAR FRIDA** | - | - | - | - | >0.999 | >0.999 |
| **SAK THR174** | - | - | - | - | - | 0.835 |

**Table S5** Flow model: descriptive statistics and p-values for thrombolytic agent efficacy expressed as relative clot reduction. The concentration of all agents was 1.3 µg/mL. The experiment was performed at 37 °C for 120 minutes in human plasma **A)** Descriptive statistics for each group (mean, SD = standard deviation, median, n = number of replicates). **B)** Pairwise p-values from Kruskal-Wallis test with post-hoc Dunn's correction. Statistical significance was set at $p<0.05$.

| **A) Relative clot reduction: Descriptive statistics** | | | | | | |
|---|---|---|---|---|---|---|
| | **Control** | **ALT** | **SAK STAR WT** | **SAK STAR FRIDA** | **SAK THR174** | **SAK SY155** |
| **At 60 minutes** | | | | | | |
| Mean [%] | 4.4 | 65.5 | 78.5 | 47.3 | 56.7 | 44.9 |
| SD [%] | 12.1 | 41.0 | 34.4 | 39.6 | 42.9 | 39.8 |
| Median [%] | 2.1 | 88.6 | 100.0 | 31.4 | 60.5 | 33.2 |
| Count | 11 | 12 | 18 | 18 | 18 | 18 |
| **At 120 minutes** | | | | | | |
| Mean [%] | 3.3 | 87.1 | 96.7 | 80.8 | 86.5 | 71.9 |
| SD [%] | 12.6 | 30.3 | 14.1 | 37.5 | 31.9 | 38.8 |
| Median [%] | -0.3 | 100.0 | 100.0 | 100.0 | 100.0 | 100.0 |
| Count | 11 | 12 | 18 | 18 | 18 | 18 |

| **B) Relative clot reduction: p-values** | | | | | | |
|---|---|---|---|---|---|---|
| | **Control** | **ALT** | **SAK STAR WT** | **SAK STAR FRIDA** | **SAK THR174** | **SAK SY155** |
| **At 60 minutes** | | | | | | |
| Control | - | 0.004 | <0.001 | 0.044 | 0.009 | 0.069 |
| ALT | - | - | >0.999 | >0.999 | >0.999 | >0.999 |
| SAK STAR WT | - | - | - | 0.520 | >0.999 | 0.348 |
| SAK STAR FRIDA | - | - | - | - | >0.999 | >0.999 |
| SAK THR174 | - | - | - | - | - | >0.999 |
| **At 120 minutes** | | | | | | |
| Control | - | <0.001 | <0.001 | <0.001 | <0.001 | <0.001 |
| ALT | - | - | >0.999 | >0.999 | >0.999 | >0.999 |
| SAK STAR WT | - | - | - | >0.999 | >0.999 | 0.780 |
| SAK STAR FRIDA | - | - | - | - | >0.999 | >0.999 |
| SAK THR174 | - | - | - | - | - | >0.999 |

**Table S6** Flow model: p-values for thrombolytic agent efficacy expressed as recanalization frequency. The concentration of all agents was 1.3 µg/mL. The experiment was performed at 37 °C for 120 minutes in human plasma. Statistical analysis was performed using Fisher's exact test with post-hoc Holm–Bonferroni correction. Statistical significance was set at $p<0.05$.

| Recanalization frequency: p-values | | | | | | |
|---|---|---|---|---|---|---|
| | **Control** | **ALT** | **SAK STAR WT** | **SAK STAR FRIDA** | **SAK THR174** | **SAK SY155** |
| **At 60 minutes** | | | | | | |
| Control | - | 0.089 | 0.006 | 0.547 | 0.122 | 0.492 |
| ALT | - | - | >0.999 | >0.999 | >0.999 | >0.999 |
| SAK STAR WT | - | - | - | 0.233 | >0.999 | 0.214 |
| SAK STAR FRIDA | - | - | - | - | >0.999 | >0.999 |
| SAK THR174 | - | - | - | - | - | >0.999 |
| **At 120 minutes** | | | | | | |
| Control | - | 0.001 | <0.001 | 0.001 | <0.001 | 0.011 |
| ALT | - | - | >0.999 | >0.999 | >0.999 | >0.999 |
| SAK STAR WT | - | - | - | >0.999 | >0.999 | 0.162 |
| SAK STAR FRIDA | - | - | - | - | >0.999 | >0.999 |
| SAK THR174 | - | - | - | - | - | >0.999 |

**Table S7** Penetration assay: Spearman's correlation between penetration rate and depth for different thrombolytic agents. Experiments were conducted at 37 °C for 180 minutes in PBS, with 19-23 repetitions per condition. All agents were used at the concentration of 0.022 µM. Values represent Spearman's rank correlation coefficient (rs), p-value, and number of repetitions (n).

| Spearman's correlation between penetration rate and depth | | | | | |
|---|---|---|---|---|---|
| | **ALT** | **SAK STAR WT** | **SAK STAR FRIDA** | **SAK THR174** | **SAK SY155** |
| Spearman's rs | 0.03 | -1.0 | -1.0 | -1.0 | -1.0 |
| n | 23 | 19 | 19 | 23 | 20 |
| p-value | 0.683 | 0.017 | 0.017 | 0.017 | 0.017 |

**Table S8** Penetration assay: descriptive statistics and p-values for thrombolytic agent penetration capability expressed as penetration rates. Experiments were conducted at 37 °C for 180 minutes in PBS, with 19-23 repetitions per condition. All agents were used at the concentration of 0.022 μM. **A)** Descriptive statistics for each group (mean ± SD, n = number of replicates). **B)** Pairwise p-values from Kruskal-Wallis test with post-hoc Dunn's correction. Statistical significance was set at p<0.05.

| **A) Penetration rates at individual distances: Descriptive statistics** | | | | | |
|---|---|---|---|---|---|
| | **ALT** | **SAK STAR WT** | **SAK STAR FRIDA** | **SAK THR174** | **SAK SY155** |
| **0.25 mm** | | | | | |
| Mean [RFU/min] | 2864 | 4287 | 4481 | 3251 | 6444 |
| SD [RFU/min] | 741 | 974 | 1194 | 1079 | 1834 |
| Median [RFU/min] | 2663 | 4487 | 4722 | 3241 | 6208 |
| Count | 23 | 19 | 19 | 23 | 20 |
| **0.50 mm** | | | | | |
| Mean [RFU/min] | 3120 | 3877 | 4069 | 2960 | 5439 |
| SD [RFU/min] | 860 | 857 | 1125 | 1018 | 1651 |
| Median [RFU/min] | 2833 | 4169 | 4381 | 3122 | 5374 |
| Count | 23 | 19 | 19 | 23 | 20 |
| **0.75 mm** | | | | | |
| Mean [RFU/min] | 3247 | 3410 | 3562 | 2599 | 4620 |
| SD [RFU/min] | 826 | 809 | 1004 | 953 | 1492 |
| Median [RFU/min] | 3018 | 3772 | 3798 | 2820 | 4481 |
| Count | 23 | 19 | 19 | 23 | 20 |
| **1.00 mm** | | | | | |
| Mean [RFU/min] | 3178 | 2942 | 3118 | 2245 | 3973 |
| SD [RFU/min] | 759 | 747 | 910 | 879 | 1332 |
| Median [RFU/min] | 3284 | 3181 | 3471 | 2511 | 3907 |
| Count | 23 | 19 | 19 | 23 | 20 |
| **1.25 mm** | | | | | |
| Mean [RFU/min] | 2899 | 2429 | 2625 | 1856 | 3362 |
| SD [RFU/min] | 645 | 638 | 785 | 742 | 1191 |
| Median [RFU/min] | 2965 | 2587 | 2830 | 2037 | 3241 |
| Count | 23 | 19 | 19 | 23 | 20 |

| **B) Penetration rates at individual distances: p-values** | | | | | |
|---|---|---|---|---|---|
| | **ALT** | **SAK STAR WT** | **SAK STAR FRIDA** | **SAK THR174** | **SAK SY155** |
| **0.25 mm** | | | | | |
| ALT | - | 0.002 | <0.001 | 0.814 | <0.001 |
| SAK STAR WT | - | - | 0.988 | 0.051 | <0.001 |
| SAK STAR FRIDA | - | - | - | 0.012 | <0.001 |
| SAK THR174 | - | - | - | - | <0.001 |
| **0.50 mm** | | | | | |
| ALT | - | 0.305 | 0.050 | >0.999 | <0.001 |
| SAK STAR WT | - | - | >0.999 | 0.170 | 0.063 |
| SAK STAR FRIDA | - | - | - | 0.025 | 0.347 |
| SAK THR174 | - | - | - | - | <0.0001 |
| **0.75 mm** | | | | | |
| ALT | - | >0.999 | >0.999 | 0.785 | 0.010 |
| SAK STAR WT | - | - | >0.999 | 0.220 | 0.107 |
| SAK STAR FRIDA | - | - | - | 0.028 | 0.604 |
| SAK THR174 | - | - | - | - | <0.001 |
| **1.00 mm** | | | | | |
| ALT | - | >0.999 | >0.999 | 0.027 | 0.431 |
| SAK STAR WT | - | - | >0.999 | 0.299 | 0.094 |
| SAK STAR FRIDA | - | - | - | 0.024 | 0.789 |
| SAK THR174 | - | - | - | - | <0.001 |
| **1.25 mm** | | | | | |
| ALT | - | 0.802 | >0.999 | 0.001 | >0.999 |
| SAK STAR WT | - | - | >0.999 | 0.450 | 0.060 |
| SAK STAR FRIDA | - | - | - | 0.025 | 0.783 |
| SAK THR174 | - | - | - | - | <0.001 |

**Table S9** Penetration assay: descriptive statistics and p-values for thrombolytic agent penetration capability expressed as the time taken for successful penetration. Experiments were conducted at 37 °C for 180 minutes in PBS, with 19-23 repetitions per condition. All agents were used at the concentration of 0.022 µM. **A)** Descriptive statistics for each group (mean ± SD, n = number of replicates). **B)** Pairwise p-values from Kruskal-Wallis test with post-hoc Dunn's correction. Statistical significance was set at p<0.05.

| **A) Penetration time at individual distances: Descriptive statistics** | | | | | |
|---|---|---|---|---|---|
| | **ALT** | **SAK STAR WT** | **SAK STAR FRIDA** | **SAK THR174** | **SAK SY155** |
| **0.25 mm** | | | | | |
| Mean [min] | 134 | 118 | 104 | 132 | 75 |
| SD [min] | 41 | 32 | 38 | 40 | 36 |
| Median [min] | 135 | 105 | 105 | 115 | 70 |
| Count | 23 | 19 | 19 | 23 | 20 |
| **0.50 mm** | | | | | |
| Mean [min] | 132 | 124 | 120 | 147 | 93 |
| SD [min] | 43 | 29 | 34 | 35 | 39 |
| Median [min] | 145 | 115 | 115 | 160 | 85 |
| Count | 23 | 19 | 19 | 23 | 20 |
| **0.75 mm** | | | | | |
| Mean [min] | 135 | 146 | 138 | 166 | 111 |
| SD [min] | 38 | 29 | 32 | 28 | 36 |
| Median [min] | 145 | 140 | 140 | 180 | 100 |
| Count | 23 | 19 | 19 | 23 | 20 |
| **1.00 mm** | | | | | |
| Mean [min] | 147 | 165 | 156 | 170 | 127 |
| SD [min] | 34 | 17 | 27 | 24 | 33 |
| Median [min] | 150 | 170 | 160 | 180 | 115 |
| Count | 23 | 19 | 19 | 23 | 20 |
| **1.25 mm** | | | | | |
| Mean [min] | 162 | 177 | 171 | 174 | 143 |
| SD [min] | 26 | 9 | 14 | 19 | 30 |
| Median [min] | 180 | 180 | 180 | 180 | 150 |
| Count | 23 | 19 | 19 | 23 | 20 |

| **B) Penetration time at individual distances: p-values** | | | | | |
|---|---|---|---|---|---|
| | **ALT** | **SAK STAR WT** | **SAK STAR FRIDA** | **SAK THR174** | **SAK SY155** |
| **0.25 mm** | | | | | |
| ALT | - | >0.999 | 0.186 | >0.999 | <0.001 |
| SAK STAR WT | - | - | >0.999 | >0.999 | 0.048 |
| SAK STAR FRIDA | - | - | - | 0.263 | 0.584 |
| SAK THR174 | - | - | - | - | <0.001 |
| **0.50 mm** | | | | | |
| ALT | - | >0.999 | >0.999 | >0.999 | 0.015 |
| SAK STAR WT | - | - | >0.999 | 0.539 | 0.131 |
| SAK STAR FRIDA | - | - | - | 0.310 | 0.238 |
| SAK THR174 | - | - | - | - | <0.001 |
| **0.75 mm** | | | | | |
| ALT | - | >0.999 | >0.999 | 0.041 | 0.355 |
| SAK STAR WT | - | - | >0.999 | 0.585 | 0.048 |
| SAK STAR FRIDA | - | - | - | 0.097 | 0.320 |
| SAK THR174 | - | - | - | - | <0.001 |
| **1.00 mm** | | | | | |
| ALT | - | >0.999 | >0.999 | 0.045 | 0.565 |
| SAK STAR WT | - | - | >0.999 | >0.999 | 0.012 |
| SAK STAR FRIDA | - | - | - | 0.710 | 0.072 |
| SAK THR174 | - | - | - | - | <0.001 |
| **1.25 mm** | | | | | |
| ALT | - | 0.607 | >0.999 | 0.583 | 0.078 |
| SAK STAR WT | - | - | >0.999 | >0.999 | <0.001 |
| SAK STAR FRIDA | - | - | - | >0.999 | 0.007 |
| SAK THR174 | - | - | - | - | <0.001 |

**Table S10** Penetration assay: descriptive statistics and p-values for thrombolytic agent penetration capability expressed as penetration success frequency. Experiments were conducted at 37 °C for 180 minutes in PBS, with 19-23 repetitions per condition. All agents were used at the concentration of 0.022 μM. **A)** Descriptive statistics for each thrombolytic agent. **B)** Pairwise p-values from Fisher's exact test with post-hoc Holm-Bonferroni correction. Statistical significance was set at $p<0.05$.

| **A) Penetration success frequency: Descriptive statistics** | | | | | |
|---|---|---|---|---|---|
| | **ALT** | **SAK STAR WT** | **SAK STAR FRIDA** | **SAK THR174** | **SAK SY155** |
| **0.25 mm** | | | | | |
| Penetration | 16 (70%) | 16 (84%) | 18 (95%) | 17 (74%) | 20 (100%) |
| No penetration | 7 (30%) | 3 (16%) | 1 (5%) | 6 (26%) | 0 (0%) |
| Count (100%) | 23 | 19 | 19 | 23 | 20 |
| **0.50 mm** | | | | | |
| Penetration | 18 (78%) | 16 (84%) | 16 (84%) | 15 (65%) | 19 (95%) |
| No penetration | 5 (22%) | 3 (16%) | 3 (16%) | 8 (35%) | 1 (5%) |
| Count (100%) | 23 | 19 | 19 | 23 | 20 |
| **0.75 mm** | | | | | |
| Penetration | 18 (78%) | 13 (68%) | 14 (74%) | 7 (30%) | 17 (85%) |
| No penetration | 5 (22%) | 6 (32%) | 5 (26%) | 16 (70%) | 3 (15%) |
| Count (100%) | 23 | 19 | 19 | 23 | 20 |
| **1.00 mm** | | | | | |
| Penetration | 15 (65%) | 10 (53%) | 10 (53%) | 4 (17%) | 17 (85%) |
| No penetration | 8 (35%) | 9 (47%) | 9 (47%) | 19 (83%) | 3 (15%) |
| Count (100%) | 23 | 19 | 19 | 23 | 20 |
| **1.25 mm** | | | | | |
| Penetration | 9 (39%) | 10 (53%) | 10 (53%) | 4 (17%) | 17 (85%) |
| No penetration | 14 (61%) | 9 (47%) | 9 (47%) | 19 (83%) | 3 (15%) |
| Count (100%) | 23 | 19 | 19 | 23 | 20 |

| **B) Penetration frequency in individual distances: p-values** | | | | | |
|---|---|---|---|---|---|
| | **ALT** | **SAK STAR WT** | **SAK STAR FRIDA** | **SAK THR174** | **SAK SY155** |
| **0.25 mm** | | | | | |
| ALT | - | >0.999 | 0.433 | >0.999 | 0.100 |
| SAK STAR WT | - | - | >0.999 | >0.999 | 0.736 |
| SAK STAR FRIDA | - | - | - | 0.736 | >0.999 |
| SAK THR174 | - | - | - | - | 0.206 |
| **0.50 mm** | | | | | |
| ALT | - | >0.999 | >0.999 | >0.999 | >0.999 |
| SAK STAR WT | - | - | >0.999 | >0.999 | >0.999 |
| SAK STAR FRIDA | - | - | - | >0.999 | >0.999 |
| SAK THR174 | - | - | - | - | 0.243 |
| **0.75 mm** | | | | | |
| ALT | - | >0.999 | >0.999 | 0.024 | >0.999 |
| SAK STAR WT | - | - | >0.999 | 0.201 | >0.999 |
| SAK STAR FRIDA | - | - | - | 0.097 | >0.999 |
| SAK THR174 | - | - | - | - | 0.006 |
| **1.00 mm** | | | | | |
| ALT | - | >0.999 | >0.999 | 0.021 | 0.701 |
| SAK STAR WT | - | - | >0.999 | 0.184 | 0.244 |
| SAK STAR FRIDA | - | - | - | 0.184 | 0.244 |
| SAK THR174 | - | - | - | - | <0.001 |
| **1.25 mm** | | | | | |
| ALT | - | >0.999 | >0.999 | 0.758 | 0.038 |
| SAK STAR WT | - | - | >0.999 | 0.184 | 0.244 |
| SAK STAR FRIDA | - | - | - | 0.184 | 0.244 |
| SAK THR174 | - | - | - | - | <0.001 |

**Table S11** SAK Variants Overview - Wild-type staphylokinases and low-immunogenic staphylokinases that were compared in this study. Production yields are expressed in miligrams per liter of culture [mg / L] of all five studied proteins after cultivation and full purification procedure. Proteins were produced in *E.coli* BL21(DE3) cells for 4 hours at 37 °C and purified by two rounds of immobilized metal affinity chromatography and one run of size-exclusion chromatography with a cleavage step during dialysis

| **Protein overview part 1** | | | | |
|---|---|---|---|---|
| **Protein** | **Mutations** | **Source** | **Yield** | **Reference** |
| Staphylokinase SAK STAR WT | | *Staphylococcus aureus* genome | 60 mg/L | [31] |
| Staphylokinase SAK 42D WT | S34G, G36R, H43R | Bacteriophage genome | 10 mg/L | [31] |
| Staphylokinase SAK STAR FRIDA | K74A, E75A, R77A | Russian “FRIDA” Clinical Trial | 40 mg/L | [14] |
| Staphylokinase SAK THR174 | E65D, K74Q, R77E, E80S, D82S, V112T, K130Y, E134R | Desiré Collen' group / ThromboGenics | 20 mg/L | [30] |
| Staphylokinase SAK SY155 | K35A, E65Q, K74R, D82A, S84A, T90A, E99D, T101S, E108A, K109A, K130T, K135R | Desiré Collen' group | 25 mg/L | [29] |

| **Protein overview part 2** | | |
|---|---|---|
| **Protein** | **Molecular Weight** | **Sequence** |
| Staphylokinase SAK STAR WT | 15.49 kDa | SSSFDKGKYKKGDDASYFEPTGPYLMVNVTGVDSKGNELLSPHYVEFPI KPGTTLTKEKIEYYVEWALDATAYKEFRVVELDPSAKIEVTYYDKNKKKEE TKSFPITEKGFVVPDLSEHIKNPGFNLITKVVIEKK |
| Staphylokinase SAK 42D WT | 15.58 kDa | SSSFDKGKYKKGDDASYFEPTGPYLMVNVTGVDGKRNELLSPRYVEFPI KPGTTLTKEKIEYYVEWALDATAYKEFRVVELDPSAKIEVTYYDKNKKKEE TKSFPITEKGFVVPDLSEHIKNPGFNLITKVVIEKK |
| Staphylokinase SAK STAR FRIDA | 15.29 kDa | SSSFDKGKYKKGDDASYFEPTGPYLMVNVTGVDSKGNELLSPHYVEFPI KPGTTLTKEKIEYYVEWALDATAYAAFAVVELDPSAKIEVTYYDKNKKKEET KSFPITEKGFVVPDLSEHIKNPGFNLITKVVIEKK |
| Staphylokinase SAK THR174 | 15.45 kDa | SSSFDKGKYKKGDDASYFEPTGPYLMVNVTGVDSKGNELLSPHYVEFPI KPGTTLTKEKIEYYVDWALDATAYQEFEVVSLSPSAKIEVTYYDKNKKKEE TKSFPITEKGFTVPDLSEHIKNPGFNLITYVVIRKK |
| Staphylokinase SAK SY155 | 15.23 kDa | SSSFDKGKYKKGDDASYFEPTGPYLMVNVTGVDSAGNELLSPHYVEFPI KPGTTLTKEKIEYYVQWALDATAYREFRVVELAPAAKIEVAYYDKNKKKDE SKSFPITAAGFVVPDLSEHIKNPGFNLITTVVIERK |

Expressed sequence (SAK STAR WT):

MHHHHHHENLYFQSSSFDKGKYKKGDDASYFEPTGPYLMVNVTGVDSKGNELLSPHYVEF PIKPGTTLTKEKIEYYVEWALDATAYKEFRVVELDPSAKIEVTYYDKNKKKEETKSFPITEKGF VVPDLSEHIKNPGFNLITKVVIEKK

Final sequence (SAK STAR WT) after TEV cleavage:

SSSFDKGKYKKGDDASYFEPTGPYLMVNVTGVDSKGNELLSPHYVEFPIKPGTTLTKEKIEY YVEWALDATAYKEFRVVELDPSAKIEVTYYDKNKKKEETKSFPITEKGFVVPDLSEHIKNPGF NLITKVVIEKK

To prevent confusion when the mutations are introduced to the sequence, a uniform numbering is suggested. The amino acids are counted from the first triplet of serines (marked blue as S1, S2, and S3)

25 kDa
17 kDa
10 kDa

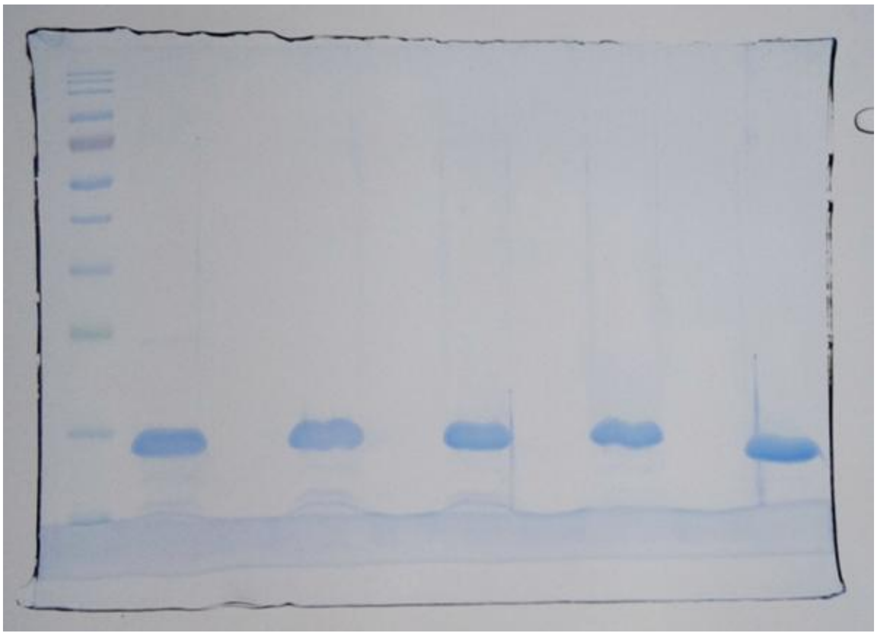

**Figure S3** SDS-PAGE gel showing the purity of SAK 42D WT, SAK STAR WT, SAK STAR FRIDA, SAK THR174, and SAK SY155, verifying their purity after the full purification cycle. 10 µL of each sample was loaded onto the gel. The sample concentration of the sample being around 1 mg/ml. The sample was prepared by mixing 20 µL of the original sample and 20 µL of Laemmli Sample buffer, incubated for 10 minutes at 95°C, and then run at 150 V, 400 mA for 60 minutes in 15% polyacrylamide gel.

25 kDa
17 kDa
10 kDa

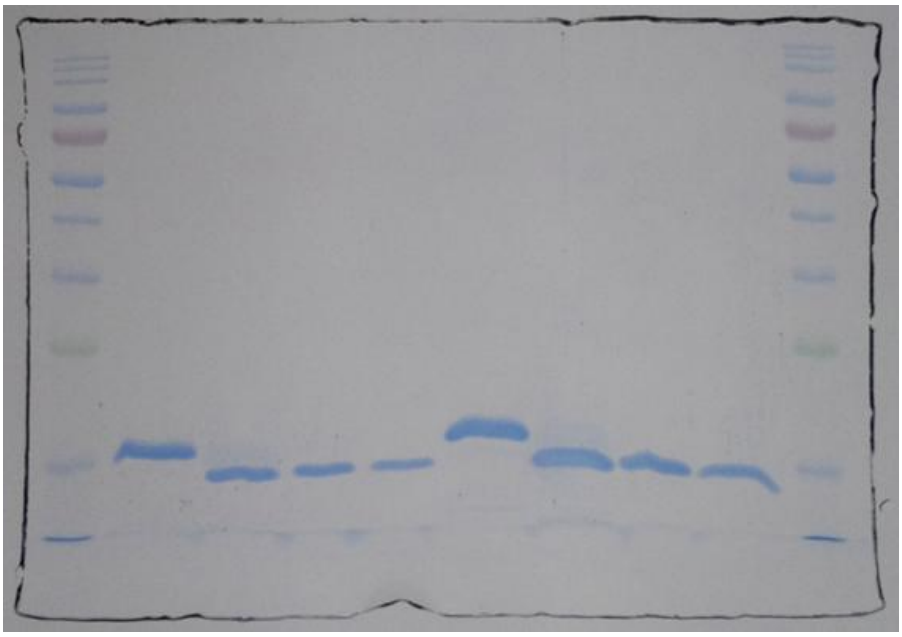

**Figure S4** SDS-PAGE gel performed on samples collected during the purification of SAK 42D WT, showing the cleavage of the HIS-TEV sequence (samples after the first metal-affinity purification step, after cleavage by TEV protease and dialysis, after the second metal-affinity purification step, and after gel permeation chromatography (SEC)). The loading was repeated twice with 8 µL and 16 µL of the sample, with a concentration of the sample being around 1 mg/ml. The sample was prepared by mixing 20 µL of the original sample and 20 µL of Laemmli Sample buffer, incubated for 10 minutes at 95°C, and then run at 150 V, 400 mA for 60 minutes in 15% polyacrylamide gel.

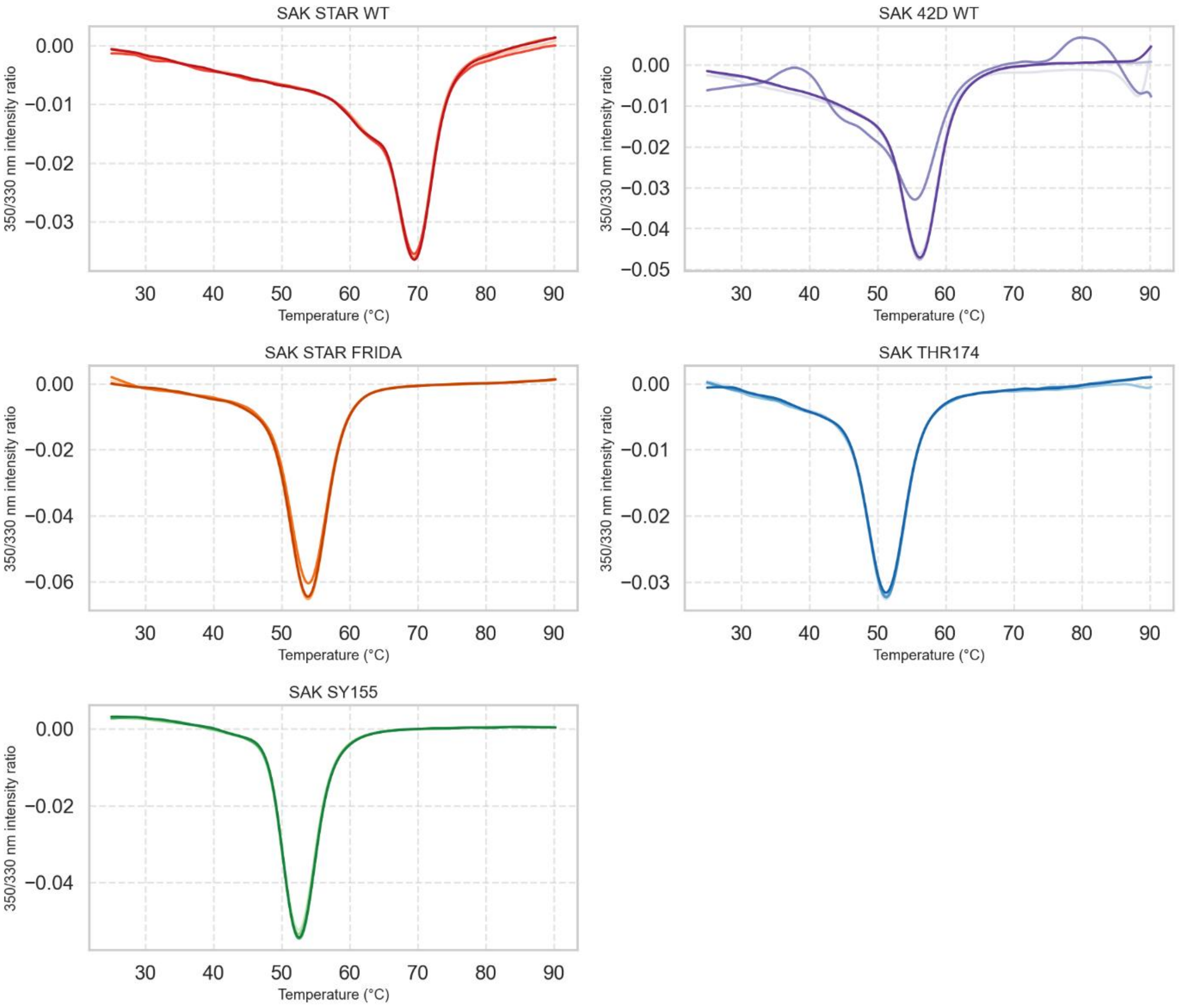


**Figure S5** Differential scanning fluorimetry (DSF) curve of thermal stability showing the first derivative of the 350/330 nm fluorescence intensity ratio for all 5 studied variants, SAK STAR WT in shades of red, SAK 42D WT in shades of violet, SAK STAR FRIDA in shades of orange, SAK THR174 in shades of blue, and SAK SY155 in shades of green. The minimum of the curve represents the melting temperature, while the initial decrease towards the minimum is the onset of the melting. Melting was measured from 25 to 90 °C with the rate of increase of 1 °C per minute, and the environment was PBS (pH=7.4).

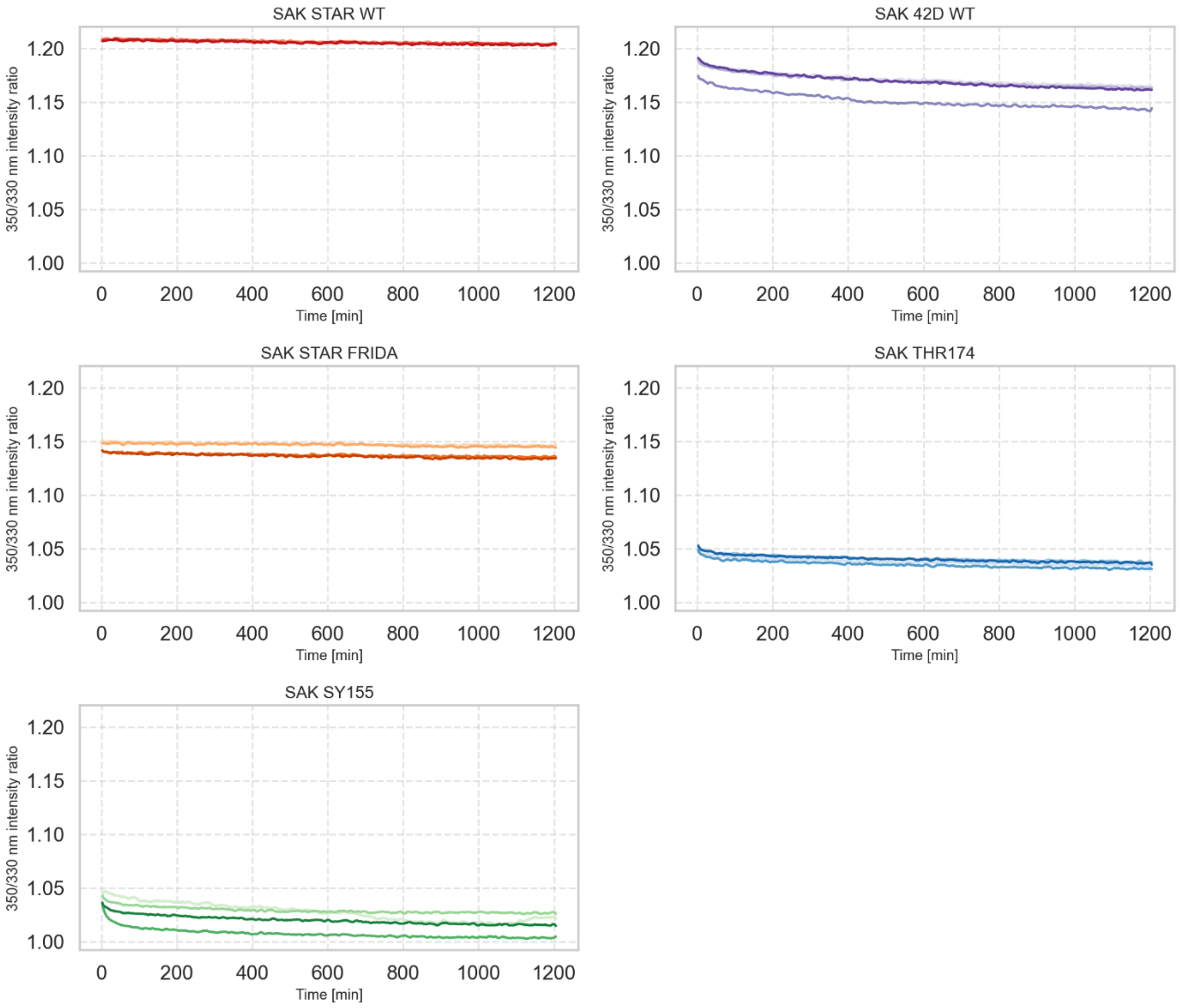


**Figure S6** Isothermal DSF measurement of protein aggregation performed at 37 °C for 20 hours in tetraplicate (n=4) in PBS (pH=7.4),  SAK STAR WT in shades of red, SAK 42D WT in shades of violet, SAK STAR FRIDA in shades of orange, SAK THR174 in shades of blue, and SAK SY155 in shades of green. Neither variant exhibits any significant changes, so we can conclude that no aggregation is happening in this time window.

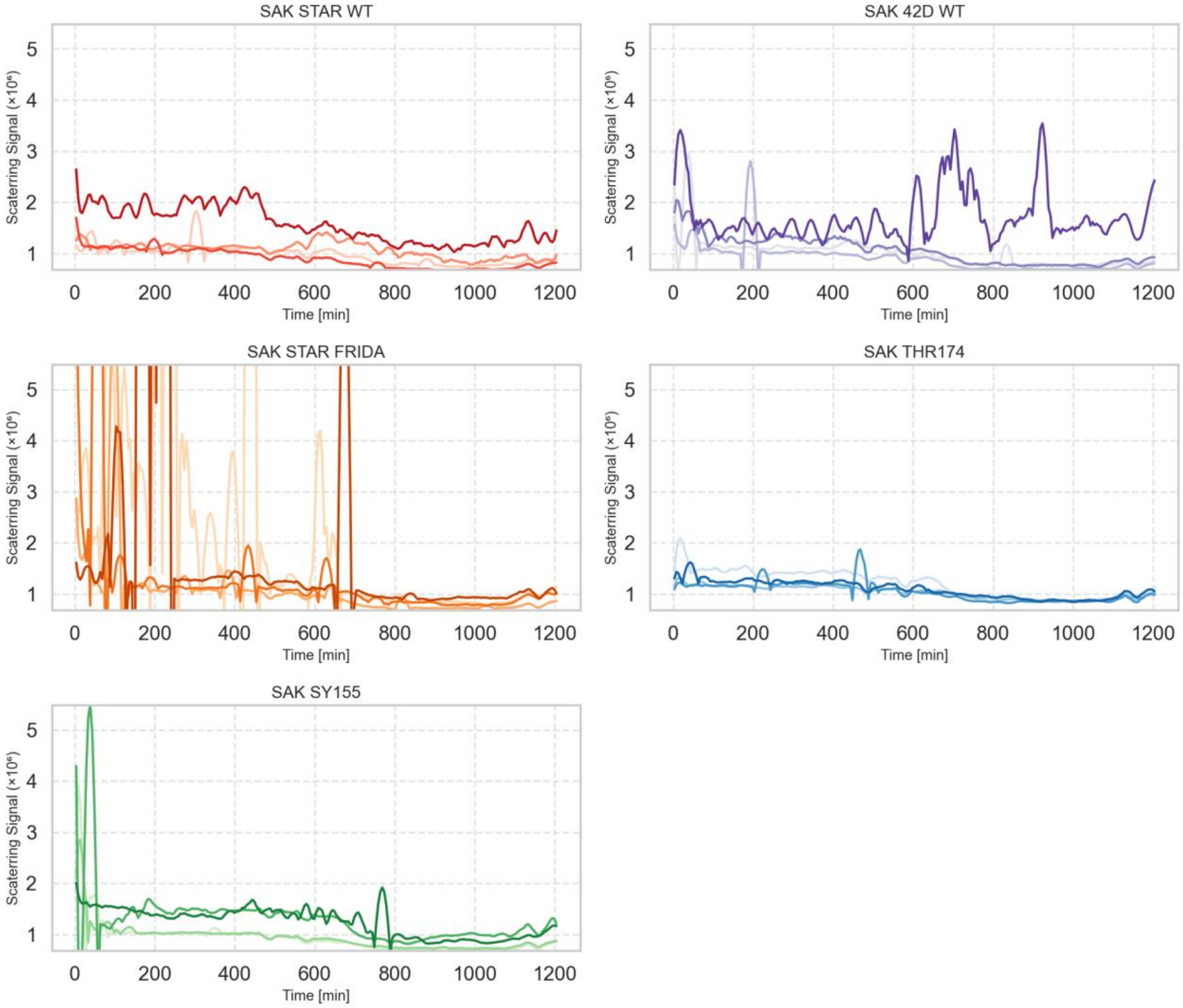


**Figure S7** Isothermal DLS measurement of protein aggregation performed at 37 °C for 20 hours in tetraplicate (n=4) in PBS (pH=7.4), SAK STAR WT in shades of red, SAK 42D WT in shades of violet, SAK STAR FRIDA in shades of orange, SAK THR174 in shades of blue, and SAK SY155 in shades of green. Some noise can be seen in the curves, especially for SAK STAR FRIDA, but the response is limited, which suggests no significant changes are happening to any of the studied variants.

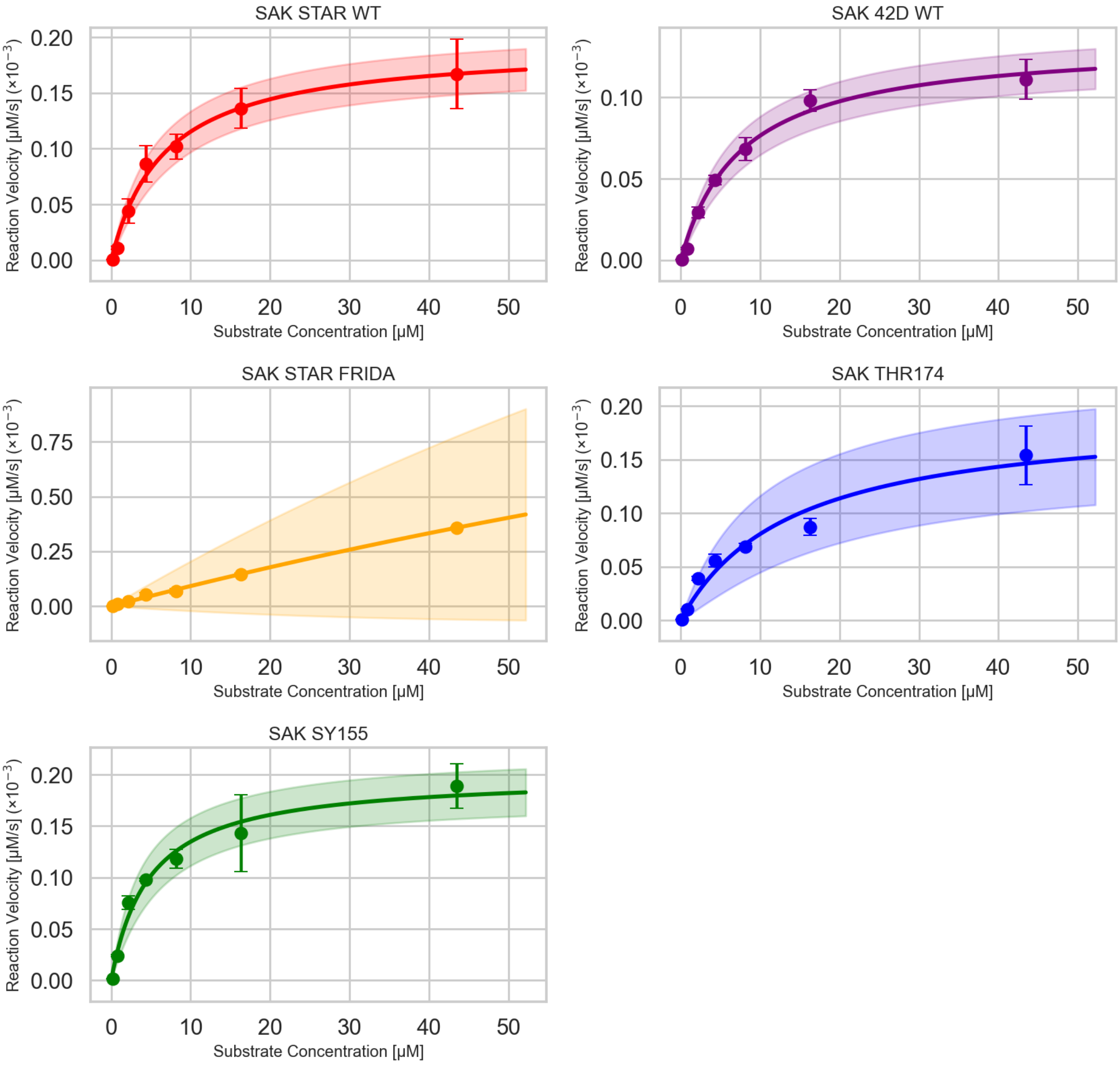


**Figure S8** The maximum reaction velocities extracted from the initial parabolic phase at different substrate concentrations. The data were fitted by the Michaelis-Menten equation to obtain the apparent maximum reaction velocity and apparent Michaelis constant for each variant. The experiment was run at 37 °C in PBS buffer (pH=7.4) with plasminogen concentrations of 0, 0.16, 0.81, 2.17, 4.35, 8.15, 16.3, 43.5 μM and staphylokinase concentration of 0.5 nM. VLK-AMC was present at the final concentration of 200 μM. For all variants, except for SAK STAR FRIDA, the fitting was satisfactory. For SAK STAR FRIDA, the kinetic parameters would have to be extrapolated, which significantly decreases their reliability, so an alternative evaluation, a linear fit, was used instead.

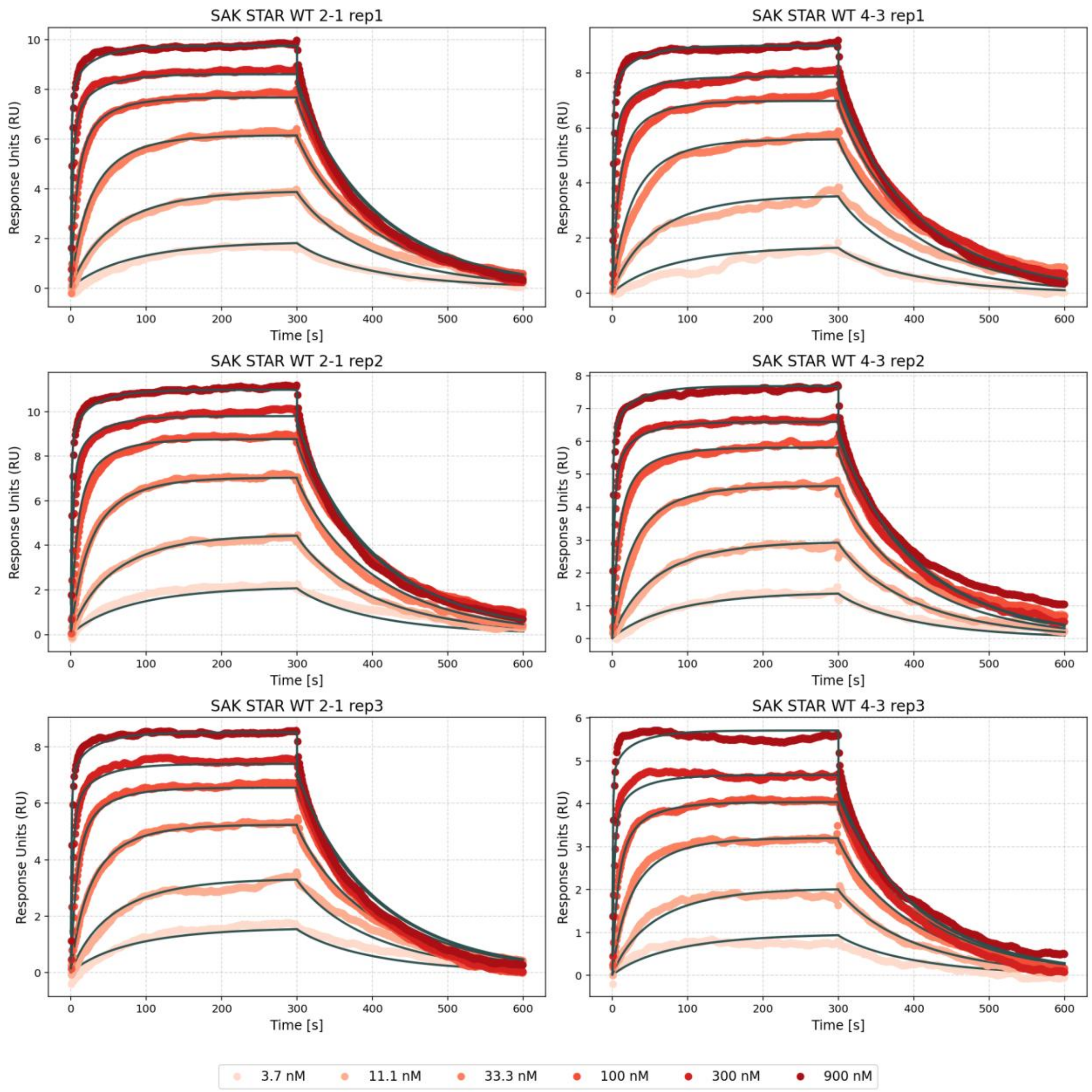


**Figure S9** The binding kinetics of SAK STAR WT to human plasmin were analysed by surface-based assays using a CM5 sensor chip. Human plasmin (~850 RU on channel 2, ~550 RU on channel 4) was immobilized by amine coupling, and SAK was injected at concentrations of 3.7, 11, 33, 100, 300, and 900 nM in PBS/Tween80 (pH 7.4) at 25 °C. Binding curves were fitted numerically to a model shown in Scheme 1 in the Methods section. Curves labelled 2-1 refer to data obtained from channels 1 and 2, while curves labelled 4-3 refer to data obtained from channels 3 and 4.

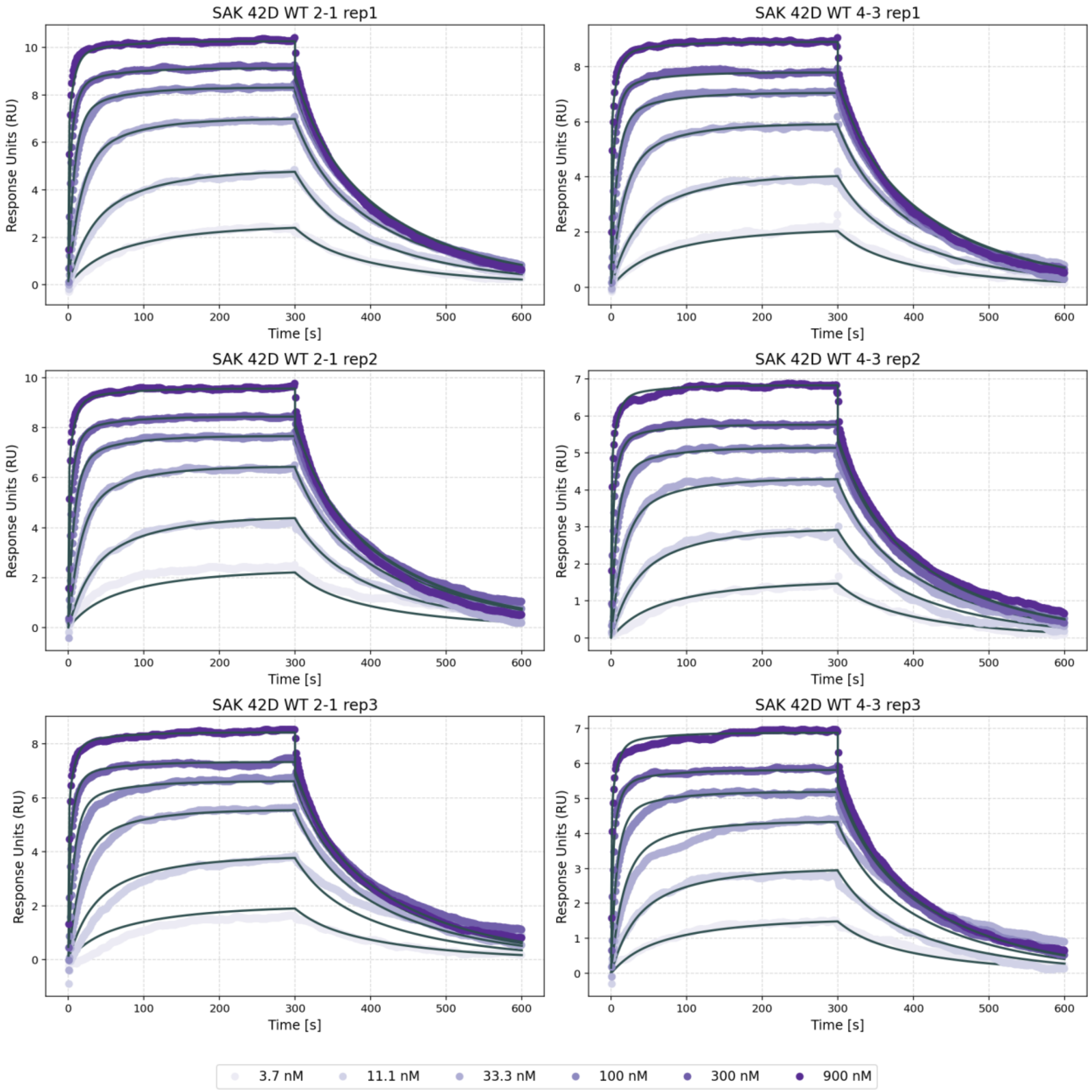


**Figure S10** The binding kinetics of SAK 42D WT to human plasmin were analysed by surface-based assays using a CM5 sensor chip. Human plasmin (~850 RU on channel 2, ~550 RU on channel 4) was immobilized by amine coupling, and SAK was injected at concentrations of 3.7, 11, 33, 100, 300, and 900 nM in PBS/Tween80 (pH 7.4) at 25 °C. Binding curves were fitted numerically to a model shown in Scheme 1 in the Methods section. Curves labelled 2-1 refer to data obtained from channels 1 and 2, while curves labelled 4-3 refer to data obtained from channels 3 and 4.

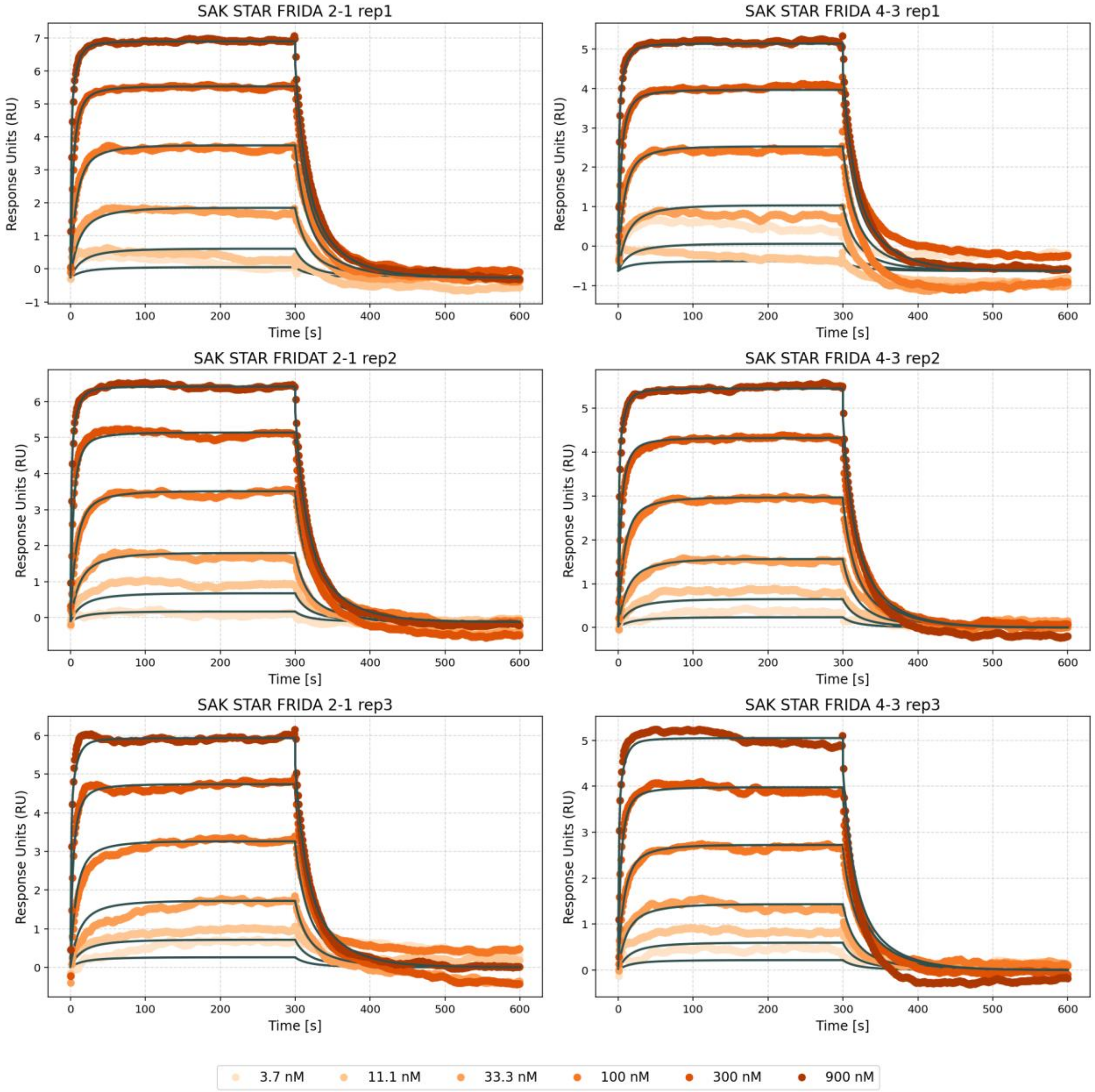


**Figure S11** The binding kinetics of SAK STAR FRIDA to human plasmin were analysed by surface-based assays using a CM5 sensor chip. Human plasmin (~850 RU on channel 2, ~550 RU on channel 4) was immobilized by amine coupling, and SAK was injected at concentrations of 3.7, 11, 33, 100, 300, and 900 nM in PBS/Tween80 (pH 7.4) at 25 °C. Binding curves were fitted numerically to a model shown in Scheme 1 in the Methods section. Curves labelled 2-1 refer to data obtained from channels 1 and 2, while curves labelled 4-3 refer to data obtained from channels 3 and 4.

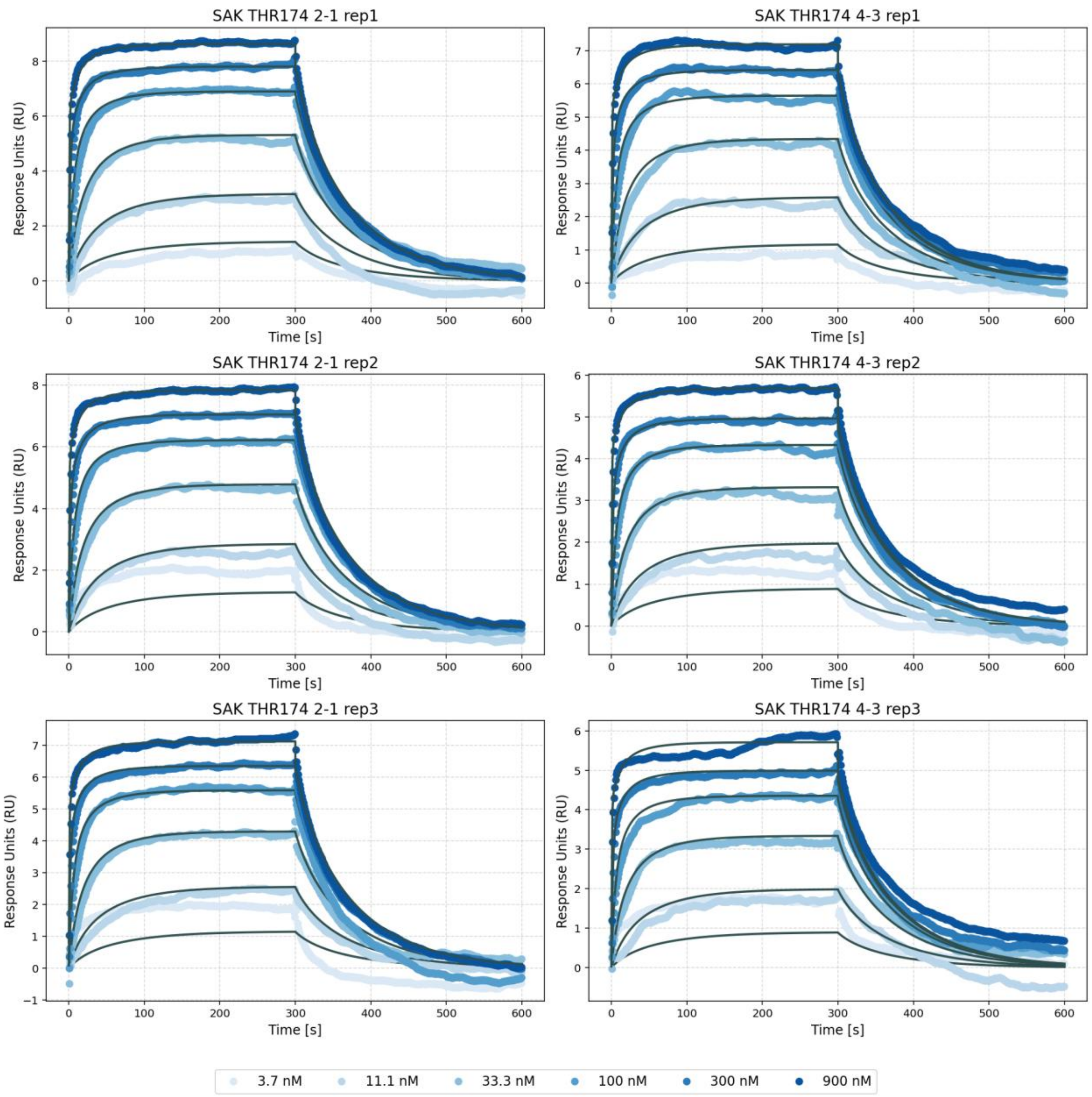


**Figure S12** The binding kinetics of SAK THR174 to human plasmin were analysed by surface-based assays using a CM5 sensor chip. Human plasmin (~850 RU on channel 2, ~550 RU on channel 4) was immobilized by amine coupling, and SAK was injected at concentrations of 3.7, 11, 33, 100, 300, and 900 nM in PBS/Tween80 (pH 7.4) at 25 °C. Binding curves were fitted numerically to a model shown in Scheme 1 in the Methods section. Curves labelled 2-1 refer to data obtained from channels 1 and 2, while curves labelled 4-3 refer to data obtained from channels 3 and 4.

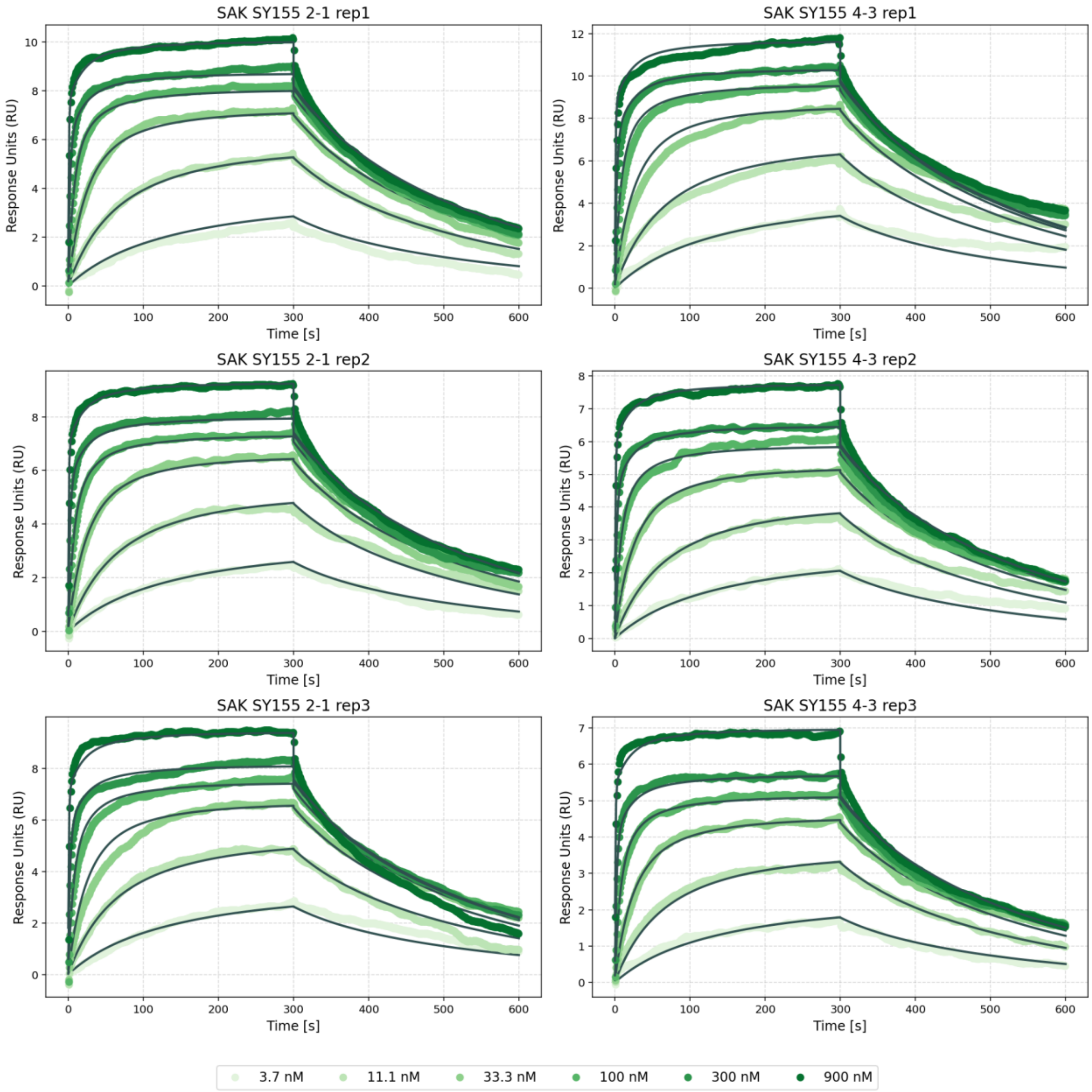


**Figure S13** The binding kinetics of SAK SY155 to human plasmin were analysed by surface-based assays using a CM5 sensor chip. Human plasmin (~850 RU on channel 2, ~550 RU on channel 4) was immobilized by amine coupling, and SAK was injected at concentrations of 3.7, 11, 33, 100, 300, and 900 nM in PBS/Tween80 (pH 7.4) at 25 °C. Binding curves were fitted numerically to a model shown in Scheme 1 in the Methods section. Curves labelled 2-1 refer to data obtained from channels 1 and 2, while curves labelled 4-3 refer to data obtained from channels 3 and 4.

1

MYO 9/95-22

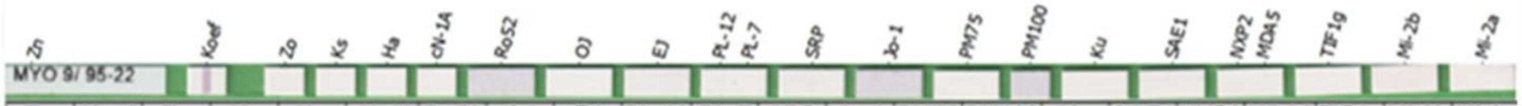


| Zn | Koef | Zo | Ks | Ha | cN-1A | Ro52 | OJ | EJ | PL-12 | PL-7 | SRP | Jo-1 | PM75 | PM100 | Ku | SAE1 | NXP2 | MDA5 | TIF1g | Mi-2b | Mi-2a |
|---|---|---|---|---|---|---|---|---|---|---|---|---|---|---|---|---|---|---|---|---|---|
| -1 | 39 | 1 | 0 | 0 | 1 | 1 | 1 | 1 | 2 | 0 | 1 | 0 | 1 | 0 | 1 | 0 | 1 | 1 | 1 | 1 | 0 |
| o | ++ | o | o | o | o | o | o | o | o | o | o | o | o | o | o | o | o | o | o | o | o |

2

MYO 9/95-23

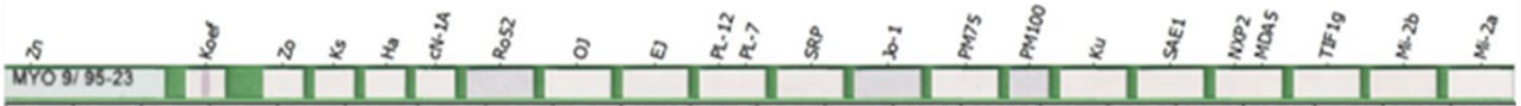


| Zn | Koef | Zo | Ks | Ha | cN-1A | Ro52 | OJ | EJ | PL-12 | PL-7 | SRP | Jo-1 | PM75 | PM100 | Ku | SAE1 | NXP2 | MDA5 | TIF1g | Mi-2b | Mi-2a |
|---|---|---|---|---|---|---|---|---|---|---|---|---|---|---|---|---|---|---|---|---|---|
| -1 | 34 | 1 | 2 | 0 | 1 | 0 | 1 | 0 | 2 | 1 | 1 | 1 | 0 | 1 | 1 | 0 | 2 | 1 | 0 | 1 | 0 |
| o | ++ | o | o | o | o | o | o | o | o | o | o | o | o | o | o | o | o | o | o | o | o |

3

MYO 9/95-24

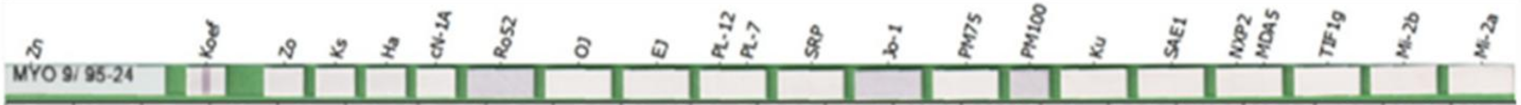


| Zn | Koef | Zo | Ks | Ha | cN-1A | Ro52 | OJ | EJ | PL-12 | PL-7 | SRP | Jo-1 | PM75 | PM100 | Ku | SAE1 | NXP2 | MDA5 | TIF1g | Mi-2b | Mi-2a |
|---|---|---|---|---|---|---|---|---|---|---|---|---|---|---|---|---|---|---|---|---|---|
| -1 | 50 | 1 | 1 | 1 | 1 | 2 | 0 | 1 | 1 | 0 | 1 | 0 | 1 | 2 | 1 | 0 | 2 | 1 | 0 | 1 | 0 |
| o | ++ | o | o | o | o | o | o | o | o | o | o | o | o | o | o | o | o | o | o | o | o |

4

MYO 9/95-25

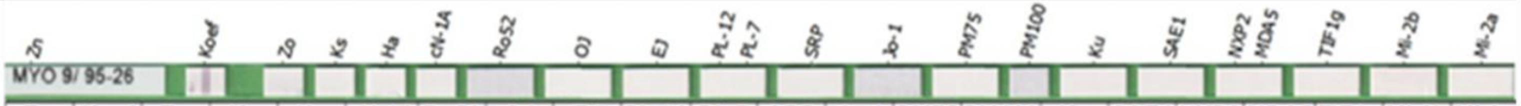


| Zn | Koef | Zo | Ks | Ha | cN-1A | Ro52 | OJ | EJ | PL-12 | PL-7 | SRP | Jo-1 | PM75 | PM100 | Ku | SAE1 | NXP2 | MDA5 | TIF1g | Mi-2b | Mi-2a |
|---|---|---|---|---|---|---|---|---|---|---|---|---|---|---|---|---|---|---|---|---|---|
| -1 | 42 | 1 | 1 | 1 | 0 | 1 | 0 | 0 | 1 | 1 | 0 | 2 | 1 | 0 | 1 | 0 | 1 | 0 | 0 | 1 | 0 |
| o | ++ | o | o | o | o | o | o | o | o | o | o | o | o | o | o | o | o | o | o | o | o |

**Figure S14** Reactivity of mice sera with myositis-associated/specific human autoantigens. 1 - 5 pooled sera of mice vaccinated with individual SAK versions with adjuvant. 1 - SAK THR174, 2 - SAK SY155, 3 - SAK STAR FRIDA, 4 – SAK STAR WT. Autoantibodies against the following autoantigens were tested: Mi-2a, Mi-2b, TIF1g (transcriptional intermediary factor 1) MDA5 (melanoma differentiation-associated protein 5), NXP2 (nuclear matrix protein 2), SAE1 (small ubiquitin-like modifier 1-activating enzyme), Ku, PM-Scl100 (polymyositis/scleroderma), PM-Scl75, Jo-1 (histidyl-tRNA synthetase), SRP (signal recognition particle), PL-7 (threonyl-tRNA synthetase), PL-12 (alanyltRNA synthetase), EJ (glycyl-tRNA synthetase), OJ (isoleucyl-tRNA synthetase), Ro-52 (TRIM21).

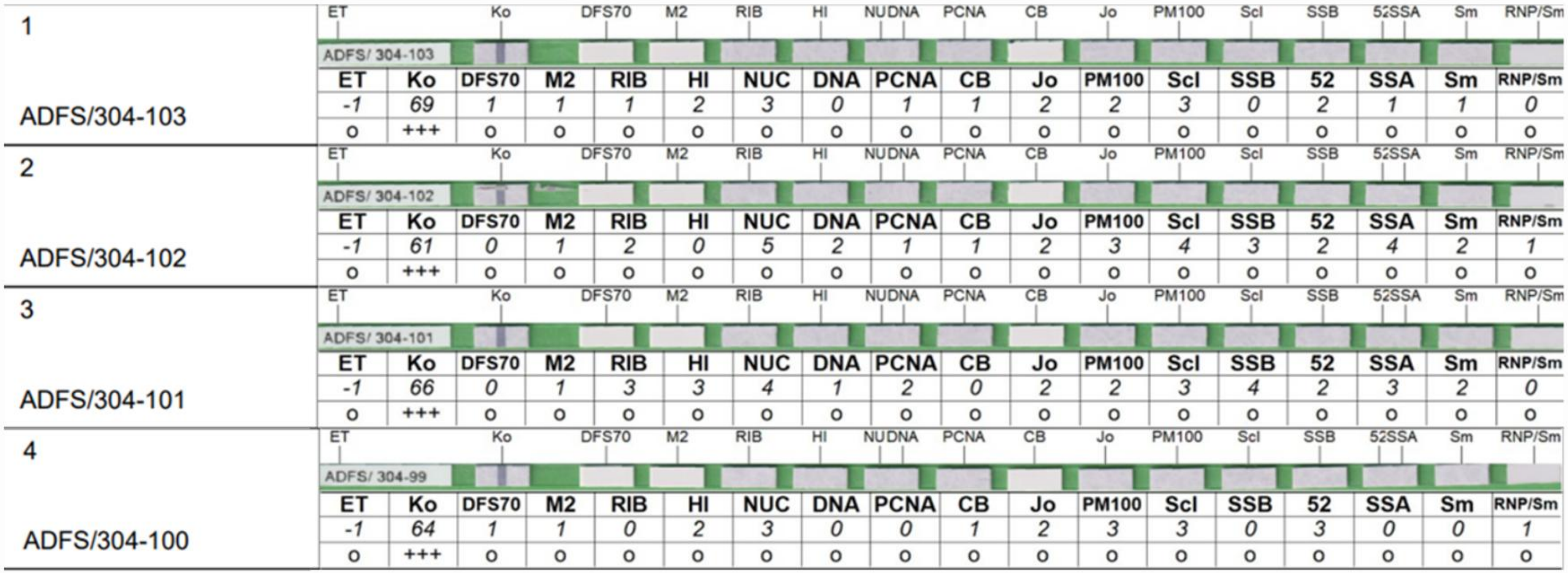

| Sample | | ET | Ko | DFS70 | M2 | RIB | HI | NUC | DNA | PCNA | CB | Jo | PM100 | Scl | SSB | 52 | SSA | Sm | RNP/Sm |
|---|---|---|---|---|---|---|---|---|---|---|---|---|---|---|---|---|---|---|---|
| 1 ADFS/304-103 | | -1 | 69 | 1 | 1 | 1 | 2 | 3 | 0 | 1 | 1 | 2 | 2 | 3 | 0 | 2 | 1 | 1 | 0 |
| | | o | +++ | o | o | o | o | o | o | o | o | o | o | o | o | o | o | o | o |
| 2 ADFS/304-102 | | -1 | 61 | 0 | 1 | 2 | 0 | 5 | 2 | 1 | 1 | 2 | 3 | 4 | 3 | 2 | 4 | 2 | 1 |
| | | o | +++ | o | o | o | o | o | o | o | o | o | o | o | o | o | o | o | o |
| 3 ADFS/304-101 | | -1 | 66 | 0 | 1 | 3 | 3 | 4 | 1 | 2 | 0 | 2 | 2 | 3 | 4 | 2 | 3 | 2 | 0 |
| | | o | +++ | o | o | o | o | o | o | o | o | o | o | o | o | o | o | o | o |
| 4 ADFS/304-100 | | -1 | 64 | 1 | 1 | 0 | 2 | 3 | 0 | 0 | 1 | 2 | 3 | 3 | 0 | 3 | 0 | 0 | 1 |
| | | o | +++ | o | o | o | o | o | o | o | o | o | o | o | o | o | o | o | o |

**Figure S15** Reactivity of murine sera with nuclear human autoantigens. 1 - 5 pooled sera of mice vaccinated with individual SAK versions with adjuvant. 1 - SAK THR174, 2 - SAK SY155, 3 - SAK STAR FRIDA, 4 SAK STAR WT. Autoantibodies against the following autoantigens were tested: nRNP/Sm, Sm (Smith), SS-A/Ro, Ro-52 (TRIM21), SS-B/La, Scl-70 (topoisomerase I), PM-Scl (polymyositis/scleroderma), Jo-1 (histidyl-tRNA synthetase), CENP B (centromere protein B), PCNA (proliferating cell nuclear antigen), dsDNA, nucleosomes, histones, ribosomal proteins, AMA M2, DFS70 (dense fine speckled, 70kDa molecular weight protein).

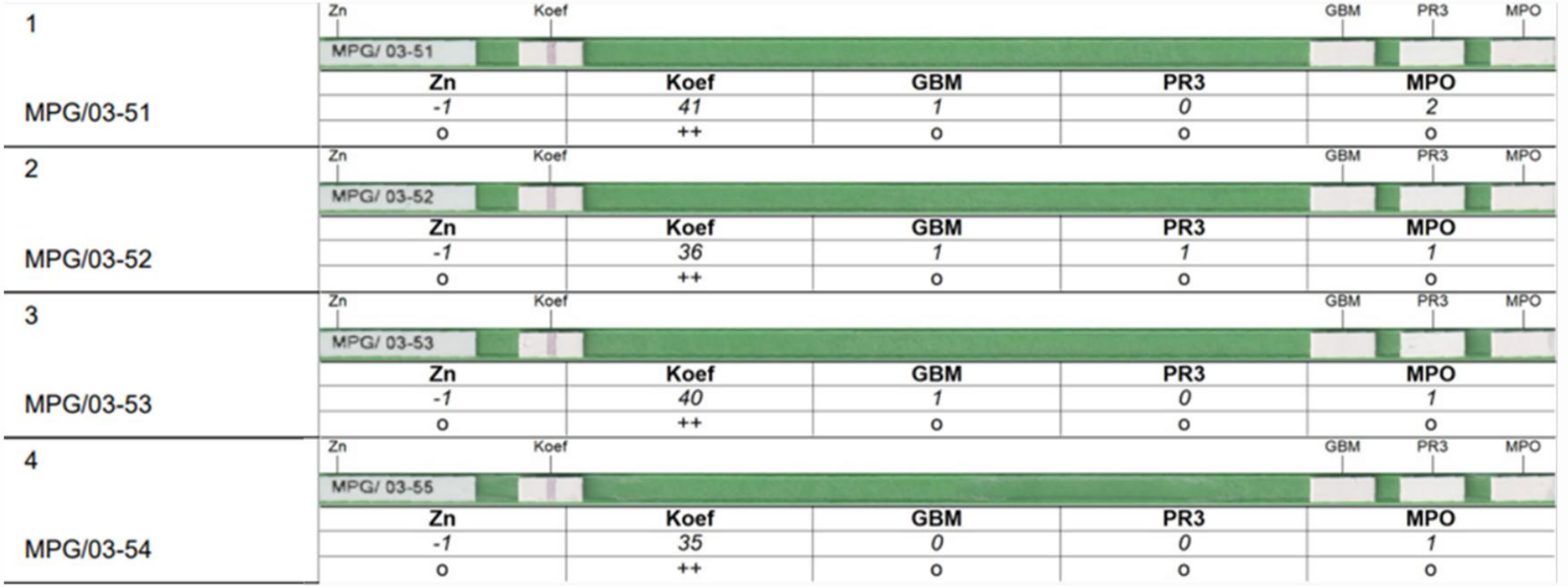


| | | Zn | Koef | GBM | PR3 | MPO |
|---|---|---|---|---|---|---|
| 1 | MPG/03-51 | -1 | 41 | 1 | 0 | 2 |
| | | o | ++ | o | o | o |
| 2 | MPG/03-52 | -1 | 36 | 1 | 1 | 1 |
| | | o | ++ | o | o | o |
| 3 | MPG/03-53 | -1 | 40 | 1 | 0 | 1 |
| | | o | ++ | o | o | o |
| 4 | MPG/03-54 | -1 | 35 | 0 | 0 | 1 |
| | | o | ++ | o | o | o |

**Figure S16** Reactivity of murine sera with MPO, PR3, and GBM human autoantigens. 1 - 5 pooled sera of mice vaccinated with individual SAK versions with adjuvants. 1 - SAK THR174, 2 - SAK SY155, 3 - SAK STAR FRIDA, 4 - SAK STAR WT. Autoantibodies against the following autoantigens were tested: GMB (glomerular basement membrane antigen - collagen alpha3(IV)), PR3 (neutrophil protease 3), and MPO (neutrophil myeloperoxidase).

*FluoroSpot Analysis*

Cells were isolated as described for the proliferation assay and analyzed using the FluoroSpot Flex Human IFN-γ/IL-2 kit (Mabtech, Sweden) according to the manufacturer's instructions. Cells were plated at a density of 1 × 105 cells per well with either medium alone (negative control), PHA (5 µg/mL; positive control), ALT (1.3 µg/mL, clinically relevant, and 13 µg/mL), or the SAK variants (1.3 and 13 µg/mL). Cells were maintained for 3 days at 37 °C in a humidified 5% $CO_2$ incubator under sterile conditions. Following incubation, secreted IFN-γ was detected using fluorophore-conjugated antibodies, and fluorescent spots were quantified using the ImmunoSpot® reader (Cellular Technology Limited, Germany).

According to internal routine criteria, spot counts below 10 were considered background.

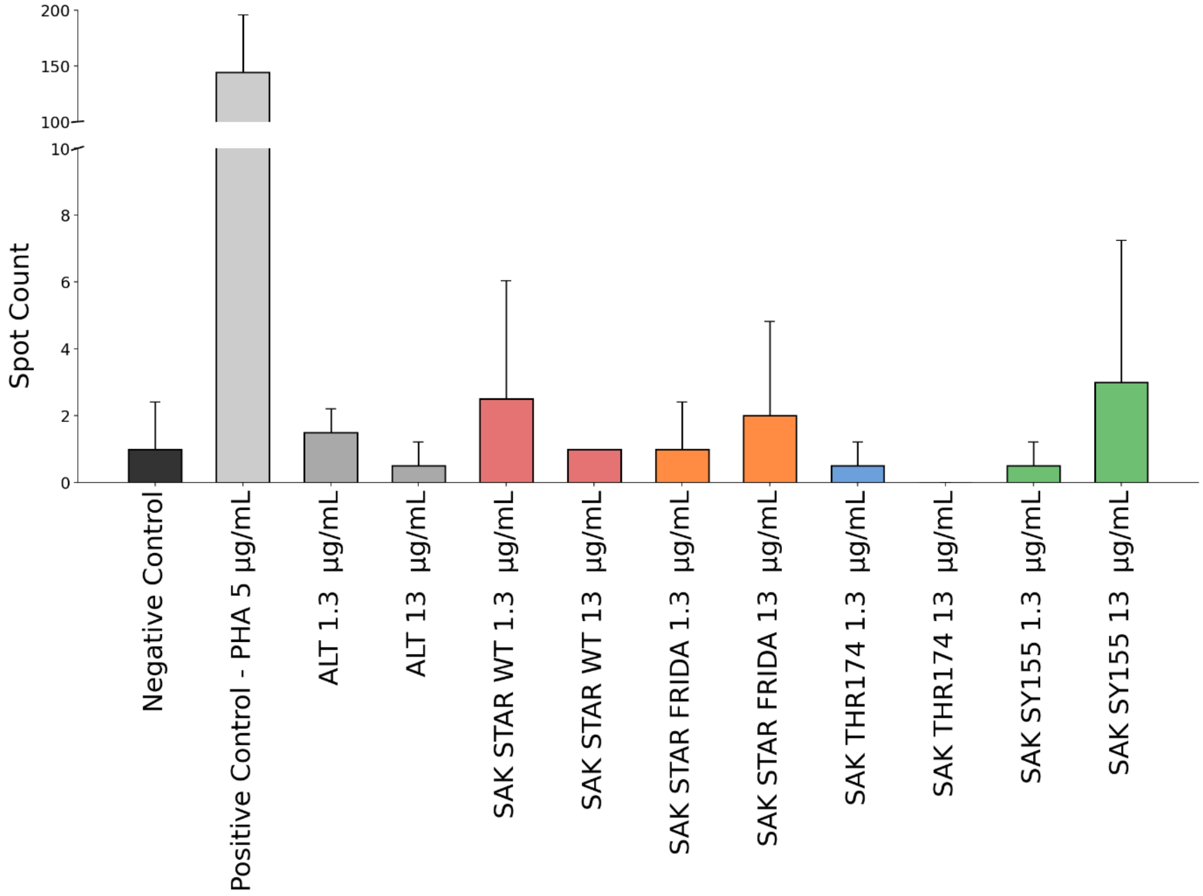


**Figure S17** FluoroSpot analysis of IFN-γ - CD8-depleted PBMCs were stimulated with medium alone (negative control), positive control (PHA, 5 µg/mL), or with ALT and SAK variants (1.3 and 13 µg/mL) for 3 days. IFN-γ secretion was quantified using fluorophore-labeled antibodies and analysed with an ImmunoSpot® reader (520 nm). Data are shown as mean spot counts ± standard deviation (n=2).

**Table S12** Monocyte activation test (MAT) statistics - Descriptive statistics and p-values for endotoxin removal confirmation, presented as IL-6 release. (A) Descriptive statistics for each condition (mean, SD = standard deviation, median, n = number of replicates). (B) Pairwise p-values from Kruskal-Wallis test with post-hoc Dunn's correction, with all conditions compared to the negative control and 0.007 EU/mL LPS (clinically acceptable limit). Statistical significance was set at p<0.05.

| **(A) MAT: Descriptive statistics** | | | | | | | | | | | | | |
|---|---|---|---|---|---|---|---|---|---|---|---|---|---|
| | **Negative control** | **LPS 0.007 EU/mL** | **LPS 0.07 EU/mL** | **ALT 1.3 µg/mL** | **ALT 13 µg/mL** | **SAK STAR WT 1.3 µg/mL** | **SAK STAR WT 13 µg/mL** | **SAK STAR FRIDA 1.3 µg/mL** | **SAK STAR FRIDA 13 µg/mL** | **SAK THR174 1.3 µg/mL** | **SAK THR174 13 µg/mL** | **SAK SY155 1.3 µg/mL** | **SAK SY155 13 µg/mL** |
| **Mean [pg/mL]** | 0.20 | 1.84 | 94.28 | 0.22 | 0.16 | 0.43 | 0.83 | 0.92 | 2.77 | 0.72 | 1.16 | 0.56 | 0.29 |
| **SD [pg/mL]** | 0.34 | 2.43 | 55.72 | 0.28 | 0.21 | 0.46 | 0.82 | 0.60 | 2.87 | 0.58 | 1.35 | 0.59 | 0.40 |
| **Median [pg/mL]** | 0.02 | 0.51 | 76.35 | 0.09 | 0.03 | 0.22 | 0.53 | 1.06 | 1.54 | 0.51 | 0.62 | 0.32 | 0.08 |
| **Count** | 10 | 11 | 11 | 10 | 9 | 9 | 11 | 6 | 5 | 4 | 6 | 8 | 6 |
| **(B) MAT: p-values** | | | | | | | | | | | | | |
| | **Negative control** | **LPS 0.007 EU/mL** | **LPS 0.07 EU/mL** | **ALT 1.3 µg/mL** | **ALT 13 µg/mL** | **SAK STAR WT 1.3 µg/mL** | **SAK STAR WT 13 µg/mL** | **SAK STAR FRIDA 1.3 µg/mL** | **SAK STAR FRIDA 13 µg/mL** | **SAK THR174 1.3 µg/mL** | **SAK THR174 13 µg/mL** | **SAK SY155 1.3 µg/mL** | **SAK SY155 13 µg/mL** |
| **Negative control** | - | 0.319 | <0.001 | >0.999 | >0.999 | >0.999 | 0.673 | 0.286 | 0.079 | >0.999 | 0.732 | >0.999 | >0.999 |
| **LPS 0.007 EU/mL** | - | 0.319 | 0.019 | 0.518 | 0.421 | >0.999 | >0.999 | >0.999 | >0.999 | >0.999 | >0.999 | >0.999 | >0.999 |

**Table S13** T lymphocyte proliferation statistics - Descriptive statistics and p-values for T lymphocyte proliferation, presented as percentage of proliferation. (A) Descriptive statistics for each condition (mean, SD = standard deviation, median, n = number of replicates). (B) Pairwise p-values from ordinary one-way ANOVA with Tukey's post hoc correction, with all conditions compared against each other. Statistical significance was set at p<0.05.

| **(A) T lymphocyte proliferation: Descriptive statistics** | | | | | | | | | | | | |
|---|---|---|---|---|---|---|---|---|---|---|---|---|
| | **Negative control** | **PHA 5 µg/mL** | **ALT 1.3 µg/mL** | **ALT 13 µg/mL** | **SAK STAR WT 1.3 µg/mL** | **SAK STAR WT 13 µg/mL** | **SAK STAR FRIDA 1.3 µg/mL** | **SAK STAR FRIDA 13 µg/mL** | **SAK THR174 1.3 µg/mL** | **SAK THR174 13 µg/mL** | **SAK SY155 1.3 µg/mL** | **SAK SY155 13 µg/mL** |
| **Mean [%]** | 1.19 | 95.20 | 0.96 | 0.61 | 1.23 | 2.83 | 1.12 | 1.18 | 0.99 | 1.53 | 0.80 | 1.95 |
| **SD [%]** | 1.46 | 4.98 | 1.01 | 0.89 | 1.59 | 5.19 | 1.57 | 1.34 | 1.21 | 2.64 | 0.97 | 3.64 |
| **Median [%]** | 0.71 | 97.40 | 0.89 | 0.19 | 0.65 | 0.34 | 0.52 | 1.09 | 0.67 | 0.28 | 0.52 | 0.20 |
| **Count** | 4 | 4 | 4 | 4 | 4 | 4 | 4 | 4 | 4 | 4 | 4 | 4 |

| **(B) T lymphocyte proliferation: p-values** | | | | | | | | | | | | |
|---|---|---|---|---|---|---|---|---|---|---|---|---|
| | **Negative control** | **PHA 5 µg/mL** | **ALT 1.3 µg/mL** | **ALT 13 µg/mL** | **SAK STAR WT 1.3 µg/mL** | **SAK STAR WT 13 µg/mL** | **SAK STAR FRIDA 1.3 µg/mL** | **SAK STAR FRIDA 13 µg/mL** | **SAK THR174 1.3 µg/mL** | **SAK THR174 13 µg/mL** | **SAK SY155 1.3 µg/mL** | **SAK SY155 13 µg/mL** |
| **Negative control** | - | <0.001 | >0.999 | >0.999 | >0.999 | 0.999 | >0.999 | >0.999 | >0.999 | >0.999 | >0.999 | >0.999 |
| **PHA 5 µg/mL** | - | - | <0.001 | <0.001 | <0.001 | <0.001 | <0.001 | <0.001 | <0.001 | <0.001 | <0.001 | <0.001 |
| **ALT 1.3 µg/mL** | - | - | - | >0.999 | >0.999 | 0.997 | >0.999 | >0.999 | >0.999 | >0.999 | >0.999 | >0.999 |
| **ALT 13 µg/mL** | - | - | - | - | >0.999 | 0.987 | >0.999 | >0.999 | >0.999 | >0.999 | >0.999 | >0.999 |
| **SAK STAR WT 1.3 µg/mL** | - | - | - | - | - | 0.999 | >0.999 | >0.999 | >0.999 | >0.999 | >0.999 | >0.999 |
| **SAK STAR WT 13 µg/mL** | - | - | - | - | - | - | 0.999 | 0.999 | 0.997 | >0.999 | 0.994 | >0.999 |
| **SAK STAR FRIDA 1.3 µg/mL** | - | - | - | - | - | - | - | >0.999 | >0.999 | >0.999 | >0.999 | >0.999 |
| **SAK STAR FRIDA 13 µg/mL** | - | - | - | - | - | - | - | - | >0.999 | >0.999 | >0.999 | >0.999 |
| **SAK THR174 1.3 µg/mL** | - | - | - | - | - | - | - | - | - | >0.999 | >0.999 | >0.999 |
| **SAK THR174 13 µg/mL** | - | - | - | - | - | - | - | - | - | - | >0.999 | >0.999 |
| **SAK SY155 1.3 µg/mL** | - | - | - | - | - | - | - | - | - | - | - | >0.999 |

**Table S14** Descriptive statistics for clot formation and lysis assay expressed as lysis time. The concentration of all agents was 2.5 nM. The experiment was performed in at 37 °C in PBS/$Ca^{2+}$/Tween80 buffer. **A)** Descriptive statistics for each group (mean, SD = standard deviation, median, n = number of replicates). **B)** Pairwise p-values from the Kruskal-Wallis test with post-hoc Dunn's correction. Statistical significance was set at p<0.05.

| **A) Clot formation and lysis assay: Descriptive statistics** | | | | | |
|---|---|---|---|---|---|
| | **SAK STAR WT** | **SAK 42D WT** | **SAK STAR FRIDA** | **SAK THR174** | **SAK SY155** |
| **Mean [min]** | 19.3 | 20.7 | 50.8 | 24.4 | 16.8 |
| **SD [min]** | 2.4 | 2.4 | 4.6 | 1.4 | 1.8 |
| **Median [min]** | 19.1 | 20.7 | 51.3 | 24.3 | 16.4 |
| **Count** | 11 | 11 | 11 | 11 | 11 |
| **B) Clot formation and lysis assay: p-values** | | | | | |
| | **SAK STAR WT** | **SAK 42D WT** | **SAK STAR FRIDA** | **SAK THR174** | **SAK SY155** |
| **SAK STAR WT** | - | 0.380 | <0.001 | 0.010 | 0.194 |
| **SAK 42D WT** | - | - | <0.001 | 0.079 | 0.042 |
| **SAK STAR FRIDA** | - | - | - | 0.088 | <0.001 |
| **SAK THR174** | - | - | - | - | <0.001 |

**Table S15** Determination of antibodies against individual SAK variants together with negative control (phosphate after vaccination of laboratory BALB/c mice. ELISA panels were coated with SAK STAR WT with a protein concentration of 5 µg/ml in carbonate buffer (100 mM bicarbonate/carbonate, pH 9.6) at 4 °C overnight. Sera of mice (dilution 1:500) immunized with individual SAK variants or with none (negative control) were applied to the wells of ELISA plates. The values are absorbance data measured at 450 nm.

| **Antibody determination using ELISA panels** | | | | | |
|---|---|---|---|---|---|
| | **SAK STAR WT** | **SAK STAR FRIDA** | **SAK THR174** | **SAK SY155** | **Negative control** |
| Mouse 1 | 0.222 | 0.164 | 0.248 | 0.172 | 0.216 |
| Mouse 2 | 0.153 | 0.208 | 0.478 | 0.162 | 0.199 |
| Mouse 3 | 0.140 | 0.306 | 0.187 | 0.169 | - |

**Table S16** Best-fit estimates kinetic and thermodynamic parameters determined via numerical integration of rate equations based on the model in illustrated in Scheme 1. Standard errors (S.E.) were calculated from the covariance matrix. Parameter confidence intervals (lower/upper limits) were rigorously defined by confidence contour analysis using a $\chi 2$ threshold of 0.98.

| **SPR affinity kinetic rates – standard error** | | | | | |
|---|---|---|---|---|---|
| | **SAK STAR WT** | **SAK 42D WT** | **SAK STAR FRIDA** | **SAK THR174** | **SAK SY155** |
| $k_1$ [nM$^{-1}$*s$^{-1}$] (*10$^{-3}$) | 0.58 ± 0.07 | 1.3 ± 0.2 | 0.55 ± 0.1 | 1.06 ± 0.62 | 0.90 ± 0.04 |
| $k_{-1}$ [s$^{-1}$] (*10$^{3}$) | 82 ± 7 | 88 ± 3 | 162 ± 18 | 93 ± 7 | 26 ± 2 |
| $k_2$ [nM$^{-1}$*s$^{-1}$] (*10$^{3}$) | 36 ± 10 | 51 ± 2 | 34 ± 3 | 20 ± 7 | 37 ± 1 |
| $k_{-2}$ [s$^{-1}$] (*10$^{3}$) | 73 ± 25 | 65 ± 7 | 28 ± 8 | 62 ± 33 | 10 ±1 |
| $k_3$ [nM$^{-1}$*s$^{-1}$] (*10$^{3}$) | 0.58 ± 0.02 | 0.57 ± 0.02 | 0.39 ± 0.06 | 0.55 ± 0.05 | 0.42 ± 0.05 |
| $k_{-3}$ [s$^{-1}$] (*10$^{3}$) | 8.8 ± 0.6 | 7.1 ± 0.4 | 75 ± 13 | 12 ± 1 | 20 ± 3 |
| **SPR affinity kinetic rates – confidence intervals** | | | | | |
| | **SAK STAR WT** | **SAK 42D WT** | **SAK STAR FRIDA** | **SAK THR174** | **SAK SY155** |
| $k_1$ [nM$^{-1}$*s$^{-1}$] (*10$^{-3}$) | 0.54 – 0.81 | 1.0 – 1.4 | 0.51 – 0.91 | 1.06 – 1.13 | 0.68 – 0.95 |
| $k_{-1}$ [s$^{-1}$] (*10$^{3}$) | 82 – 93 | 56 – 97 | 130 – 253 | 93 – 93 | 13 – 46 |
| $k_2$ [nM$^{-1}$*s$^{-1}$] (*10$^{3}$) | 15 – 45 | 41 – 60 | 28 – 54 | 10 – 32 | 33 – 96 |
| $k_{-2}$ [s$^{-1}$] (*10$^{3}$) | 69 – 177 | 62 – 67 | 25 – 34 | 62 – 134 | 8 – 16 |
| $k_3$ [nM$^{-1}$*s$^{-1}$] (*10$^{3}$) | 0.53 – 0.59 | 0.51 – 0.62 | 0.35 – 0.45 | 0.51 – 0.56 | 0.34 – 0.63 |
| $k_{-3}$ [s$^{-1}$] (*10$^{3}$) | 8.5 – 9.0 | 6.8 – 7.4 | 6.6 – 8.3 | 12 - 13 | 16 – 32 |
| **SPR affinity scaling parameters – standard error** | | | | | |
| | **SAK STAR WT** | **SAK 42D WT** | **SAK STAR FRIDA** | **SAK THR174** | **SAK SY155** |
| $a_1$ | 4.25 ± 0.09 | 3.96 ± 0.07 | 2.34 ± 0.70 | 4.20 ± 0.13 | 5.38 ± 0.07 |
| $a_2$ | 4.87 ± 0.10 | 3.65 ± 0.07 | 2.11 ± 0.63 | 3.77 ± 0.12 | 4.88 ± 0.07 |
| $a_3$ | 3.62 ± 0.08 | 3.13 ± 0.06 | 1.98 ± 0.57 | 3.39 ± 0.11 | 4.98 ± 0.07 |
| $a_4$ | 3.68 ± 0.08 | 3.35 ± 0.06 | 1.83 ± 0.55 | 3.42 ± 0.11 | 6.44 ± 0.09 |
| $a_5$ | 3.20 ± 0.07 | 2.42 ± 0.05 | 1.72 ± 0.52 | 2.61 ± 0.08 | 3.89 ± 0.05 |
| $a_6$ | 2.20 ± 0.05 | 2.45 ± 0.05 | 1.57 ± 0.47 | 2.63 ± 0.08 | 3.38 ± 0.05 |
| b | 1.26 ± 0.11 | 2.98 ± 0.12 | 3.53 ± 1.07 | 2.23 ± 0.32 | 1.71 ± 0.02 |
| c | 2.15 ± 0.05 | 2.38 ± 0.04 | 3.87 ± 1.56 | 2.10 ± 0.07 | 1.31 ± 0.03 |
| d (*10$^{-3}$) | 1.5 ± 0.1 | 1.6 ± 0.1 | 0.79 ± 0.01 | 0.91 ± 0.01 | 2.0 ± 0.1 |
| **SPR affinity scaling parameters – confidence intervals** | | | | | |
| | **SAK STAR WT** | **SAK 42D WT** | **SAK STAR FRIDA** | **SAK THR174** | **SAK SY155** |
| $a_1$ | 4.25 – 4.27 | 3.95 – 3.97 | 2.33 – 2.34 | 4.20 – 4.20 | 5.34 – 5.41 |
| $a_2$ | 4.87 – 4.90 | 3.64 – 3.65 | 2.10 – 2.11 | 3.77 – 3.77 | 4.84 – 4.90 |
| $a_3$ | 3.62 – 3.63 | 3.13 – 3.14 | 1.88 – 1.98 | 3.39 – 3.39 | 4.93 – 5.00 |
| $a_4$ | 3.86 – 3.88 | 3.34 – 3.35 | 1.82 – 1.83 | 3.42 – 3.42 | 6.36 – 6.46 |
| $a_5$ | 3.20 – 3.22 | 2.42 – 2.43 | 1.71 -1.72 | 2.61 – 2.61 | 3.85 – 3.90 |
| $a_6$ | 2.20 – 2.20 | 2.44 – 2.45 | 1.57 – 1.57 | 2.63 – 2.63 | 3.35 – 3.39 |
| b | 1.26 – 1.29 | 2.97 – 2.99 | 3.50 – 3.54 | 2.23 – 2.23 | 1.70 – 1.83 |
| c | 2.15 – 2.21 | 2.37 – 2.40 | 3.85 – 3.87 | 2.09 – 2.11 | 1.16 – 1.32 |
| d (*10$^{-3}$) | 1.4 – 1.6 | 1.5 – 1.7 | 0.63 – 1.0 | 0.84 – 0.92 | 1.9 – 2.1 |

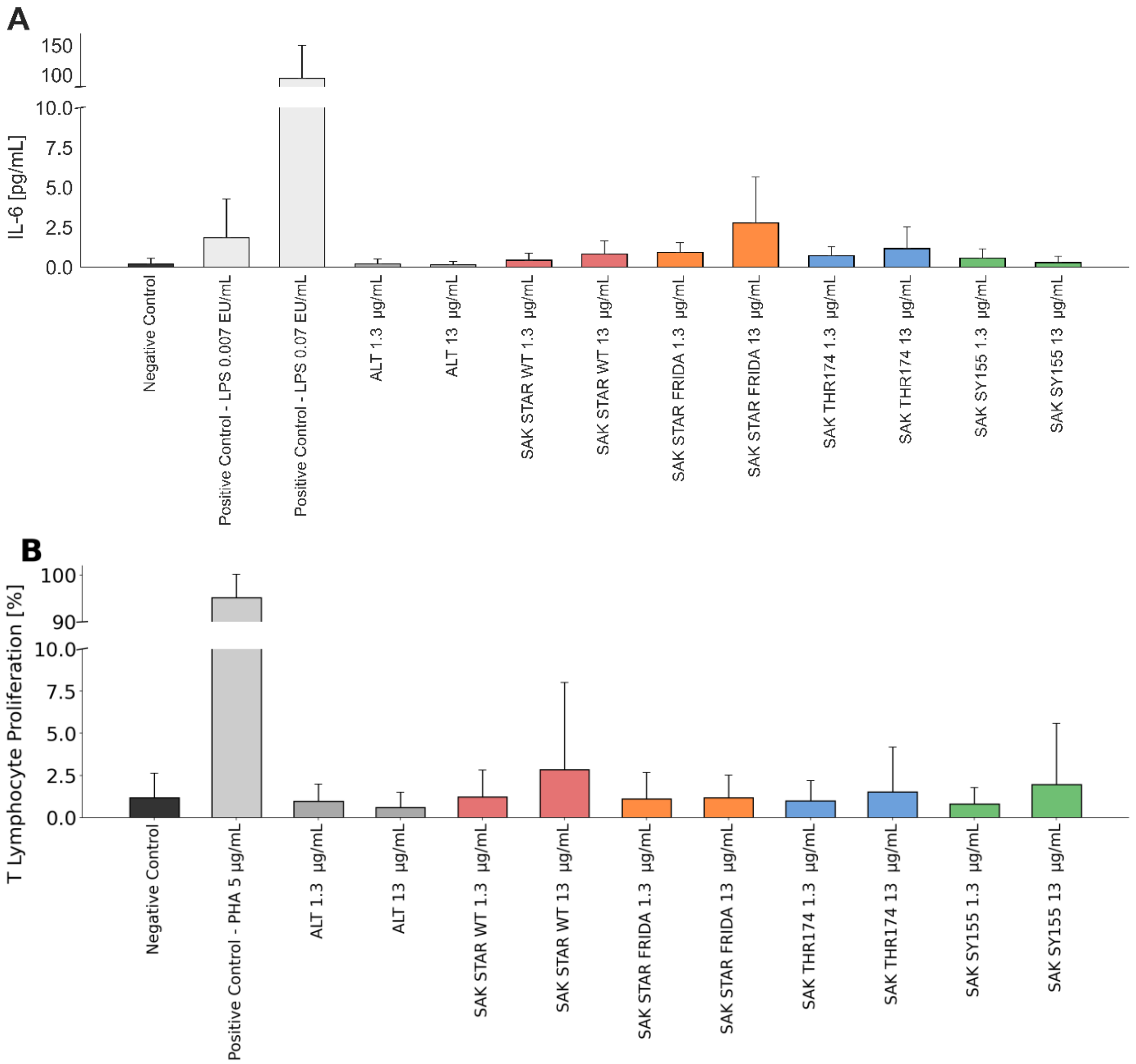


**Figure S18** Immunological characterization of SAK variants using Monocyte activation test (MAT) and T lymphocyte proliferation test. A) Monocyte activation test (MAT). IL-6 release was measured in 10-fold diluted human blood stimulated with negative control, LPS at 0.007 EU/mL (clinically acceptable limit), and LPS at 0.07 EU/mL (10-fold higher, assay functionality control), as well as with ALT and SAK variants at 1.3 and 13 µg/mL. Data are shown as mean ± standard deviation from 4-11 independent donors. Kruskal-Wallis test with Dunn's post hoc correction was performed against negative control and 0.007 EU/mL LPS. B) T lymphocyte proliferation test. Proliferation of CD8-depleted PBMCs from four healthy donors was assessed following stimulation with negative control, positive control (PHA 5 µg/mL), ALT (1.3 and 13 µg/mL), or SAK variants (1.3 and 13 µg/mL). Data are shown as mean ± standard deviation. One-way ANOVA with Tukey's post hoc correction was performed between all conditions.

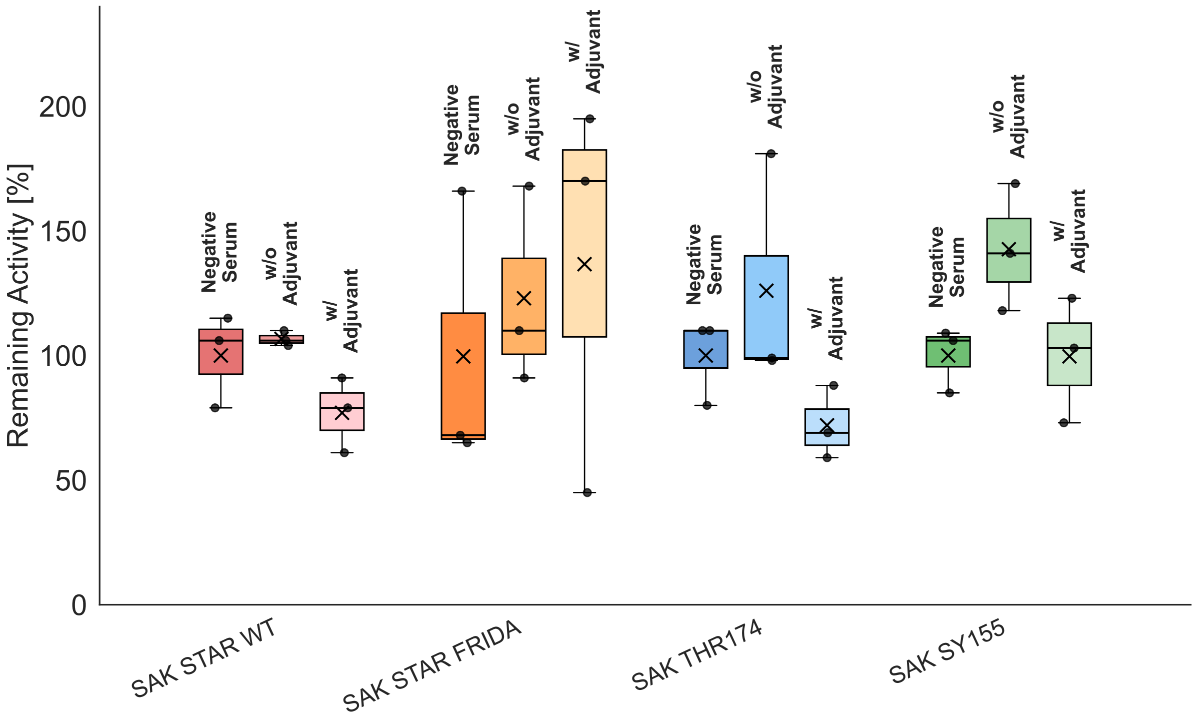


**Figure S19** Comparison of serum inhibitory potency after administration of four SAK variants under different conditions: negative control, administration without adjuvant, and administration with adjuvant. Boxplots represent the distribution of three biological replicates for each condition, with whiskers indicating the full data range (min–max). Individual replicate values are shown as black dots, while black crosses indicate mean values. For each protein variant, condition-specific measurements are displayed using different tonal shades of the same base color: dark shades represent negative serum controls, medium shades represent samples without adjuvant, and light shades represent samples with adjuvant.

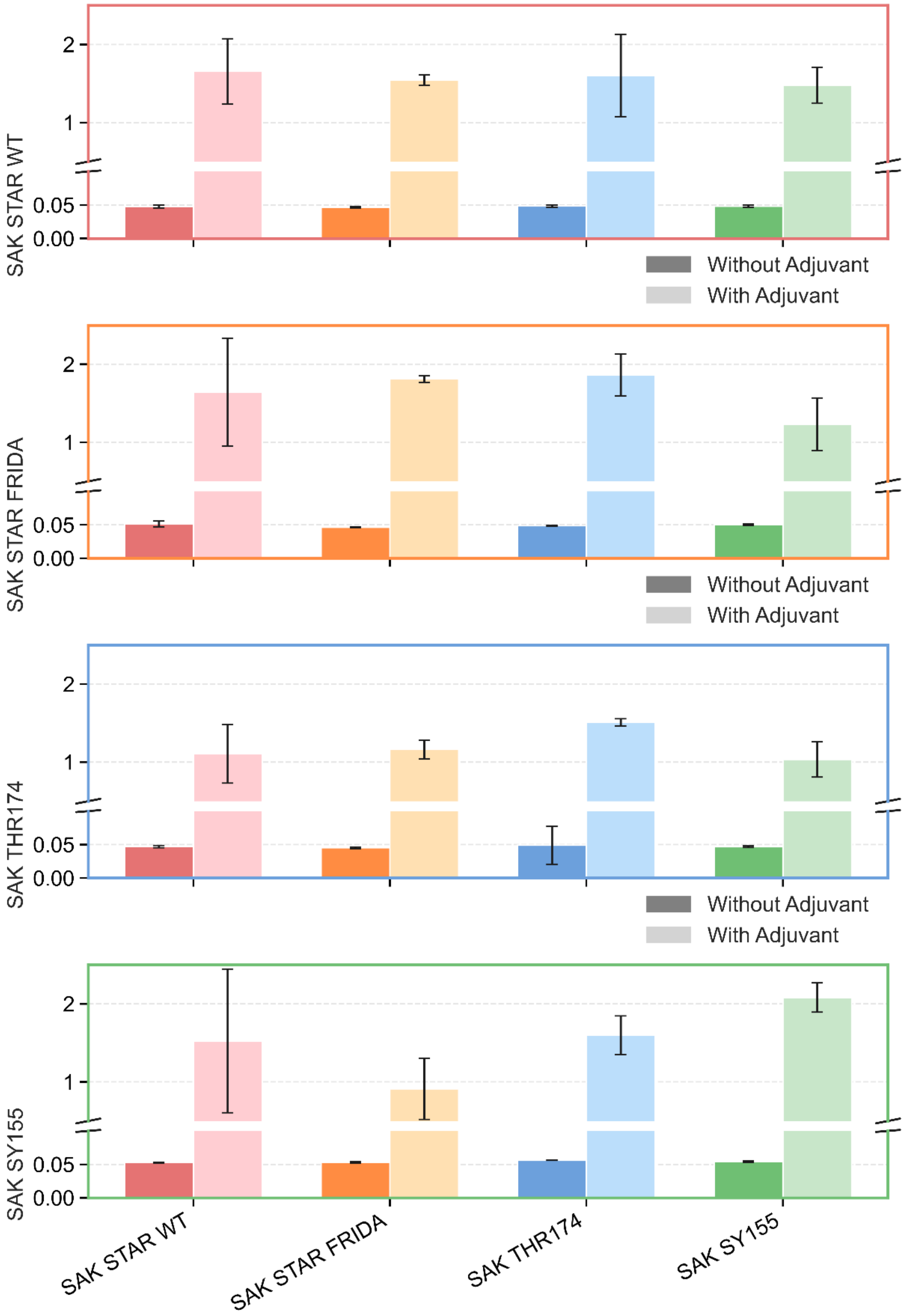


**Figure S20** Determination of antibodies against individual SAK variants after vaccination of laboratory BALB/c mice. ELISA panels were coated with respective variants of SAK and sera of mice (dilution 1:500) immunized with individual SAK variants without (darker color bars) and with (lighter color bars) aluminum adjuvant were applied to respective wells of ELISA plates. The SAK variant used for coating is indicated on the left side of the individual subplots. The values are absorbance data measured at 450 nm. Data are shown as mean ± standard deviation from two independent ELISA replicates.

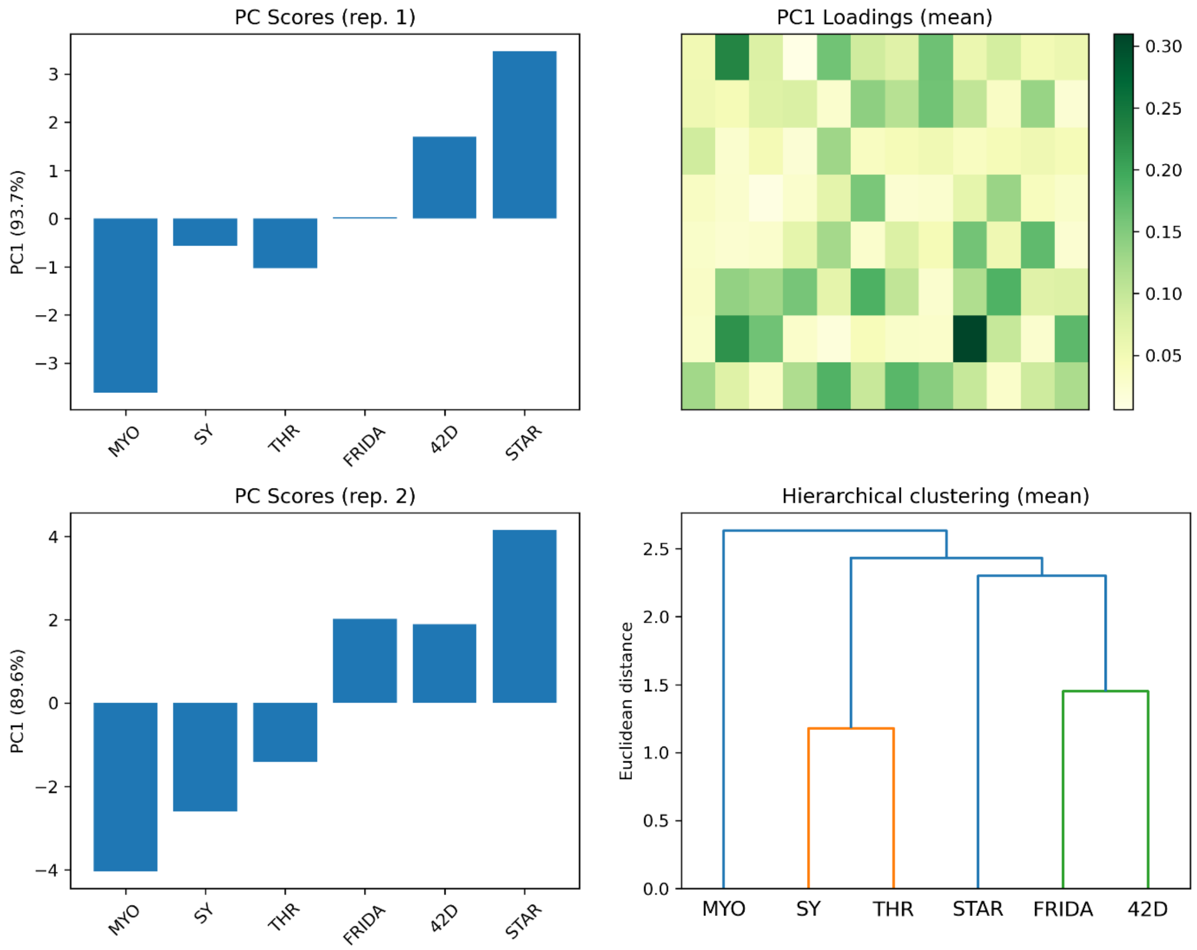


**Figure S21** Data analysis of reactivity of human sera for individual SAK variants. Left: the first principal component of the centered non-normalized absorbance values of human sera reactivity for each independent replica (left); Top right: the loadings graph for the first principal component of the mean values of absorbances; Bottom right: hierarchical clustering of the SAK variants and the mock protein (Myo) for mean values of absorbances.
